# Evolution of neural activity in circuits bridging sensory and abstract knowledge

**DOI:** 10.1101/2022.01.29.478317

**Authors:** Francesca Mastrogiuseppe, Naoki Hiratani, Peter Latham

## Abstract

The ability to associate sensory stimuli with abstract classes is critical for survival. How are these associations implemented in brain circuits? And what governs how neural activity evolves during abstract knowledge acquisition? To investigate these questions, we consider a circuit model that learns to map sensory input to abstract classes via gradient descent synaptic plasticity. We focus on typical neuroscience tasks (simple, and context-dependent, categorization), and study how both synaptic connectivity and neural activity evolve during learning. To make contact with the current generation of experiments, we analyze activity via standard measures such as selectivity, correlations, and tuning symmetry. We find that the model is able to recapitulate experimental observations, including seemingly disparate ones. We determine how, in the model, the behaviour of these measures depends on details of the circuit and the task. These dependencies make experimentally-testable predictions about the circuitry supporting abstract knowledge acquisition in the brain.

## Introduction

Everyday decisions do not depend on the state of the world alone; they also depend on internal, non-sensory variables that are acquired with experience. For instance, over time we learn that in most situations salads are good for us while burgers are not, while in other contexts (for example, before a long hike in the mountains) the opposite is true. The ability to associate sensory stimuli with abstract variables is critical for survival; how these associations are learned is, however, poorly understood.

Although we do not know how associations are learned, we do have access to a large number of experimental studies addressing how neural activity evolves while animals learn to classify stimuli into abstract categories [1–5]. Such experiments have probed two kinds of associations between stimuli and categories: fixed associations [4,6,7] (in which, for example, stimuli are either in category `A` or in category `B`), and flexible ones [5, 8–10] (in which, for example, stimuli are in category `A` in one context and category `B` in another).

A consistent finding in these experiments is that activity of single neurons in associative cortex develops selectivity to task-relevant abstract variables, such as category [3, 5, 6] and context [8, 9, 11]. Neurons, however, typically display selectivity to multiple abstract variables [12], and those patterns of mixed selectivity are often hard to intepret [7, 10, 13].

Instead of focusing on one neuron at the time, one can alternatively consider large populations of neurons and quantify how those, as a whole, encode abstract variables. This approach has led, so far, to apparently disparate observations. Classical work indicates that neurons in visual cortex encode simple sensory variables (for example, two opposite orientations) via negatively-correlated responses [14, 15]: neurons that respond strongly to a given variable respond weakly to the other one, and vice versa. Those responses, furthermore, are symmetric [16]: about the same number of neurons respond strongly to one variable, or the other. In analogy with sensory cortex, one can thus hypothesize that neurons in associative cortex encode different abstract variables (for example, category `A` and `B`) via negatively-correlated, and symmetric responses. Evidence in favour of this type of responses has been reported in monkeys [7, 10, 11, 17] and mice [5] prefrontal cortex (PFC). However, evidence in favour of a different type of responses has been reported in a different set of experiments from monkeys lateral intraparietal (LIP) cortex [18]. In that case, resposes to category `A` and `B` were found to be positively correlated: neurons that learn to respond strongly to category `A` also respond strongly to category `B`, and neurons that learn to respond weakly to `A` also respond weakly to `B`. Furthermore, responses were strongly asymmetric: almost all neurons displayed the strongest response to the same category (despite monkeys did not display behavioural biases towards one category or the other).

In this work, we use neural circuit models to shed light on these experimental results. To this end, we hypothesize that synaptic connectivity in neural circuits evolves by implementing gradient descent on an error function [19]. A large body of work has demonstrated that, under gradient-descent plasticity, neural networks can achieve high performance on both simple and complex tasks [20]. Recent studies have furthermore shown that gradient-descent learning can be implemented, at least approximately, in a biologically plausible way [21–27]. Concomitantly, gradient-based learning has been used to construct network models for a variety of brain regions and functions [28–31]. A precise understanding of how gradient descent learning shapes representations in neural circuits is however still lacking.

Motivated by this hypothesis, we study a minimal circuit model that learns through gradient descent to associate sensory stimuli with abstract categories, with a focus on tasks inspired by those used in experimental studies. Via mathematical analysis and simulations, we show that the model can capture the experimental findings discussed above. In particular, after learning, neurons in the model become selective to category and, if present, context; this result is robust, and independent of the details of the circuit and the task. On the other hand, whether correlations after learning are positive or negative, and whether population tuning to different categories is asymmetric or not, is not uniquely determined, but depends on details. We determined how, in the model, activity measures are modulated by circuit details (activation function of single neurons, learning rates, initial connectivity) and task features (number of stimuli, and whether or not the associations are context-dependent). These dependencies make experimentally testable predictions about the underlying circuitry. Overall, the model provides a framework for interpreting seemingly disparate experimental findings, and for making novel experimental predictions.

## Results

We consider classification into mutually exclusive abstract classes which, as above, we refer to as category `A` and category `B`. We consider two tasks: a simple, linearly-separable one [4, 6, 7] and a context-dependent, non linearly-separable, one [5, 8, 10] (Fig. 1A). We assume that for both, categorization is implemented with a two-layer circuit, as shown in Fig. 1B, and that the synaptic weights evolve via gradient descent. Our goal is to determine how the activity in the intermediate layer evolves with learning, and how this evolution depends on the task and the biophysical details of the circuit. We start by describing the model. We then consider circuits that learn the simple, linearly-separable, categorization task, and analyze how learning drives changes in activity. Finally, we extend the analysis to the context-dependent, non linearly-separable, task.

**Fig. 1:**
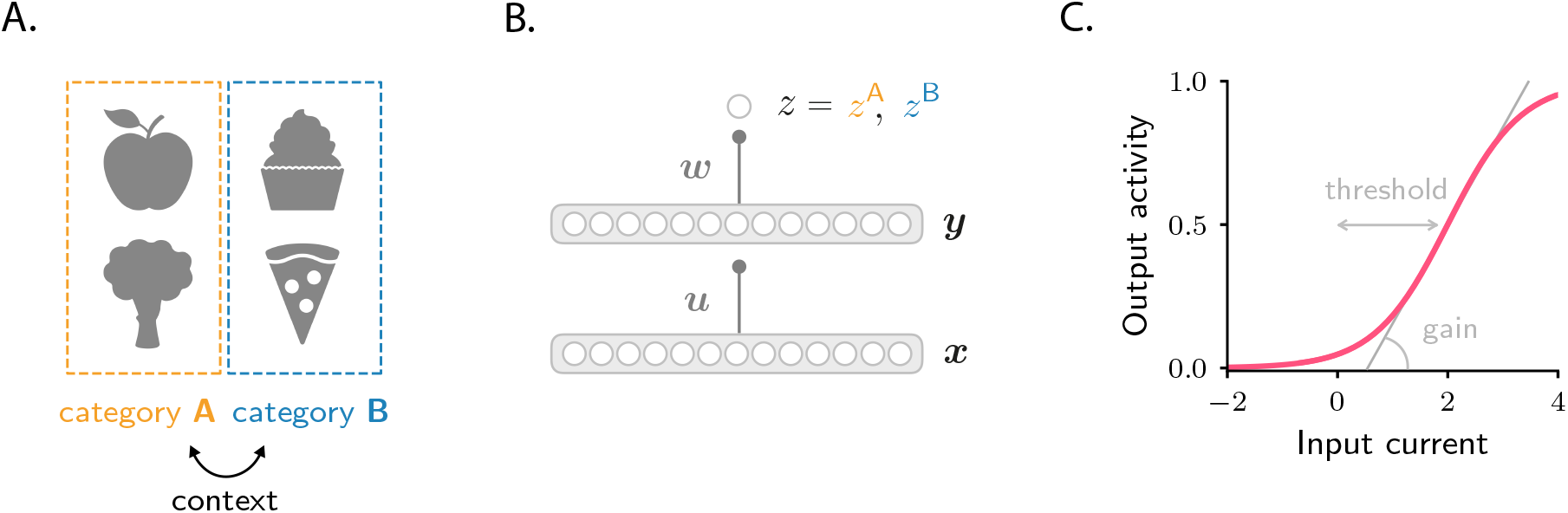
Schematics of tasks and circuit model used in the study. **A.** Illustration of the two categorization tasks. In the simple categorization task, half the stimuli are associated with category `A` and the other half with category `B`. In the context-dependent task, associations are reversed across contexts: stimuli associated with category `A` in context `1` are associated with category `B` in context `2`, and vice versa. **B.** The circuit consists of a sensory input layer, ***x***, an intermediate layer, **y**, and a readout neuron, *z*. The intermediate (***u***) and readout (***w***) weights evolve under gradient descent plasticity (Eq. 1). **C.** The activation functions, **Ψ** and **Φ**, are taken to be sigmoids characterized by a threshold and a gain. The gain, which controls the sensitivity of activity to input, is the slope of the function at its steepest point; the threshold, which controls activity sparsity, is the distance from the steepest point to zero.

### Circuit model

We consider a simple feedforward circuit as in Fig. 1B. A vector ***x***, which models the input from sensory areas, is fed into an intermediate layer of neurons which represents a higher-level, associative area. The intermediate layer activity is given by ***y*** = **Ψ**(***u*** · ***x***), where ***u*** is a all-to-all connectivity matrix. That activity projects to a readout neuron, which learns, over time, to predict the category associated with each sensory input. The activity of the readout neuron, *z*, is taken to be *z* = **Φ**(***w*** · ***y***), where ***w*** is a readout vector. The activation functions **Φ** and **Ψ** are sigmoidals that encapsulate the response properties of single neurons; they are parametrized by a threshold and a gain (Fig. 1C; Methods 2.1).

The goal of the circuit is to adjust the synaptic weights, ***u*** and ***w***, so that the readout neuron fires at rate *z* = *z*^A^ when the sensory input is associated with category `A`, and at rate *z* = *z*^B^ when the sensory input is associated with `B` (Fig. 1B). In the simple categorization task, half the stimuli are associated with category `A` and the other half with `B`. In the context-dependent task, associations are reversed across contexts: stimuli associated with category `A` in context `1` are associated with category `B` in context `2`, and vice versa (Fig. 1A). We use 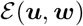 to denote the average error between *z* and its target value, and assume that the synaptic weights evolve, via gradient descent, to minimize the error. If the learning rates are small, the weights evolve according to

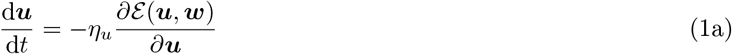

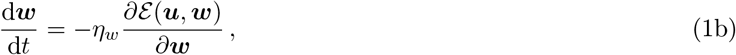

where *t* represents learning time and *η_u_* and *η_w_* are learning rates which, for generality, we allow to be different.

Before learning, the synaptic weights are random. Consequently, activity in the intermediate layer, ***y***, is unrelated to category, and depends only on sensory input. As the circuit learns to associate sensory inputs with abstract categories, task-relevant structure emerges in the connectivity matrix ***u***, and thus in the intermediate layer as well. Analyzing how activity in the intermediate layer evolves over learning is the focus of this work.

### Evolution of activity during the simple categorization task

We first analyze the simple task, for which we can derive results in a transparent and intuitive form. We then go on to show that similar (although richer) results hold for the context-dependent one.

In the simple categorization task, each sensory input vector ***x***^s^ represents a stimulus (for example, an odor, or an image), which is associated with one of the two mutually-exclusive categories `A` and `B`. In the example shown below, we used 20 stimuli, of which half are associated with category `A`, and the other half are associated with category `B`. Sensory input vectors corresponding to different stimuli are generated at random and assumed to be orthogonal to each other; orthogonality is motivated by the decorrelation performed by sensory areas (but this assumption can be dropped without qualitatively changing the main results, see Methods 4.8 and Fig. S6).

We start our analysis by simulating the circuit numerically, and investigating the properties of neural activity, ***y***, in the intermediate layer. A common way to characterize the effects of learning on single-neuron activity is through the category selectivity index, a quantity that is positive when activity elicited by within-category stimuli is more similar than activity elicited by across-category stimuli, and negative otherwise. It is defined as [3–5] (Methods 4.3)

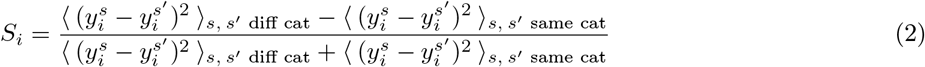

where 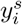 represents the activity of neuron *i* in response to sensory input *s*, and angle brackets, 〈·〉_s,s′_, denote an average over sensory input pairs. The subscript “same cat” refers to averages over the same category (`A-A` or `B-B`) and “diff cat” to averages over different categories (`A-B`).

Before learning, the responses of single neurons to different stimuli are random and unstructured. Thus, responses to stimuli paired with category `A` are statistically indistinguishable from responses to stimuli paired with category `B` (Fig. 2A). This makes the category selectivity index zero on average (Fig. 2B). After learning, the responses of single neurons depend on category: within-category responses become more similar than across-category responses, resulting in two separate distributions (Fig. 2E). As a consequence, the category selectivity index for most neuron increases; correspondingly, average selectivity increases from zero to positive values (Fig. 2F), thus reproducing the behaviour observed in experimental studies [3–5]. To determine whether this effect is robust, we varied the parameters that describe the task (number of stimuli) and the biophysical properties of the circuit (the threshold and gain of neurons, Fig. 1C, and the learning rates of the two sets of synaptic weights, *η_u_* and *η_w_*). We found that the selectivity increase is a universal property – it is observed in all circuit models that successfully learned the task, independent of the parameters. Activity from a second example circuit is shown in Figs. 2I-J; additional simulations are shown in Fig. S1A.

**Fig. 2:**
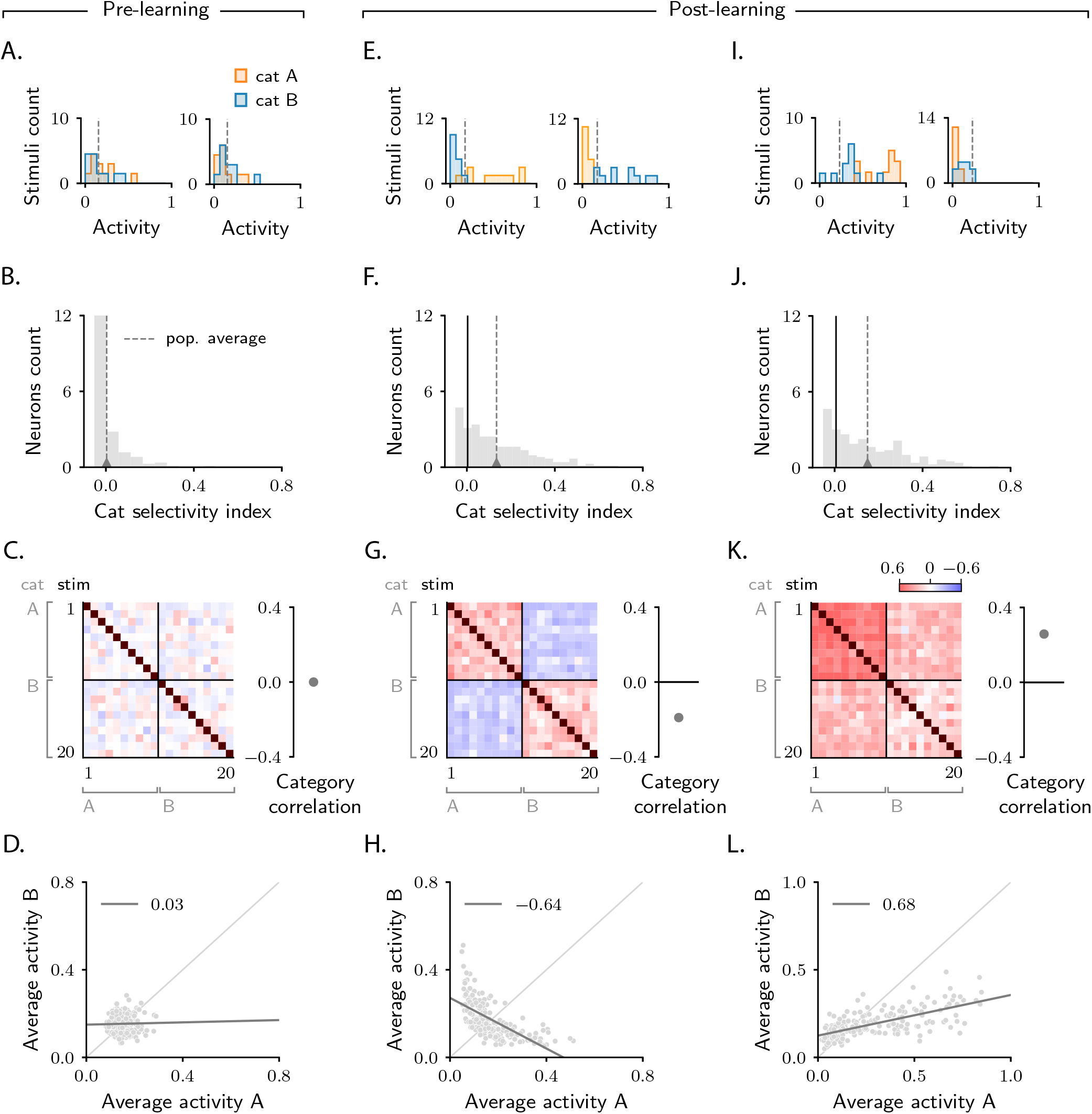
Characterization of activity evolution during the simple categorization task. Results from simulations. The first column (A-D) shows a naive circuit (pre-learning); the second (E-H) and third (I-L) columns show two trained circuits (post-learning), characterized by different sets of parameters (see below). **A, E, I.** Histograms of single-neuron activity in response to stimuli associated with category `A` (orange) and category `B` (blue). Left and right show two sample neurons from the intermediate layer. Grey dashed lines indicate the average activity across the population. **B, F, J.** Histograms of category selectivity (Eq. 2) across the population of neurons in the intermediate layer. Grey dashed lines indicate the average selectivity across the population. In panels F and J, the black vertical lines indicate the initial value of average selectivity. **C, G, K.** Signal correlation matrices. Each entry shows the Pearson correlation coefficient, averaged over neurons (Eq. 72), between activity elicited by different stimuli. In these examples, we used 20 stimuli. Diagonal entries (brown) are all equal to 1. Category correlation (namely, the average of the correlations within the off-diagonal blocks, which contain stimuli in different categories) is shown on the right of the matrices. In panels G and K, the black horizontal lines near zero indicate the initial values of category correlation. **D, H, L.** Population responses to categories `A` and `B`. Each dot represents a neuron in the intermediate layer, with horizontal and vertical axes showing the responses to stimuli associated with categories `A` and `B`, respectively, averaged over stimuli. Grey line: linear fit, with Pearson correlation coefficient shown in the figure legend. Parameters are summarized in Table 1 (Methods 6.2).

Category selectivity tells us about the behavior of single neurons. But how does the population as a whole change its activity over learning? To quantify that, we compute signal correlations, defined to be the Pearson correlation coefficient between the activity elicited by two different stimuli [7]. Results are summarized in the correlation matrices displayed in Figs. 2C, G and K. As the task involves 20 stimuli, the correlation matrix is 20 × 20; stimuli are sorted according to category.

As discussed above, before learning the responses of neurons in the intermediate layer are random and unstructured. Thus, activity in response to different stimuli is uncorrelated; this is illustrated in Fig. 2C, where all non-diagonal entries of the correlation matrix are close to zero. Of particular interest are the upper-right and lower-left blocks of the matrix, which correspond to pairs of activity vectors elicited by stimuli in different categories. The average of those correlations, which we refer to as category correlation, is shown to the right of each correlation matrix. Before learning, the category correlation is close to zero (Fig. 2C). Over learning, the correlation matrices develop structure. Correlations become different within the two diagonal, and the two off-diagonal blocks, indicating that learning induces category-dependent structure. In Fig. 2G, the average correlation within the off-diagonal blocks is negative; the category correlation is thus negative [7, 10, 17]. The model does not, however, always produce negative correlation: varying model details – either the parameters of the circuit or the number of stimuli – can switch the category correlation from negative to positive [18]; one example is shown in Fig. 2K.

To illustrate the difference in population response when category correlation is negative versus positive, for each neuron in the intermediate layer we plot the average response to stimuli associated with category `B` (vertical axis) versus `A` (horizontal axis). Before learning, activity is unstructured, and the dots form a random, uncorrelated cloud (Fig. 2D). After learning, the shape of this cloud depends on category correlation. In Fig. 2H, where the category correlation is negative, the cloud has a negative slope. This is because changes in single-neuron responses to categories `A` and `B` have opposite sign: a neuron that increases its activity in response to category `A` decreases its activity in response to category `B` (Fig. 2E left), and vice versa (Fig. 2E right). In Fig. 2L, where the category correlation is positive, the cloud has, instead, a positive slope. Here, changes in single-neuron responses to categories `A` and `B` have the same sign: a neuron that increases its activity in response to category `A` also increases its activity in response to category `B` (Fig. 2I left), and similarly for a decrease (Fig. 2I right).

Negative versus positive slope is not the only difference between Figs. 2H and L: they also differ in symmetry with respect to the two categories. In Fig. 2H, about the same number of neurons respond more strongly to category `A` than to category `B` [5]. In Fig. 2L, however, the number of neurons that respond more strongly to category `A` is significantly larger than the number of neurons that respond more strongly to `B` [18]. Furthermore, as observed in experiments reporting positive correlations [18], the mean population activity in response to category `A` is larger than to category `B`, and the range of activity in response to `A` is larger than to `B`. The fact that the population response to `A` is larger than to `B` is not a trivial consequence of having set a larger target for the readout neuron in response to `A` than to `B` (*z*^A^ > *z*^B^): as shown in Figs. S2B and D, example circuits displaying larger responses to `B` can also be observed. Response asymmetry is discussed in detail in Methods 4.6.

In sum, we simulated activity in circuit models that learn to associate sensory stimuli to abstract categories via gradient descent synaptic plasticity. We observed that single neurons consistently develop selectivity to abstract categories – a behaviour that is robust with respect to model details. How the population of neurons responds to category depended, however, on model details: we observed both negatively correlated, symmetric responses and positively correlated, asymmetric ones. These observations are in agreement with experimental findings [4, 5, 7, 18].

### Analysis of the simple categorization task

What are the mechanisms that drive activity changes over learning? And how do the circuit and task details determine how the population responds? To address these questions, we performed mathematical analysis of the model. Our analysis is based on the assumption that the number of neurons in each layer of the circuit is much larger than the number of sensory inputs to classify – a regime that is relevant to the systems and tasks we study here. In that regime, the number of synaptic weights that the circuit can tune is very large, and so a small change in each weight is sufficient to learn the task. This makes the circuit amenable to mathematical analysis [32–35]; full details are reported in Methods 3, 4 and 5, here we illustrate the main results.

We start with the simple categorization task illustrated in the previous section, and use the mathematical framework to shed light on the simulations described above (Fig. 2). Figure 3A shows, schematically, activity in the intermediate layer before learning (see Fig. S1B for simulated data). Axes on each plot correspond to activity of three sample neurons. Each dot represents activity in response to a different sensory input; orange and blue dots indicate activity in response to stimuli associated with categories `A` and `B`, respectively. Before learning, activity is determined solely by sensory inputs, which consist of random, orthogonal vectors. Consequently, the initial activity vectors form an unstructured cloud in activity space, with orange and blue circles intermingled (Fig. 3A).

**Fig. 3:**
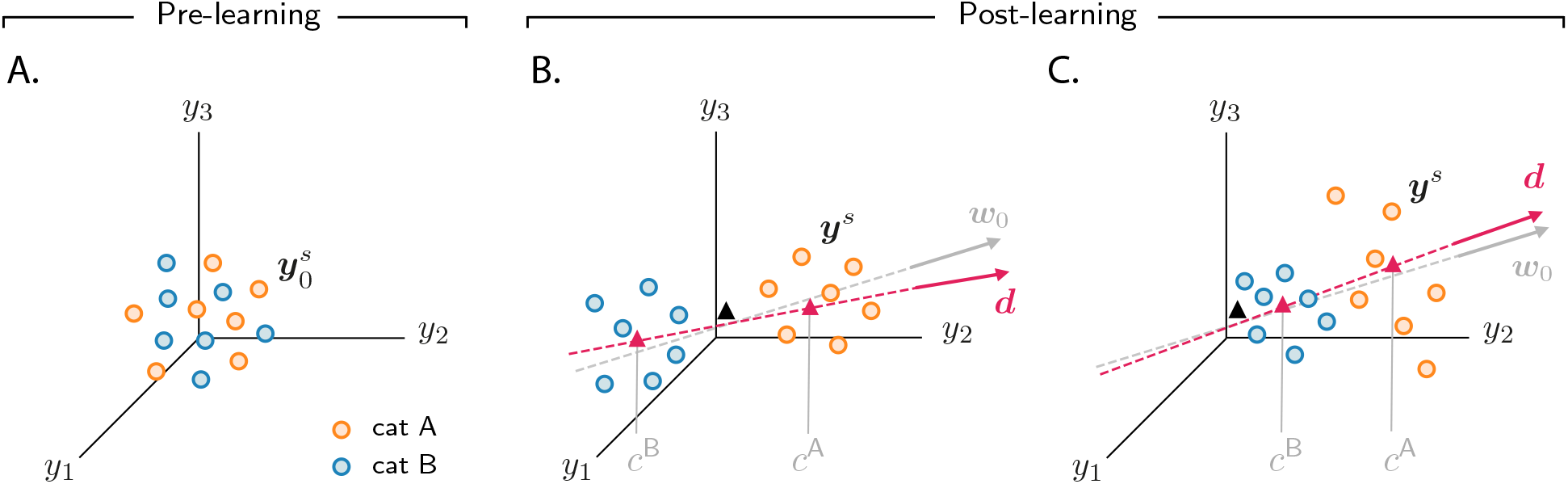
Analysis of activity evolution during the simple categorization task. Results from mathematical analysis. **A, B, C.** Cartoons illustrating how activity evolves over learning. The three columns are as in Fig. 2: pre-learning (first column) and post-learning for two different circuits (second and third columns). Circles show activity in the intermediate layer in response to different stimuli, displayed in a three-dimensional space where axes correspond to the activity of three sample neurons. Orange and blue circles are associated, respectively, with category A and B. Before learning, activity is unstructured (panel A). After learning (panels B and C), the activity vectors develop a component along the common direction ***d*** (Eq. 3), shown as a magenta line, and form two clouds, one for each category. The centers of those clouds are indicated by magenta triangles; their positions along ***d*** are given, approximately, by *c*^A^ and *c*^B^. The black triangle indicates the center of initial activity. In panel B, *c*^A^ and *c*^B^ have opposite sign, so the clouds move in opposite directions with respect to initial activity; in panel C, *c*^A^ and *c*^B^ have the same sign, so the clouds move in the same direction. For illustration purposes, we show a smaller number of stimuli (14, instead of 20) than in Fig. 2. Simulated data from the circuits displayed in Fig. 2 are shown in Fig. S1B.

Over learning, activity vectors in Fig. 3A move. Specifically, over learning all activity vectors acquire a component thatis aligned with a common, stimulus-independent direction. Activity after learning can thus be approximated by

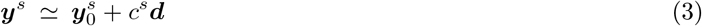

where 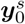 indicates initial activity in response to sensory input *s*, and ***d*** indicates the common direction along which activity acquires structure. The coefficients *c^s^*, which measure the strength of the components along the common direction **d**, are determined by category: they are approximately equal to c^A^if the sensory input is associated with category A, and *c*^B^ otherwise. Consequently, over learning, activity vectors associated with different categories are pushed apart along ***d***; this is illustrated in Figs. 3B and C, which show activity for the two circuits analyzed in the second and third column of Fig. 2, respectively. Activity thus forms two distinct clouds, one for each category; the centers of the two clouds along **d** are given, approximately, by *c*^A^ and *c*^B^. The mathematical framework detailed in Methods 4 allows us to derive closed-form expressions for the clustering direction ***d*** and the coefficients *c*^A^ and *c*^B^. In the next two sections, we take advantage of those expressions to determine how the different activity patterns shown in Fig. 2 depend on task and circuit parameters.

The fact that activity clusters by category tells us immediately that the category selectivity index of single neurons increases over learning, as observed in simulations (Figs. 2F and J). To see this quantitatively, note that from the point of view of a single neuron, *i*, Eq. 3 reads

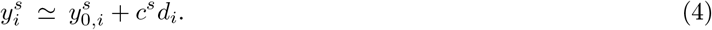

Since *c*^s^ is category-dependent, while *d_i_* is fixed, the second term in the right-hand side of Eq. 4 separates activity associated with different categories (Figs. 2E and I), and implies an increase in the category selectivity index (Eq. 2; Figs. 2F and J). The generality of Eq. 4 indicates that the increase in selectivity is a robust byproduct of gradient descent learning, and so can be observed in any circuit that learns the categorization task, regardless of model details. This explains the increase in selectivity consistently observed in simulations (Fig. 2F, J and Fig. S1A).

### Correlations reflect circuit and task properties

While the behaviour of category selectivity is consistent across all circuit models, the behaviour of population responses is not: as shown in Fig. 2, over learning responses can become negatively correlated and symmetric (Figs. 2G-H), or positively correlated and asymmetric (Figs. 2K-L). The reason is illustrated in Figs 3B and C. In Fig. 3B, the centers of the category clouds along ***d***, *c*^A^ and *c*^B^, have, respectively, a positive and a negative sign relative to the center of initial activity (denoted by a black triangle). As a consequence, the two clouds move in opposite directions. The population thus develops, over learning, negative category correlation (Fig. 2G-H): if the activity of a given neuron increases for one category, it decreases for the other, and vice versa. Furthermore, if *c*^A^ and *c*^B^ have similar magnitude (which is the case for Fig. 2G-H), activity changes for the two categories have similar amplitude, making the response to categories A and B approximately symmetric. In Fig. 3C, on the other hand, *c*^A^ and *c*^B^ are both positive; clouds associated with the two categories move in the same direction relative to the initial cloud of activity. This causes the population to develop positive category correlation (Fig. 2K-L): if the activity increases for one category, it also increases for the other, and similarly for a decrease. Because the magnitude of *c*^A^ is larger than *c*^B^, activity changes for category `A` are larger than for `B`, making the response to categories `A` and `B` asymmetric.

This analysis tells us that whether negative or positive category correlation emerges depends on the relative signs of *c*^A^ and *c*^B^. We can use mathematical analysis to compute the value and sign of *c*^A^ and *c*^B^, and thus predict how category correlation changes over learning (Methods 4.4). We find that the biophysical details of the circuit play a fundamental role in determining category correlation. In Fig. 4A, we show category correlation as a function of the threshold and gain of the readout neuron (Fig. 1C). We find that varying those can change the magnitude and sign of correlations, with positive correlations favoured by large values of the threshold and gain and negative correlations favoured by small values. Category correlation is also affected by the threshold and gain of neurons in the intermediate layer. This can be seen in Fig. 4B, which shows that larger values of the threshold and gain tend to favour positive correlation. An equally important role is played by the relative learning rates of the the readout, ***w***, and the intermediate weights, ***u***. As illustrated in Fig. 4C, increasing the ratio of the learning rates, *η_w_*/*η_u_*, causes the correlation to decrease. Overall, these results indicate that category correlation depends on multiple biophysical aspects of the circuit, which in turn are likely to depend on brain areas. This suggests that correlation can vary across brain areas, which is in agreement with the observation that positive correlations reported in monkeys area LIP are robust across experiments [18], but inconsistent with the correlations observed in monkeys PFC [7].

**Fig. 4:**
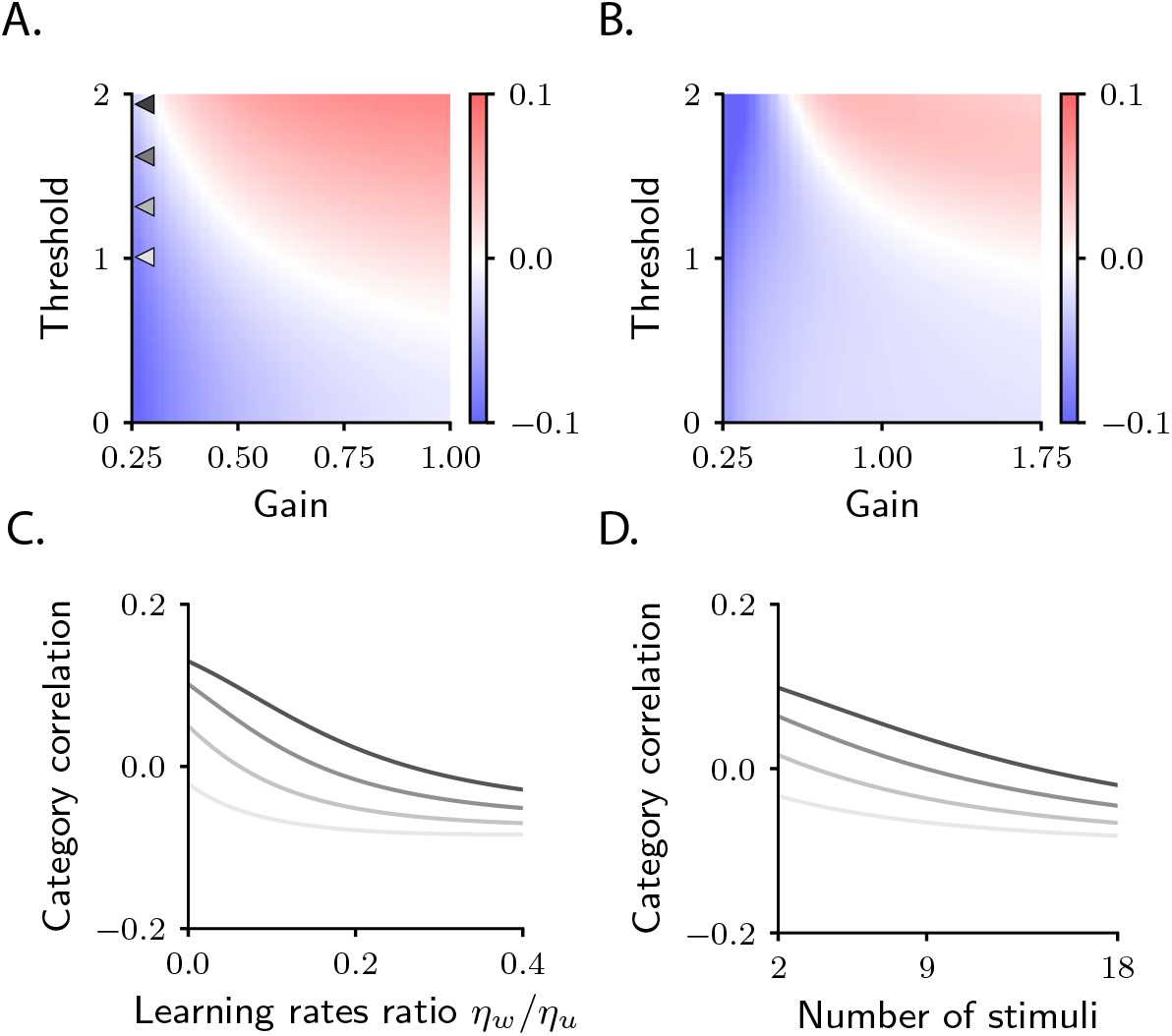
Category correlation depends on circuit and task properties. **A.** Category correlation as a function of the threshold and gain of the readout neuron. Grey arrows indicate the threshold and gain that are used in panels C and D. The learning rate ratio, *η_w_*/*η_u_*, is set to 0.4 here and in panel B and D. **B.** Category correlation as a function of the threshold and gain of neurons in the intermediate layer; details as in panel A. **C.** Category correlation as a function of the learning rate ratio. The threshold and gain of the readout neuron are given by the triangles indicated in panel A, matched by color. **D.** Category correlation as a function of the number of stimuli; same color code as in panel C. In all panels, correlations were computed from the approximate theoretical expression given in Methods 4.4 (Eq. 74). Parameters are summarized in Table 1 (Methods 6.2).

Category correlation also depends on the total number of stimuli, a property of the task rather than the circuit (Methods 4.4, Eq. 77). This is illustrated in Fig. 4D, which shows that increasing the number of stimuli causes a systematic decrease in correlation. The model thus makes the experimentally-testable prediction that increasing the number of stimuli should push category correlation from positive to negative values. This finding is in agreement with the fact that negative correlations are typically observed in sensory cortex, as well as machine-learning models trained on benchmark datasets [36] – that is, in cases where the number of stimuli is much larger than in the current task.

### Patterns of selectivity are shaped by initial connectivity

We conclude our analysis of the simple categorization task by taking a closer look at category selectivity. We have already observed, in Figs. 2F and J, that the category selectivity of neurons in the intermediate layer increase over learning. However, as shown in those figures, the amount it increases can vary markedly across the population – a finding that reproduces the variability commonly seen in experiments [4–6]. The model naturally explains this variability: as can be seen in Eq. 4, the magnitude of category-related activity changes (and, consequently, the magnitude of category selectivity) depends, for a given neuron *i*, on the magnitude of *d_i_*. Mathematical analysis (see Methods 4.2, especially Eq. 55) indicates that, for the current task, the category direction ***d*** is approximately aligned with the vector that specifies connectivity between the intermediate and the readout neurons, ***w***, before learning starts; we denote this vector ***w***_0_ (Fig. 3B-C). As a consequence, only neurons that are initially strongly connected to the readout neuron – that is, neurons for which *w*_0,*i*_ is large – exhibit a large selectivity index (Figs. 5B-C).

**Fig. 5:**
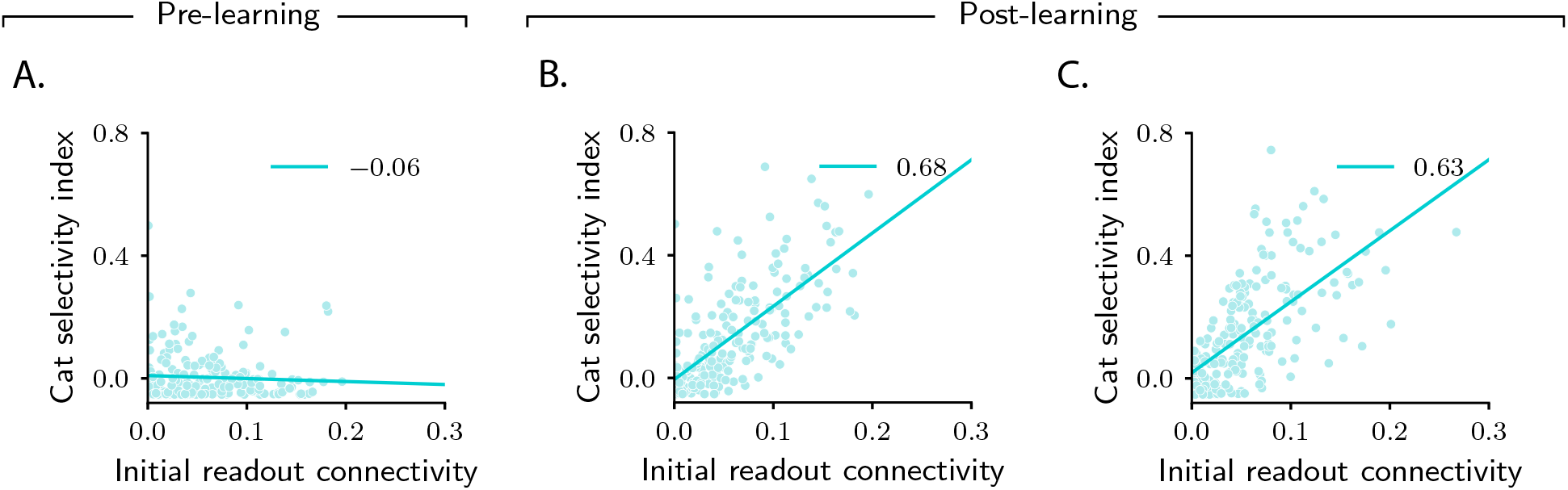
Magnitude of category selectivity depends on connectivity with the readout neuron. **A, B, C.** Category selectivity as a function of the initial readout connectivity *w*_0,*i*_ (in absolute value). The three columns are as in Fig. 2: pre-learning (first column) and post-learning for two different circuits (second and third columns). Each dot represents a neuron in the intermediate layer. Cyan line: linear fit, with Pearson correlation coefficient shown in the figure legend.

Why does activity cluster along the initial readout ***w***_0_? As described above, the output of the circuit, *z*, depends on the dot product ***w*** · ***y***, where ***w*** are the readout weights after learning. Consequently, the final activity in the intermediate layer, ***y***, must include a category-dependent component along ***w***. Such a component can be aligned either with the initial readout weights, ***w***_0_, either with the readout weight changes. The fact that activity changes are mostly aligned with ***w***_0_ indicates that the learning algorithm is characterized by a sort of inertia, which makes it rely on initial connectivity structure much more heavily than on the learned one. As showed in Methods 3.1, this is a property of networks with a large number of neurons relative to the number of stimuli, which are characterized by small weight changes [32].

In terms of biological circuits, Fig. 5 predicts that changes in selectivity are determined by the strength of synaptic connections a neuron makes, before learning, to downstream readout areas. Experiments consistent with this prediction have been recently reported: studies in rodents prefrontal cortex [13, 37] found that all neurons which were highly selective to a given abstract variable were characterized by similar downstream projections (i.e., they projected to the same area). These experiments would provide evidence for our model if two conditions were met. First, neurons in the downstream area should behave as readout neurons: over learning, their activity should increasingly depend on the abstract variable. Second, the strength of the synaptic connections that neurons make to downstream neurons should correlate with selectivity (Figs. 5B-C). Both predictions could be tested with current experimental technology.

In sum, we analyzed activity in the intermediate layer of circuits that learned the simple categorization task. We found that activity gets reshaped along a common, stimulus-independent direction (Eq. 3), which is approximately aligned with the initial readout vector ***w***_0_. Activity vectors associated with different categories develop two distinct clouds along this direction – a fact that explains the increase in category selectivity observed in Figs. 2F and J. We also found that the sign of the category correlation depends on the circuit (threshold and gain of neurons in the intermediate and readout layers, and relative learning rates) and on the task (number of stimuli). Modifying any of these can change the direction the clouds of activity move along ***w***_0_, which in turn changes the sign of category correlation, thus explaining the different behaviours observed in Figs. 2G-H and K-L.

### Evolution of activity during the context-dependent categorization task

We now consider a more complex categorization task. Here, stimuli-category associations are not fixed, but context-dependent: stimuli that are associated with category `A` in context `1` are associated with category `B` in context `2`, and vice versa. Context-dependent associations are core to a number of experimental tasks [5, 8–10, 38], and are ubiquitous in real-world experience.

In the model, the two contexts are signaled by distinct sets of context cues (for example, two different sets of visual stimuli) [8, 9]. As for the stimuli, context cues are represented by random and orthogonal sensory input vectors. On every trial, one stimulus and one context cue are presented; the corresponding sensory inputs are combined linearly to yield the total sensory input vector ***x**^s^* (Methods 5.1). This task is computationally much more involved than the previous one, primarily because context induces nontrivial correlational structure: in the simple task, all sensory input vectors were uncorrelated; that is no longer true in the context-dependent task. For instance, two sensory inputs with the same stimulus and different context cues are highly correlated. In spite of this high correlation, though, they can belong to different categories – for instance, when context cues are associated with different contexts. In contrast, two sensory inputs with different stimuli and different context cues are uncorrelated, but they can belong to the same category. From a mathematical point of view, this correlational structure makes sensory input vectors non linearly-separable. This is in stark contrast to the simple task, for which sensory input vectors were linearly-separable [39]. In fact, this task is a generalization of the classical XOR task where, rather than just two stimuli and two context cues, there are more than two of each [38]. In the example shown below, we used 8 stimuli and 8 context cues.

We are again interested in understanding how activity in the intermediate layer evolves over learning. We start by investigating this via simulations (Fig. 6). As in Figs. 2B, F and J, we first measure category selectivity (Eq. 2). Before learning, activity is characterized by small selectivity, which is weakly negative on average (Fig. 6A; the fact that average category selectivity is initially weakly negative is due to the composite nature of inputs for this task, see Methods 5.7). Over learning, the average category selectivity increases (Fig. 6D). We tested the robustness of this behaviour by varying the parameters that control both the circuit (threshold and gain of neurons, learning rates) and task (number of stimuli and context cues). As in the simple task, we found that the average category selectivity increases in all circuit models, regardless of the parameters (Figs. 6G and S7A).

**Fig. 6:**
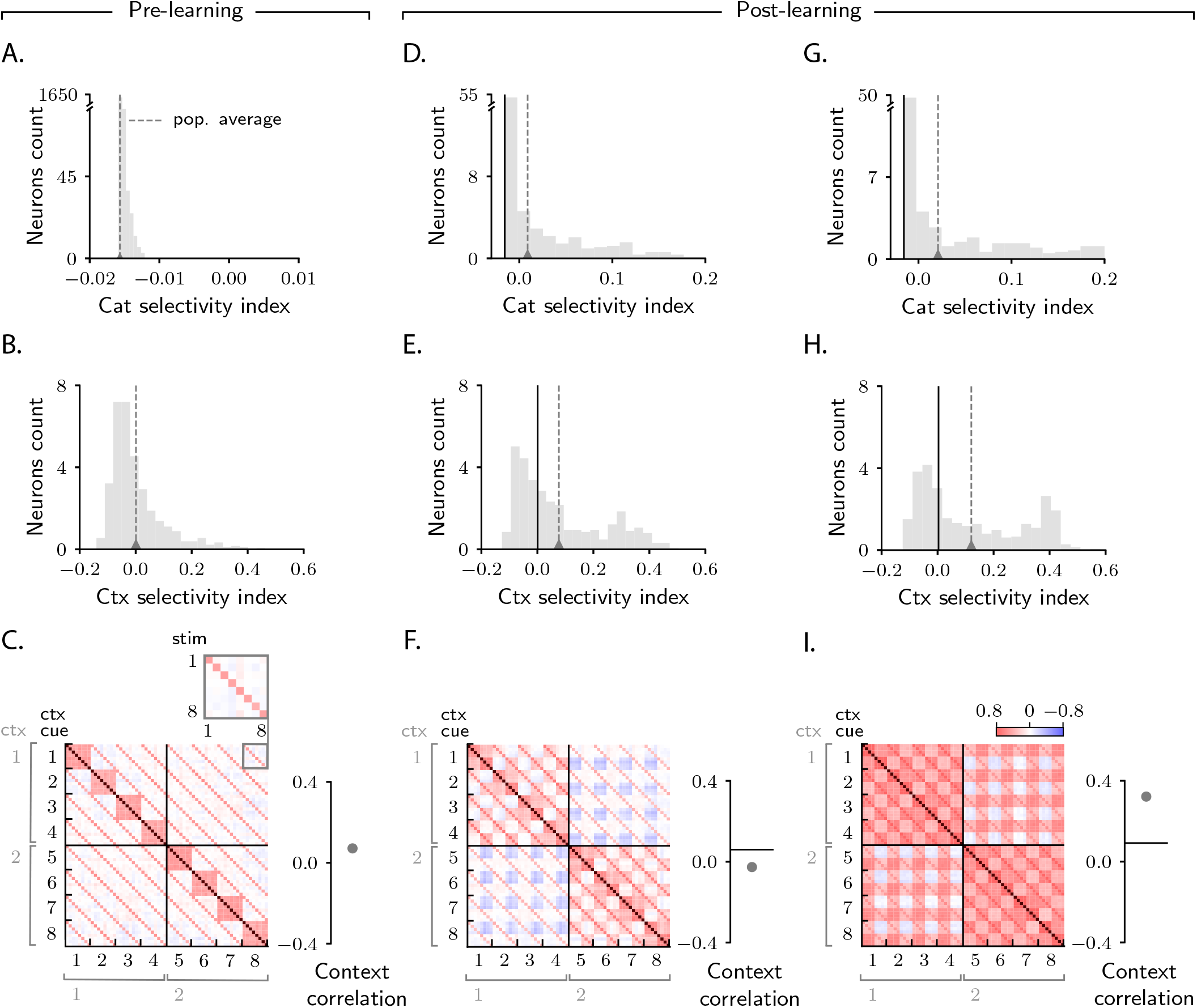
Characterization of activity evolution during the context-dependent categorization task. Results from simulations. The first column (A-C) shows a naive circuit (pre-learning); the second (D-F) and third (G-I) columns show two trained circuits (post-learning), characterized by different sets of parameters. **A, D, G.** Histogram of category selectivity (Eq. 2) across the population of neurons in the intermediate layer (note that the vertical axis has been expanded for visualization purposes). Grey dashed lines indicate the average selectivity across the population. In panels D and G, the black vertical lines indicate the initial value of the average selectivity. Note that the distribution of category selectivity is different from the distribution observed in the simple task (Fig. 2F, J); the distribution is now heavy-tailed, with only a fraction of the neurons acquiring strong category selectivity (see also Fig. 8B). **B, E, H.** Histogram of context selectivity (Methods 5.3, Eq. 122), details as in A, D and **G. C, F, I.** Correlation matrices. Each entry shows the Pearson correlation coefficient between activity from different trials. There are 8 stimuli and 8 context cues, for a total of 64 trials (i.e., 64 stimulus/context cue combinations). Diagonal entries (brown) are all equal to 1. The inset on the top of panel C shows, as an example, a magnified view of correlations among trials with context cues 1 and 8, across all stimuli (1 to 8). To the right of the matrices we show the context correlation, defined to be the average of the correlations within the off-diagonal blocks (trials in different contexts). In panels F and I, the black horizontal lines indicate the initial value of context correlation. Parameters are summarized in Table 1 (Methods 6.2).

While in the simple task we could only investigate the effect of category on activity, in this task we can also investigate the effect of context. For this we measure context selectivity which, analogously to category selectivity, quantifies the extent to which single-neuron activity is more similar within than across contexts (Methods 5.3, Eq. 122). Context selectivity is shown in Figs. 6B and E. We find, as we did for category selectivity, that average context selectivity increases over learning – a behaviour that is in agreement with experimental findings [8, 9]. The increase in context selectivity is, as for category, highly robust, and does not depend on model details (Figs. 6H and S7A).

Finally, we analyze signal correlations; these are summarized in the correlation matrices displayed in Figs. 6C, F and I. As we used 8 stimuli and 8 context cues, and all stimuli-context cues combinations are permitted, each correlation matrix is 64 × 64. Trials are sorted according to context cue first and stimulus second; with this ordering, the first half of trials corresponds to context `1` and the second half to context `2`, and the off-diagonal blocks are given by pairs of trials from different contexts.

Figure 6C shows the correlation matrix before learning. Here the entries in the correlation matrix are fully specified by sensory input, and can take only three values: large (brown), when both the stimuli and the context cues are identical across the two trials; intermediate (red), when the stimuli are identical but the context cues are not, or vice versa; and small (white), when both stimulus and context cues are different. Figures 6F and I show correlation matrices after learning for two circuits characterized by different parameters. As in the simple task, the matrices acquire additional structure during learning, and that structure can vary significantly across circuits (Figs. 6F and I). To quantify this, we focus on the off-diagonal blocks (pairs of trials from different contexts) and measure the average of those correlations, which we refer to as context correlation. Context correlation behaves differently in the two circuits displayed in Fig. 6F and I: it decreases over learning in Fig. 6F, whereas it increases in Fig. 6I. Thus, as in the simple task, the behaviour of correlations is variable across circuits. This variability is not restricted to context correlation: as in the simple task, category correlation is also variable (Fig. S7A), and the population response to categories `A` and `B` can be symmetric or asymmetric depending on model details (Figs. S8A-B).

### Analysis of the context-dependent categorization task

To uncover the mechanisms that drive learning-induced activity changes, we again analyse the circuit mathematically. The addition of context makes the analysis considerably more complicated than for the simple task; most of the details are thus relegated to Methods 5; here we discuss the main results.

Figure 7A shows, schematically, activity before learning (see Fig. S7D for simulated data). Each point represents activity on a given trial, and is associated with a category (A, orange; B, blue) and a context (1, circles; 2, squares). Before learning, activity is mostly unstructured (Figs. 7A, Methods 5.7); over learning, though, it acquires structure (Figs.7B and C). As in the simple task (Figs. 3B-C), activity vectors get re-arranged into statistically-distinguishable clouds. While in the simple task clouds were determined by category, here each cloud is associated with a combination of category and context. As a result, four clouds are formed: the cloud of orange circles corresponds to category `A` and context `1`; orange squares to category `A` and context `2`; blue circles to category `B` and context `1`; and blue squares to category `B` and context `2`.

**Fig. 7:**
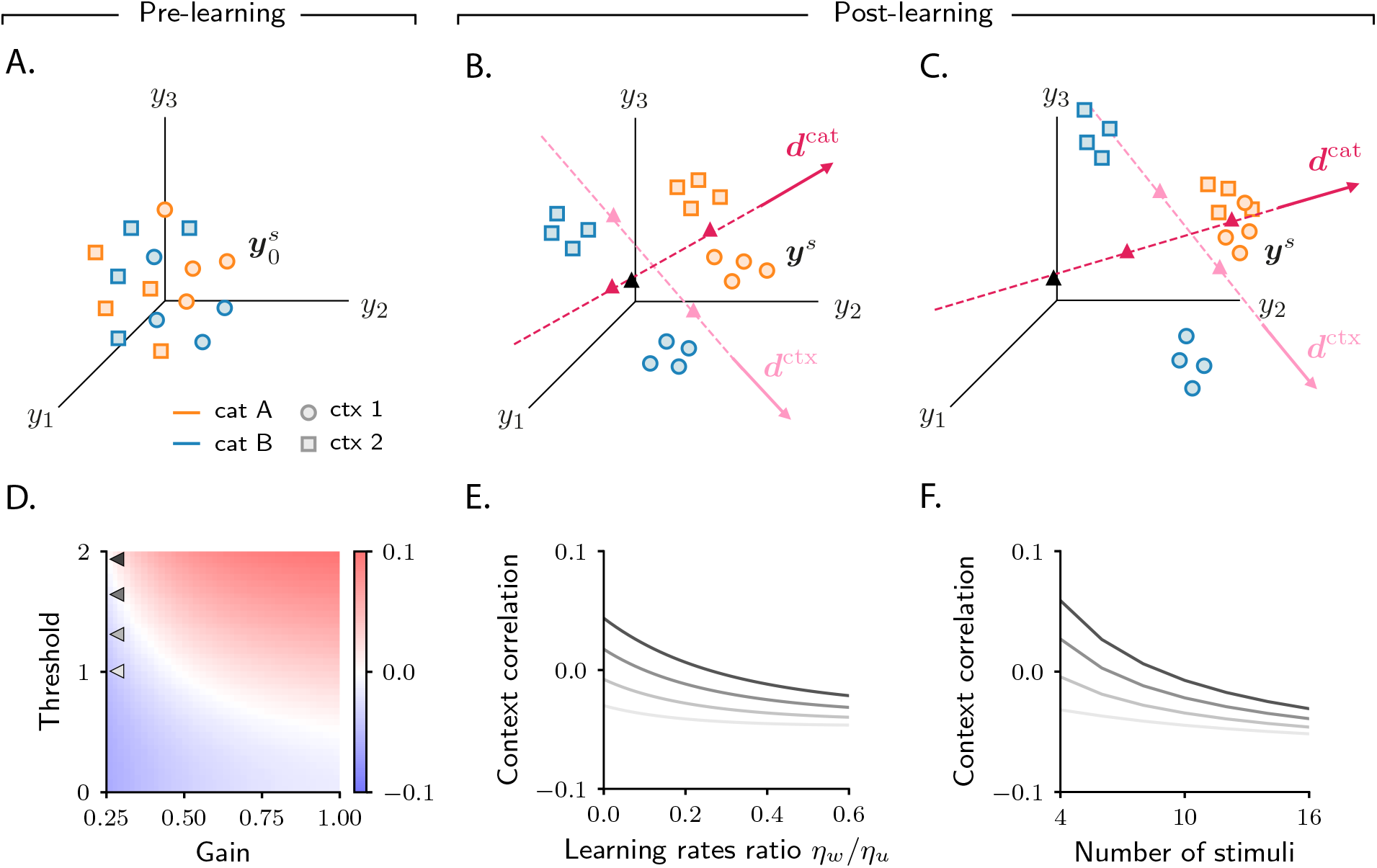
Analysis of activity evolution during the context-dependent categorization task. Results from mathematical analysis. **A, B, C.** Cartoons illustrating how activity evolves over learning. Orange and blue symbols are associated with categories `A` and `B`, respectively; circles and squares are associated with contexts `1` and `2`. Before learning, activity is mostly unstructured (panel A). After learning, activity forms four clouds, one for each combination of category and context. The center of the activity vectors associated with categories `A` and `B` and context `1` and `2` are indicated, respectively, by magenta and pink triangles. The black triangle indicates the center of initial activity. The cartoons in panels A-B-C refer to the three circuits illustrated in the three columns of Fig. 6; for illustration purposes, we show a reduced number of stimuli and context cues (4 instead of 8). Simulated data from the circuits displayed in Fig. 6 are shown in Figs. S7C-D. **D.** Change in context correlation over learning as a function of the threshold and gain of the readout neuron. Grey arrows indicate the threshold and gain that are used in panels E and F. **E.** Change in context correlation over learning as a function of the ratio of learning rates in the two layers. **F.** Change in context correlation over learning as a function of the number of stimuli. Correlations in panels D-F were computed from the approximate theoretical expression given in Methods 5.4. Parameters are given in Table 1 (Methods 6.2).

The transition from unstructured activity (Fig. 7A) to four clouds of activity (Figs. 7B-C) occurs by learning-induced movement along two directions: ***d***^cat^, which corresponds to category, and ***d***^ctx^, which corresponds to context. Activity vectors in different categories move by different amounts along ***d***^cat^; this causes the orange and blue symbols in Figs. 7B-C to move apart, so that activity vectors associated with the same category become closer than vectors associated with opposite categories. As in the simple task, this in turn causes the category selectivity to increase, as shown in Figs. 6D and G (Methods 5.7). Similar learning dynamics occurs for context: activity vectors from different contexts move by different amounts along ***d***^ctx^. This causes the squares and circles in Figs. 7B-C to move apart, so that activity vectors from the same context become closer than vectors from different contexts. Again, this in turn causes the context selectivity to increase, as shown in Figs. 6E and H (Methods 5.6). Mathematical analysis indicates that the increase in clustering by category and context is independent of model parameters (Fig. S7B), which explains the robustness of the increase in selectivity observed in simulations.

The category- and the context-related structures that emerge in Figs. 7B and C have different origins and different significance. The emergence of category-related structure is, perhaps, not surprising: over learning, the activity of the readout neuron becomes category-dependent, as required by the task; such dependence is then directly inherited by the intermediate layer, where activity clusters by category. This structure was already observed in the simple categorization task (Fig. 3B-C). The emergence of context-related structure is, on the other hand, more surprising, since the activity of the readout neuron does not depend on context. Nevertheless, context-dependent structure, in the form of clustering, emerges in the intermediate layer activity. Such novel structure is a signature of the gradient-descent learning rule used by the circuit [40]. The mechanism through which context clustering emerges is described in detail in Methods 5.6. But, roughly speaking, context clustering emerges because, for a pair of sensory inputs, how similarly their intermediate-layer representations evolve during learning is determined both by their target category and their correlations (Eq. 27, Methods 3). In the simple task, initial correlations were virtually nonexistent (Fig. 2C), and thus activity changes were specified only by category; in the context-dependent task, initial correlations have structure (Fig. 6C), and that structure critically affects neural representations. In particular, inputs with the same context tend to be relatively correlated, and those are also likely to be associated with the same category; their representations are thus clustered by the learning algorithm, resulting in context clustering.

While the clustering by category and context described in Figs. 7B and C is robust across circuits, the position of clouds in the activity space is not. As in the simple task, the variability in cloud position explains the variability in context correlation (although the relationship between clouds position and correlations is more complex in this case, see Methods 5). In Figs. 7D-F, we show how context correlation depends on model parameters. This dependence is qualitatively similar to that of category correlation in the simple task: context correlation depends on the threshold and gain of neurons (compare Figs. 7D and 4A), on the relative learning rate *η_w_*/*η_u_* (compare Figs. 7E and 4C), and on the number of stimuli (compare Figs. 7F and 4D). However, we find that the region of parameter space leading to an increase in correlation shrinks substantially compared to the simple task (Fig. S8C, see also Methods 5.2); this is in line with the observation that correlations decrease to negative values when the complexity of the task increases, as shown in Fig. 4D.

### Patterns of pure and mixed selectivity are shaped by initial activity

As a final step, we take a closer look at single-neuron selectivity. Analysis from the previous sections indicates that the average selectivity to both category and context increases over learning. And, as in the simple task, the increase is highly variable across neurons (Figs. 6D-E and G-H). To determine which neurons become the most selective to category and context, we analyze the directions along which clustering to category and context occurs, ***d***^cat^ and ***d***^ctx^ (Fig. 7B-C). In analogy with the simple task, neurons that strongly increase selectivity to category are characterized by a large component along the category direction ***d***^cat^; similarly, neurons that strongly increase selectivity to context are characterized by a large component along the context direction ***d***^ctx^ (Figs. S9A-B).

Analysis in Methods 5.8 shows that both the category and context directions, ***d***^cat^ and ***d***^ctx^, are strongly correlated with the initial readout vector ***w***_0_. As in the simple task, this leads to the prediction that neurons that strongly increase selectivity to either category or context are, before learning, strongly connected to the downstream readout neuron (Fig. 8A).

**Fig. 8:**
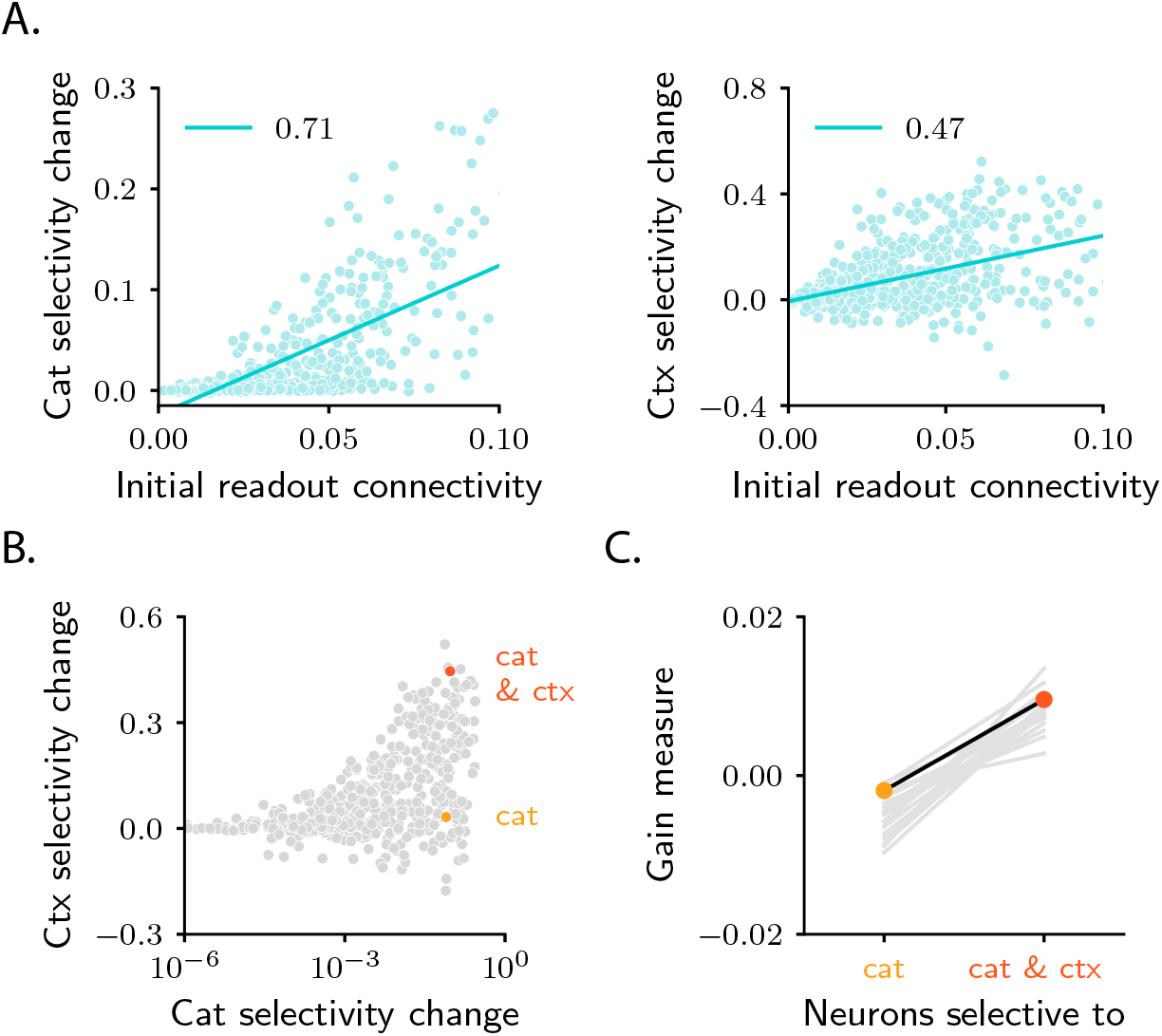
Patterns of pure and mixed selectivity to category and context. **A.** Changes in category selectivity (left) and context selectivity (right) as a function of the initial readout connectivity, *w*_0,*i*_ (in absolute value). Details as in Figs. 5B-C. **B.** Changes in context selectivity as a function of changes in category selectivity. Note the logarithmic scale on the x-axis; this is required by the heavy-tailed behaviour of category selectivity (Figs. 6D and G). We highlighted two sample neurons: one with strong, pure selectivity to category (yellow) and one with strong, mixed selectivity to category and context (orange). **C.** Neurons that develop pure and mixed selectivity are characterized by different patterns of initial activity. Here, we plot the gain-based measure of activity defined in Eq. 183 for neurons that belong to the former (left), and the latter (right) group. The former group includes neurons for which the change in category selectivity, but not the change in context selectivity, is within the top 15% across the population. The latter group includes neurons for which the change in both category and context selectivity is within the top 15%. Dots show results for the circuit analyzed in panels A and B. Grey lines show results for 20 different circuit realizations; note that the slope is positive for all circuits. All panels in the figure show results for the circuit displayed in the second column of Fig. 6; the circuit displayed in the third column yields qualitatively similar results (Figs. S9C-D).

Although ***d***^cat^ and ***d***^ctx^ are both correlated with **w**_0_, they are not perfectly aligned (Methods 5.8). In principle, then, for a given neuron (here, neuron *i*), both 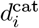 and 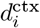 could be large (implying mixed selectivity to both abstract variables, category and context), or only one could be large (implying pure selectivity to only one abstract variable, category or context), or both could be small (implying no selectivity at all). While all combinations are possible in principle, in the model they do not all occur. In Fig. 8B, we plot changes in context selectivity as a function of changes in category selectivity. We observe that, among all the neurons that strongly increase their selectivity, some increase selectivity to both category and context (orange sample neuron), and others increase selectivity to category, but not context (yellow sample neuron). In contrast, none increases selectivity to context but not category. This makes the following experimental prediction: among all the neurons that are strongly connected to the readout, neurons with pure selectivity to category and neurons with mixed selectivity to category and context should be observed, but neurons with pure selectivity to context should not. The asymmetry between category and context arises because, in the model, the readout neuron learns to read out category, but not context. We show in Figs. S9E-F that if a second readout neuron, which learns to read out context, is included in the circuit, neurons with strong pure selectivity to context are also observed.

What determines whether a given neuron develops pure selectivity to category, or mixed selectivity to category and context? Analysis reported in Methods 5.8 indicates that these two populations are characterized by different properties of the initial activity. In particular, the two populations are characterized by different initial patterns of the response gain (defined as the slope of the activation function, Fig. 1C, at the response), which measures the local sensitivity of neurons to their inputs. The exact patterns that the response gain takes across the two populations is described in detail in Methods 5.8 (Eq. 183); the fact that pure- and mixed-selective neurons can be distinguished based on these patterns is illustrated in Fig. 8C. Overall, these results indicate that initial activity, which is mostly unstructured and task-agnostic, plays an important role in learning: it breaks the symmetry among neurons in the intermediate layer, and determines which functional specialization neurons will display over learning.

## Discussion

How does the brain learn to link sensory stimuli to abstract variables? Despite decades of experimental [1, 2, 4, 5] and theoretical [39, 41, 42] work, the answer to this question remains elusive. Here we hypothesized that learning occurs via gradient descent synaptic plasticity. To explore the implications of this hypothesis, we considered a minimal model: a feedforward circuit with one intermediate layer, assumed to contain a large number of neurons compared to the number of stimuli. This assumption allowed us to thoroughly analyze the model, and thus gain insight into how activity evolves during learning, and how that evolution supports task acquisition.

We focused on two categorization tasks: a simple one (Fig. 2), in which category was determined solely by the stimulus, and a complex one (Fig. 6), in which category was determined by both the stimulus and the context. We showed that, over learning, single neurons become selective to abstract variables: category (which is explicitly reinforced) and context (which is not; instead, it embodies the task structure, and is only implicitly cued). From a geometrical perspective, the emergence of selectivity during learning is driven by clustering: activity associated with stimuli in different categories is pushed apart, forming distinct clusters (Fig. 3). In the context-dependent task, additional clustering occurs along a second, context-related axis; this results in activity forming four different clouds, one for each combination of category and context (Fig. 7). While the behaviour of selectivity is highly stereotyped, the behaviour of signal correlations and tuning symmetry is not, but depends on details (Fig. 4). From a geometrical perspective, the variability in correlations and symmetry is due to the variability in the position of category and context clusters with respect to initial activity.

Our work was motivated partly by the observation that responses to different categories in monkeys area LIP were positively-correlated and asymmetric [18] – a finding that seems at odds with experimental observations in sensory, and other associative, brain areas [5, 7]. It has been suggested that those responses arise as a result of learning that drives activity in area LIP onto an approximately one-dimensional manifold [18, 43, 44]. Our results are broadly in line with this hypothesis: for the simple categorization task, which is similar to [18], we showed that activity stretches along a single direction (Eq. 3, Fig. 3C). Analysis in Methods 3.2 further shows that not only at the end of learning, but at every learning epoch, activity is aligned along a single direction; the whole learning dynamics is thus approximately one-dimensional. However, in the context-dependent categorization task, activity stretches along two dimensions (Fig. 7B-C), indicating that one dimension does not always capture activity.

Our analysis makes several experimental predictions. First, it makes specific predictions about how category and context correlations should vary with properties of the circuit (threshold and gain of neurons, relative learning rates) and with the task (number of stimuli, context-dependence) (Fig. 4). These could be tested with current technology; in particular, testing the dependence on task variables only requires recording neural activity. Second, it predicts that selectivity is shaped by connectivity with downstream areas, a result that is in line with recent experimental observations [13, 37, 45, 46]. More specifically, it predicts that, for a given neuron, selectivity correlates with the strength of the synaptic connection that the neuron makes to the downstream neurons that read out category (Figs. 5B-C and 8A). Across all neurons that are strongly connected to downstream readout neurons, selectivity to category and context is distributed in a highly stereotyped way: during learning, some neurons develop mixed selective to category and context, others develop pure selectivity to category, but none develop pure selectivity to context (Fig. 8B). Moreover, whether a neuron develops mixed or purse selectivity depends on initial activity (Fig. 8C).

### Previous models for categorization

Previous theoretical studies have investigated how categorization can be implemented in multi-layer neural circuits [39, 42, 47–51]. Several of those studies considered a circuit model in which the intermediate connectivity matrix, ***u***, is fixed, and only the readout vector, ***w***, evolves (via Hebbian plasticity) over learning [39, 47, 48]. This model can learn both the simple (linearly-separable) and complex (non-linearly separable) tasks [39]. Because there is no learning in the intermediate connectivity, activity in the intermediate layer remains unstructured, and high dimensional, throughout learning. This stands in sharp contrast to our model, where learning leads to structure in the form of clustering – and, thus, a reduction in activity dimensionality.

One study did consider a model in which both the intermediate and the readout connectivity evolve over learning, according to reward-modulated Hebbian plasticity [42]. This circuit could learn a simple categorization task but, in contrast to our study, learning did not lead to significant changes in the activity of the intermediate layer. When feedback connectivity was introduced, learning did lead to activity changes in the intermediate layer, and those activity changes led to an increase in category selectivity – a finding that is in line with ours. There were, however, several notable differences relative to our model. First, learning of the intermediate and readout weights occurred on separate timescales: the intermediate connectivity only started to significantly change when the readout connectivity was completely rewired; in our model, in contrast, the two set of weights evolve on similar timescales. Second, population responses were negatively-correlated and symmetric; whether positively-correlated and asymmetric responses (as seen in experiments [18], and in our model) can also be achieved remains to be established. Third, context-dependent associations, that are core to a variety of experimental tasks [5, 8, 10, 30, 38, 52], were not considered. Whether reward-modulated Hebbian plasticity can be used to learn context-dependent tasks is unclear, and represents an important avenue for future work.

### Gradient descent learning in the brain

A common feature of the studies described above is that learning is implemented via Hebbian synaptic plasticity – a form of plasticity that is known to occur in the brain. Our model, on the other hand, uses gradient descent learning in a multi-layer network, which requires back-propagation of an error signal; whether and how such learning is implemented in the brain is an open question [53]. A number of recent theoretical studies have proposed biologically-plausible architectures and plasticity rules that can approximate back-propagation on simple and complex tasks [21–27]. Understanding whether these different implementations lead to differences in activity represents a very important direction for future research. Interestingly, recent work has showed that it is possible to design circuit models where the learning dynamics is identical to the one studied in this work, but the architecture is biologically plausible [27]. We expect our results to directly translate to those models. Other biologically-plausible setups might be characterized, instead, by different activity evolution. Recent work [54, 55] made use of a formalism similar to ours to describe learning dynamics induced by a number of different biologically-plausible algorithms and uncovered non-trivial, qualitatively different dynamics. Whether any of these dynamics leads to different neural representations in neuroscience-inspired categorization tasks like the ones we studied here is an open, and compelling, question.

In this work, we used mathematical analysis to characterize the activity changes that emerge during gradient descent learning. Our analysis relied on two assumptions. First, the number of neurons in the circuit is large compared to the number of stimuli to classify. Second, the synaptic weights are chosen so that the initial activity in all layers of the network lies within an intermediate range (i.e., it neither vanishes nor saturates) before learning starts [32–34, 56]. These two assumptions are reasonable for brain circuits, across time scales ranging from development to animals’ lifetimes; a discussion on the limitations of our approach is given in Methods 3.4.

A prominent feature of learning under these assumptions is that changes in both the synaptic weights and activity are small in amplitude (Methods 3). This has an important implication: the final configuration of the circuit depends strongly on the initial one. We have showed, for example, that the selectivity properties that single neurons display at the end of learning are determined by their initial activity and connectivity (Figs. 5B-C and 8A and C). Moreover, the final distribution of selectivity indices, and the final patterns of correlations, bear some resemblance to the initial ones (see, e.g., Fig. 6); for this reason, we characterized activity evolution via changes in activity measures, rather than their asymptotic, post-learning values. Overall, these findings stress the importance of recording activity throughout the learning process to correctly interpret neural data [57, 58].

Circuits that violate either of the two assumptions discussed above may exhibit different gradient descent learning dynamics than we saw in our model [56], and could result in different activity patterns over learning.

Previous studies have analyzed circuits with linear activation functions and weak connectivity (weak enough that activity is greatly attenuated as it passes through the network). However, linear activation functions can only implement a restricted set of tasks [59–61]; in particular, they cannot implement the context-dependent task we considered. Developing tools to analyze arbitrary circuits will prove critical to achieving a general understanding of how learning unfolds in the brain [62–64].

### Beyond simplified models

Throughout this work, we focused on two simplified categorization tasks, aimed at capturing the fundamental features of the categorization tasks commonly used in systems neuroscience [4, 6, 8]. The mathematical framework we developed to analyze those tasks could, however, easily be extended in several directions, including tasks with more than two categories [5, 6, 30] and tasks involving generalization to unseen stimuli [39, 40]. An important feature missing in our tasks, though, is memory: neuroscience tasks usually involve a delay period during which the representation of the output category must be sustained in the absence of sensory inputs [4, 6, 8]. Experiments indicate that category representations are different in the stimulus presentation and the delay periods [4]. Investigating these effects in our tasks would require the addition of recurrent connectivity to the model. Mathematical tools for analyzing learning dynamics in recurrent networks is starting to become available [65–68], which could allow our analysis to be extended in that direction.

To model categorization, we assumed a quadratic function for the error 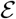 (Methods 2.1)– an assumption that effectively casts our categorization tasks into a regression problem. This made the model amenable to mathematical analysis, and allowed us to derive transparent equations to characterize activity evolution. Recent machine learning work has showed that, at least in some categorization setups [69], a cross-entropy function might result in better learning performance. The mathematical framework used here is, however, not well suited to studying networks with such an error function [33]. Investigating whether and how our findings extend to networks trained with a cross-entropy error function represents an interesting direction for future work.

Finally, in this study we focused on a circuit model with a single intermediate layer. In the brain, in contrast, sensory inputs are processed across a number of stages within the cortical hierarchy. Our analysis could easily be extended to include multiple intermediate layers. That would allow our predictions to be extended to experiments involving multi-area recordings, which are increasingly common in the field [70]. Current recording techniques, furthermore, allow monitoring neural activity throughout the learning process [5, 70]; those data could be used in future studies to further test the applicability of our model.

### Bridging connectivity and selectivity

In this work, we considered a circuit with a single readout neuron, trained to discriminate between two categories. One readout neuron is sufficient because, in the tasks we considered, categories are mutually exclusive [18]. We have found that the initial readout weights play a key role in determining the directions of activity evolution, suggesting that circuits with different or additional readout neurons might lead to different activity configurations. For example, one might consider a circuit with two readout neurons, each highly active in response to a different category. And indeed, recent work in mouse PFC suggests that two readout circuits are used for valence – one strongly active for positive valence, and one strongly active for negative one [37]. Also, in context-dependent tasks, one might consider a circuit with an additional readout for context. We have showed in Figs. S9E-F that this model leads to different experimental predictions for selectivity than the model with only one readout for category (Fig. 8B). Altogether, these observations indicate that functional properties of neurons are tighly linked to their long-range projections – a pattern that strongly resonates with recent experimental findings [13, 71]. Constraining model architectures with connectomics, and then using models to interpret neural recordings, represents a promising line of future research.

## Acknowledgements

F.M. would like to thank Friedrich Schuessler for useful discussions. F.M., N.H. and P.E.L. were supported by the Gatsby Charitable Foundation, and F.M. and P.E.L. were supported by the Wellcome Trust (110114/Z/15/Z).

## Methods

### 1 Overview

In the main text, we made qualitative arguments about the evolution of activity over learning. Here we make those arguments quantitative. We start with a detailed description of the circuit model (Section 2). We then derive approximate analytical expressions that describe how activity in the circuit evolves over learning (Section 3). To this end, we use an approach that is valid for large circuits. We apply this approach first to the simple task (Section 4), then to the context-dependent one (Section 5). Finally, we provide details on the numerical implementation of circuit models and analytical expressions (Section 6).

### 2 Model

#### 2.1 Circuit

We consider a feedforward circuit with a single intermediate layer (Fig. 1B). For simplicity, we assume that the input and the intermediate layer have identical size *N*, and we consider *N* to be large. The sensory input vector is indicated with ***x***. Activity in the intermediate layer reads

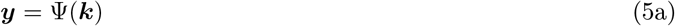

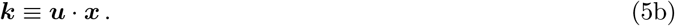

Here, ***k*** represents the synaptic drive and ***u*** is an *N* × *N* connectivity matrix. Activity in the readout layer is given by

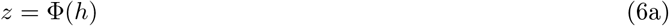

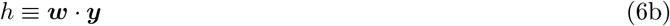

where *h* is the synaptic drive and ***w*** is an *N*-dimensional readout vector.

The activation functions *Ψ* and *Φ* are non-negative, monotonically increasing functions that model the input-to-output properties of units in the intermediate and readout layer, respectively. In simulations, we use sigmoidal functions,

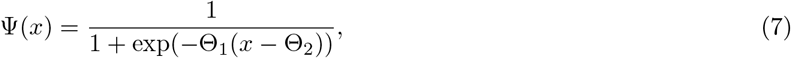

and similarly for *Φ*(*x*) (Fig. 1C). The parameters of the activation functions, Θ_1_ and Θ_2_, determine the gain and threshold, respectively, with the gain (defined to be the slope at *x* = Θ_2_) given by Θ_1_/4. Their values, which vary across simulations, are given in Section 6.2, Tables 1 and 2.

**Table 1:**
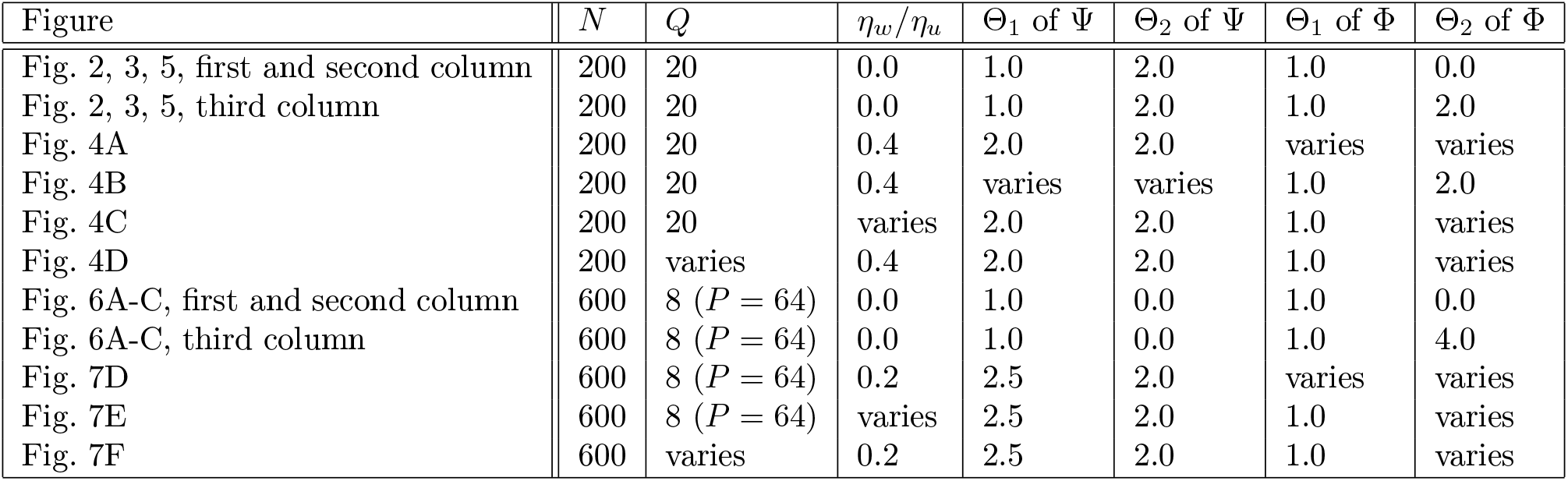
Table of parameters for Figures in the main text.

**Table 2:** Table of parameters for Supplementary Figures.

| Supplementary Figure | $N$ | $Q$ | $\eta_w/\eta_u$ | $\Theta_1$ of $\Psi$ | $\Theta_2$ of $\Psi$ | $\Theta_1$ of $\Phi$ | $\Theta_2$ of $\Phi$ |
| --- | --- | --- | --- | --- | --- | --- | --- |
| Fig. S1A | 200 | varies | varies | varies | varies | varies | varies |
| Fig. S2E | 200 | 20 | 0.0 | 1.0 | 2.0 | 1.0 | 0.0 |
| Fig. S4, first column | varies | 20 | 0.0 | 1.0 | 0.0 | 1.0 | 0.0 |
| Fig. S4, second column | 200 | 20 | varies | 1.0 | 0.0 | 1.0 | 0.0 |
| Fig. S4, third column | 200 | 20 | 0.0 | 1.0 | varies | 1.0 | 0.0 |
| Fig. S5, first column | varies | 20 | 0.0 | 1.0 | 0.0 | 1.0 | 2.0 |
| Fig. S5, second column | 200 | 20 | varies | 1.0 | 0.0 | 1.0 | 2.0 |
| Fig. S5, third column | 200 | 20 | 0.0 | 1.0 | varies | 1.0 | 2.0 |
| Fig. S6A-B | 200 | 12 | 0.0 | 1.0 | 2.0 | 1.0 | 0.0 |
| Fig. S6C | 200 | 12 | 0.0 | 1.0 | 2.0 | 1.0 | 2.0 |
| Fig. S6D-E, first column | 200 | 12 | 0.1 | 2.0 | varies | 1.0 | varies |
| Fig. S6D-E, second column | 200 | 12 | varies | 2.0 | 2.0 | 1.0 | varies |
| Fig. S6D-E, third column | 200 | varies | 0.1 | 2.0 | 2.0 | 1.0 | varies |
| Fig. S7A-B | 600 | varies | varies | varies | varies | varies | varies |
| Fig. S8A | 600 | 8 ($Q = 64$) | 0.0 | 1.0 | 3.0 | 1.0 | 0.0 |
| Fig. S8B | 600 | 8 ($Q = 64$) | 0.0 | 1.0 | 3.0 | 1.0 | 4.0 |
| Fig. S8C | 600 | varies | varies | varies | varies | varies | varies |
| Fig. S10A-C | 600 | varies | 0.0 | 1.0 | 0.0 | 1.0 | 0.0 |
| Fig. S10D-F | 600 | varies | 0.0 | 1.0 | 0.0 | 1.0 | 4.0 |

The synaptic weights, ***u*** and ***w***, are initialized at random from a zero-mean Gaussian distribution with variance 1/*N*. The sensory input vectors ***x*** are also drawn from a zero-mean Gaussian distribution (see Sections 4.1 and 5.1), but with variance equal to 1,

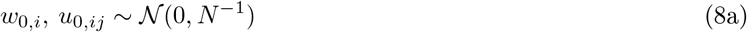

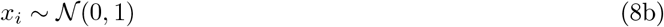

where the subscript “0” on the weights indicates that those are evaluated before learning starts. This choice of initialization ensures that, before learning, the amplitude of both the synaptic drive (*h*, and the components of ***k***) and the activity (*z*, and the components of ***y***) are independent of the circuit size (i.e., *O*(1) in *N*).

#### 2.2 Gradient descent plasticity

The circuit learns to categorize *P* sensory input vectors ***x**^s^*(*s* = 1,…, *P*), with *P* ≪ *N*. For each input vector, the target activity of the readout neuron, *z^s^*, is equal to either *z*^A^ or *z*^B^ (Section 4.1 and 5.1), which correspond to high and low activity, respectively. The weights are adjusted to minimize the loss, 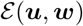, which is defined to be

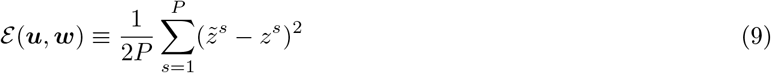

where *z^s^* is the activity of the readout neuron (Eq. 6a) in response to the sensory input ***x**^s^*. The weights are updated according to full-batch vanilla gradient descent. If the learning rates, *η_u_* and *η_w_*, are sufficiently small, the evolution of the connectivity weights can be described by the continuous-time equations (Eq. 1)

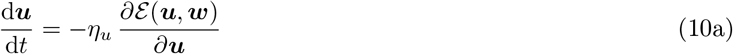

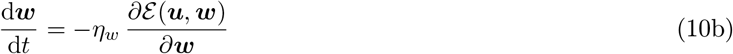

where *t* indicates learning time.

### 3 Evolution of connectivity and activity in large circuits

Our goal is to understand how learning affects activity in the intermediate layer, ***y***. We do that in two steps. In the first step, we analyze the evolution of the synaptic weights. In particular, we determine the weights after learning is complete – meaning after the loss (Eq. 9) has been minimized (Section 3.1). In the second step, we use the learned weights to determine activity (Section 3.2). We work in the large-*N* regime, which allows us to make analytic headway [32–34]. We then validate our large-*N* analysis with finite-*N* simulations (Section 3.4, Figs. S4, S5, S7, S10).

#### 3.1 Evolution of connectivity

It is convenient to make the definitions

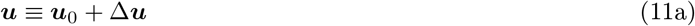

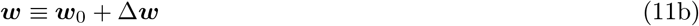

where ***u***_0_ and ***w***_0_ are the initial weights (Eq. 8), and Δ***u*** and Δ***w*** are changes in the weights induced by learning (Eq. 10). Using Eq. 10, with the loss given by Eq. 9, we see that Δ**u** and Δ**w** evolve according to

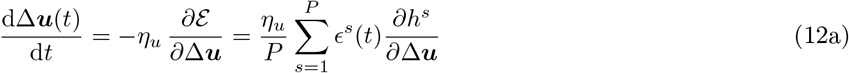

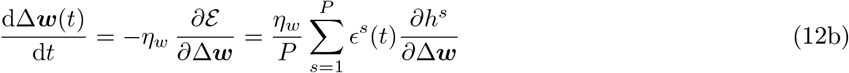

where *ϵ^s^* is proportional to the error associated with sensory input ***x**^s^*,

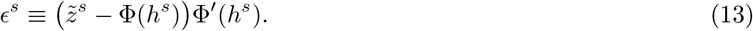

To evaluate the partial derivatives on the right-hand side of Eq. 12, we need to express *h^s^* in terms of Δ***u*** and Δ***w***. Combining Eq. 6b with 5 and 11, we have

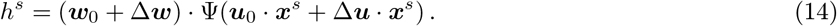

To proceed, we assume that changes in the connectivity, Δ***u*** and Δ***w***, are small. That holds in the large-*N* limit (the limit we consider here) because when each neuron receives a large number of inputs, none of them has to change very much to cause a large change in the output (we make this reasoning more quantitative in Section 3.3). Then, Taylor-expanding the nonlinear activation function Ψ in Eq. 14, and keeping only terms that are zeroth and first order in the weight changes Δ***u*** and Δ***w***, we have

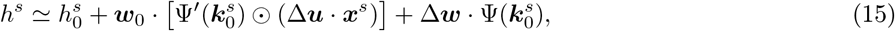

where ⊙ indicates element-wise multiplication, and we have defined

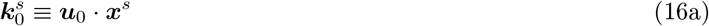

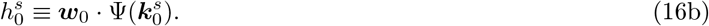

For now, we assume that the three terms in the right-hand side of Eq. 15 are of similar magnitude, and that higher-order terms in Δ***u*** and Δ***w*** are smaller, and so can be neglected. We will verify these assumptions post-hoc (Section 3.3). Inserting Eq. 15 into Eq. 12, we arrive at

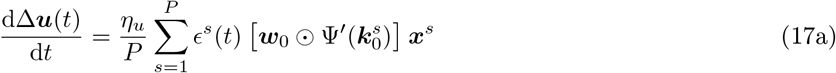

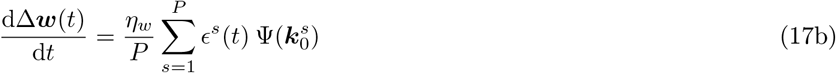

(we used the notation where two adjacent vectors correspond to an outer product; i.e. (***ab***)_*ij*_ = *a_i_b_j_*).

The only quantity on the right-hand side of Eq. 17 that depends on time is *ϵ^s^*. Consequently, we can immediately write down the solution,

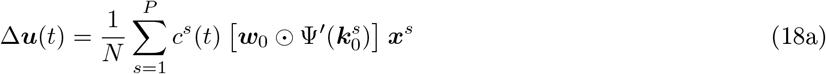

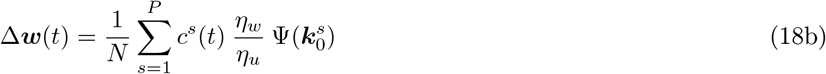

where the coefficients *c^s^* are found by solving the differential equation

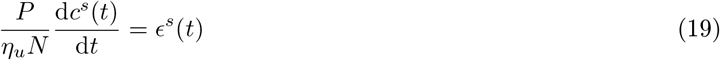

with initial conditions *c^s^*(*t* = 0) = 0. The right-hand side of Eq. 19 depends on time through the synaptic drive, *h^s^* (Eq. 13), which in turn depends on Δ***u*** and Δ***w*** through Eq. 15, and thus, via Eq. 18, on the coefficients *c*^s^(*t*). Consequently, Eq. 19 is a closed differential equation for the coefficients *c^s^*(*t*).

In the general case, Eq. 19 must be solved numerically. If, however, we are not interested in the full learning dynamics, but care only about connectivity and activity once learning has converged (*t* → ∞), we can use the fact that dynamics in Eq. 17 are guaranteed to converge to a global minimum of the error function 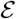 [34]. For our loss function and tasks, the minimum occurs at 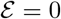. At that point, 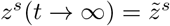; equivalently,

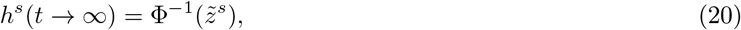

where Φ^−1^ is the inverse of the activation function of the readout neurons (which exists because Φ is a monotonically increasing function).

To find *c^s^*(*t* → ∞), we simply express *h^s^*(*t* → ∞) in terms of *c^s^*(*t* → ∞), and insert that into Eq. 20. To reduce clutter, we define (in a slight abuse of notation) *c^s^* without an argument to be its asymptotic value,

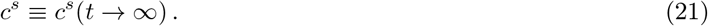

Combining Eq. 15 for *h^s^* with Eq. 18 for Δ***w*** and Δ***u***, we have

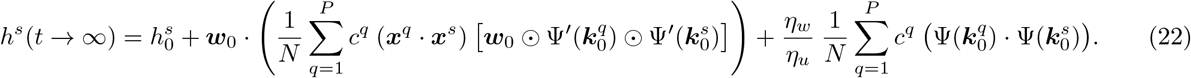

We can simplify the second term in the right-hand side by explicitly evaluating the dot product,

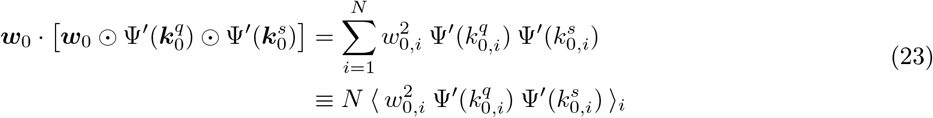

where the notation 〈. 〉_*i*_ indicates an average over the index *i*.

Since *N* is large, we can interpret population averages such as Eq. 23 as expectations over the probability distribution of *w*_0,*i*_ and 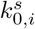. An immediate implication is that Eq. 23 simplifies,

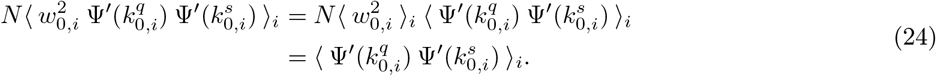

For the first equality we used the independence of *w*_0,*i*_ and *k*_0,*i*_; for the second we used the fact that the elements of ***w***_0_ are drawn from a zero-mean Gaussian with variance *N*^−1^ (Eq. 8a). We can thus rewrite Eq. 22 as

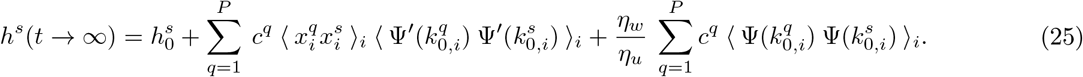

Combining this with Eq. 20, we conclude that

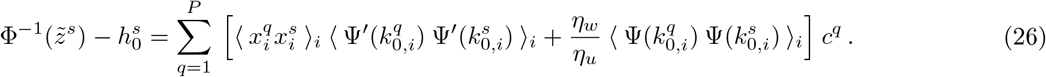

Equation 26 is a *P*-dimensional linear system of equations for the coefficients *c^s^, s* = 1,…, *P* (the term in brackets is a *P* × *P* matrix with indices *s* and *q*). For the tasks we consider (Sections 4 and 5), this system can be solved analytically, yielding a closed-form expression for the coefficients *c^s^*.

#### 3.2 Evolution of activity

It is now straightforward to determine how activity in the intermediate layer, ***y**^s^*(*t*), evolves. Inserting Eq. 18 into Eq. 5, and Taylor-expanding the nonlinear activation function Ψ to first order in Δ***u***, we arrive at

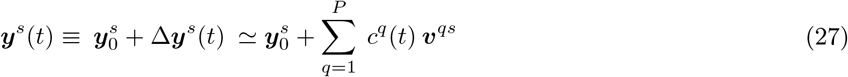

where

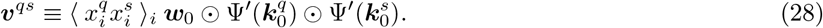

To reduce clutter, we define (following the notation in the previous section) ***y**^s^* without an argument to be its asymptotic value: ***y**^s^* ≡ ***y**^s^*(*t* → ∞). Thus, Eq. 27 becomes

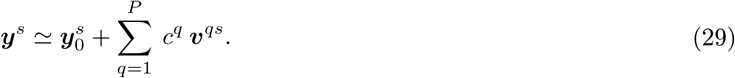

Because of the term ***w***_0_ on the right-hand side of Eq. 28, the elements of ***v***^qs^ scale as *N*^−1/2^. Thus, changes in activity are small compared to the initial activity, which is *O*(1).

In what follows, we refer to {***v**^qs^*}_*qs*_ as spanning vectors, and to the coefficients *c^q^* as the activity coordinates. We observe that all spanning vectors have a non-zero overlap with the initial readout vector ***w***_0_, as

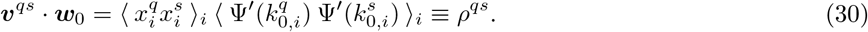

This implies that, for every spanning vector, we can write

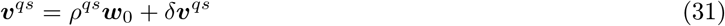

where *ρ^qs^* is given by Eq. 30 (since ***w***_0_ · ***w***_0_ = 1) and *δ**v**^qs^* is a residual component due to the nonlinearity of the activation function Ψ:

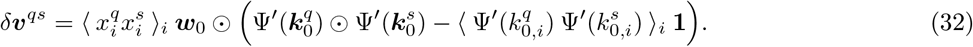

The notation **1** indicates a vector whose components are all equal to 1: **1** ≡ (1,1,…, 1).

#### 3.3 A low-order Taylor expansion is self-consistent in large circuits

To conclude our theoretical derivation, we verify that the approximations we made in Section 3.1 are valid in large circuits. Specifically, we show that the approximate expression for *h^s^*, Eq. 15 (which was derived by Taylor-expanding the nonlinear activation function Ψ), is self-consistent when N is large. As a first step, we compute the size of Δ***u*** and Δ***w***, and show that in the large-*N* limit they are small compared to ***u***_0_ and ***w***_0_, respectively. We then Taylor-expand Ψ in Eq. 14 to all orders, and show that the terms that were included in Eq. 15 (zeroth- and first-order terms in connectivity changes) are indeed the dominant ones.

Assuming that the term in brackets in Eq. 26, when viewed as a *P* × *P* matrix, is invertible (which is generically the case when *P* ≪ *N*), it follows that, with respect to *N*

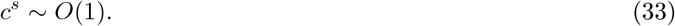

This result applies to the asymptotic (*t* → ∞) value of *c^s^* (Eq. 21). We assume, though, that the learning process is smooth enough that *c^s^*(*t*) remains at most *O*(1) for all *t*. Under this assumption, the results we derive in this section are valid at any point during learning.

Using Eq. 33, along with the fact that *w*_0,*i*_ ~ *O*(*N*^−1/2^) while all other variables are *O*(1), we see from Eq. 18 that

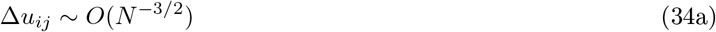

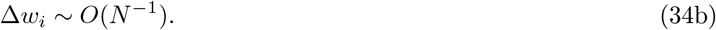

When *N* is large, both are small compared to the initial weights ***u***_0_ and ***w***_0_, whose elements are *O*(*N*^−1/2^) (Eq. 8).

Equation 34 suggests that a low-order Taylor expansion is self-consistent, but it is not proof. We thus turn directly to Eq. 14. The three terms in the right-hand side of Eq. 15 are re-written in Eq. 25, and it is clear from that expression that they are all *O*(1). To determine the size of the higher order terms, we need the complete Taylor expansion of Eq. 14. That is given by

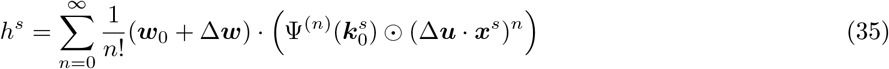

where Ψ^(*n*)^ is the *n*^th^ derivative of Ψ, and the exponentiation in (Δ***u*** · ***x**^s^*)^*n*^ is taken element-wise. The higher order terms (i.e., the terms not included in Eq. 15) are

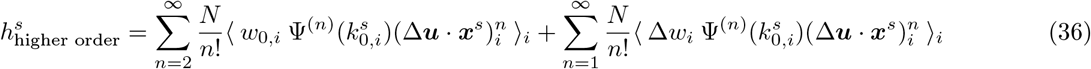

where we have replaced dot products with averages over indices. Using Eq. 18a, and taking into account the fact that *w*_0,*i*_ and *k*_0,*i*_ are independent, we observe that

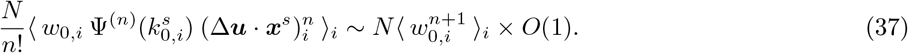

Similarly, this time using both Eq. 18a and b, we have

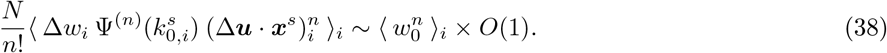

Inserting these into Eq. 36 then gives us

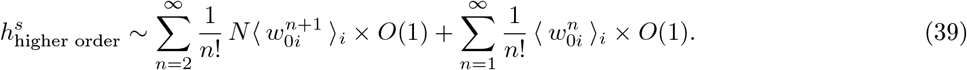

Finally, using the fact that the *w*_0,*i*_ are drawn independently from a zero-mean Gaussian with variance *N*^−1^ (Eq. 8a), we see that 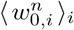 is proportional to *N*^−n/2^ when *n* is even and N^−(n+1)/2^ when *n* is odd. Consequently, the largest term in the expression for 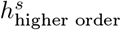 is proportional to *N*^−1^. The higher order terms can, therefore, be neglected in the large *N* limit.

#### 3.4 Evolution of activity in finite-size networks

The equations that describe the evolution of connectivity and activity that were derived in Sections 3.1 and 3.2 are accurate if two assumptions are satisfied: (i) the circuit is very large (*N* ≫ 1), and (ii) the synaptic weights are initialized to be *O*(*N*^−1/2^) (Eq. 8a), which guarantees that synaptic drives and activity neither vanish nor explode at initialization. Both assumptions are reasonable for brain circuits, and correspond to rather standard modeling choices in theoretical neuroscience.

In this work, we use the analytical expressions derived for large N to describe activity evolution in finite-size networks. This is a crude approximation, as dealing with finite N would require, in principle, integrating corrective terms into our equations [72]. How accurate is this approximation? Several machine-learning studies have investigated this question across tasks, architectures, and loss functions [32, 35, 56, 63, 73]. Because of the Taylor expansions used in Sections 3.1 and 3.2, for fixed *N*, good accuracy is expected when the amplitude of activity changes is small. Via Eq. 29, we see that the latter increases with the number of sensory input vectors *P*, implying that good accuracy is expected when *P* is small. For fixed *P*, furthermore, the amplitude of activity changes increases with correlations among sensory inputs (Eq. 28), implying that good accuracy is expected when sensory input correlations are small. As detailed in Sections 4.1 and 5.1, sensory input correlations are smaller in the simple than in the context-dependent task, which implies that accuracy in the former task is expected to be higher than in the latter. The amplitude of activity changes also depends on the amplitude of activity coordinates *c^s^* (Eq. 29). We show in Sections 4.2 and 5.2 that activity coordinates are usually smaller in the simple than in the complex task, which again implies that accuracy in the former task is expected to be higher than in the latter. Overall, those arguments suggest that good accuracy is expected when the task is easy, and thus the training loss converges to zero very quickly [35]. Finally, we expect accuracy to depend on properties of the activation function Ψ, with accuracy increasing as Ψ becomes more linear in its effective activation range.

In Figs. S4, S5, S7 and S10, we evaluate accuracy by performing a systematic comparison between approximate analytical expressions (large *N*) and circuit simulations (finite *N*). We find good agreement for the full range of parameters considered in the study. Specifically, the theory correctly predicts qualitatively, and in some cases also quantitatively, the behaviour of all activity measures discussed in the main text. As expected, the agreement is stronger in the simple (Figs. S4 and S5) than in the context-dependent task (Figs. S7 and S10).

### 4 Simple categorization task

#### 4.1 Simple task: task definition

We first consider a simple categorization task. Each stimulus is represented by an input pattern ***μ**^S^*, with *S* = 1,…, *Q* where *Q* is the total number of stimuli. The ***μ**^S^* are random vectors whose entries are drawn independently from a zero-mean, unit-variance Gaussian distribution. Every sensory input vector ***x**^s^* corresponds to a stimulus,

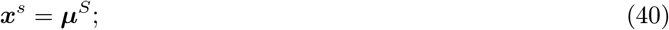

consequently, the number of sensory input vectors, *P*, is equal to *Q* (the upper-case notation *S* is used for consistency withthe context-dependent task; see Section 5.1). To leading order in *N*, sensory input vectors are thus orthonormal,

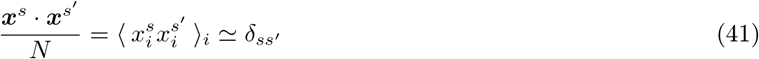

where *δ_ss′_* is the Kronecker delta.

Each stimulus is associated with one among the two mutually-exclusive categories A and B: the first half of stimuli is associated with A, the second half with B. The target value 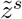 for the readout neuron is thus equal to *z*^A^ for the first half of sensory inputs and *z^B^* for the second half. Since sensory input vectors are approximately orthogonal to each other, they are also linearly separable.

Our goal is to derive explicit expressions for the quantities analyzed in the main text: category selectivity (defined in Eq. 56 below), and category correlation (defined in Eq. 71 below). Both quantities depend on activity in the intermediate layer, ***y**^s^*, after learning, which is given in Eq. 29. In the next section, we then write down an explicit expression for ***y**^s^*; after that, we compute category selectivity (Section 4.3) and category correlation (Section 4.4). Further mathematical details are discussed in Sections 4.5, 4.6 and 4.7; a generalization of the current task is discussed and analyzed in Section 4.8.

#### 4.2 Simple task: computing activity

Examining Eq. 29, we see that to compute the activity in the intermediate layer, ***y**^s^*, we need the asymptotic activity coordinates, *c^q^*, and the spanning vectors, ***v**^qs^*. We start with the coordinates. To compute them, we solve the linear system of equations given in Eq. 26. Using Eq. 41, that system of equations becomes

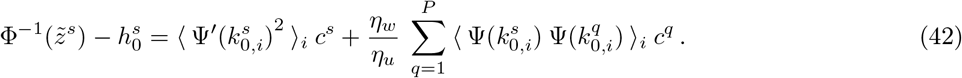

As a first step, we simplify the averages in the right-hand side. The law of large number guarantees that, when *N* is large, the elements of the synaptic drive, 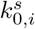, are independently drawn from a Gaussian distribution. The statistics of this distribution are given by

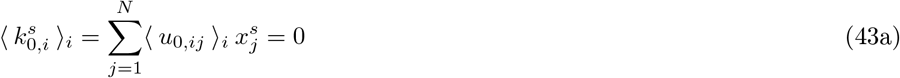

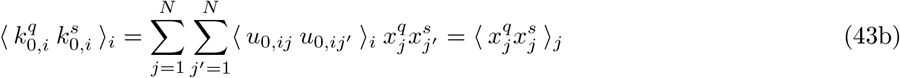

where we have used the fact that, because of Eq. 8a, 〈*u*_0,*ij*_ *u*_0,*ij*′_〉_*i*_ = *δ*_*jj*〉_/*N*. Equation 43, combined with Eq. 41, implies that the 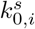 have zero mean and unit variance, and are uncorrelated across stimuli. In addition, because the statistics of 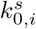 are independent of s, averages over i of any function of 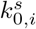 are independent of *s*.

Using these observations, Eq. 42 can be written as

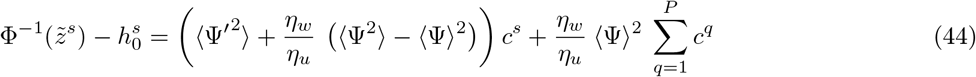

here we used the short-hand notation 〈*F*〉 to indicatethe average of a function *F* whose argument is drawn from a zero-mean, unit-variance Gaussian distribution. I.e.,

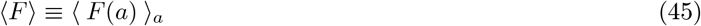

where *a* is a zero-mean, unit-variance Gaussian variable. This average can be computed via numerical integration, as detailed in Section 6.3 (Eq. 184).

The left-hand side of Eq. 44 consists of two terms: the target 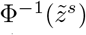, which is fixed by the task, and 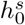 (representing the synaptic drive of the readout neuron at initialization), which fluctuates across model realizations. The presence of the latter term indicates that connectivity and activity changes are not fully self-averaging; they are rather tuned to compensate for the initial state of the readout neuron. Here we seek to analyze the *average* behaviour of the model, and so we drop the second, variable term. This approximation is discussed in detail in Section 4.7.

With the variable terms neglected, the left-hand side of Eq. 44 can take only two values: Φ^−1^(z^A^) and Φ^−1^(z^B^). Combined with the symmetry of the right-hand side, this implies that the coordinates *c^s^* themselves can take only two values. Specifically, we have

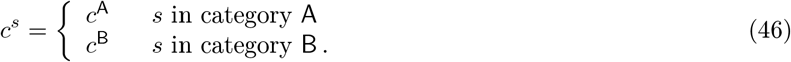

The category-dependent coordinates, *c*^A^ and *c*^B^, are determined by the two-dimensional linear system of equations

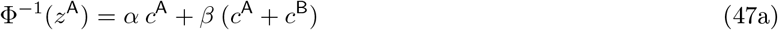

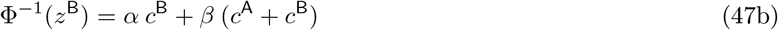

where the scalars *α* and *β* are defined as

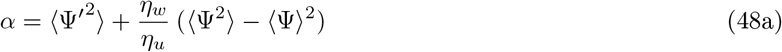

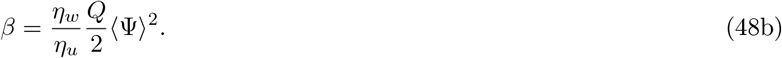

This system is easily solved, yielding

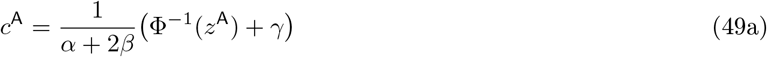

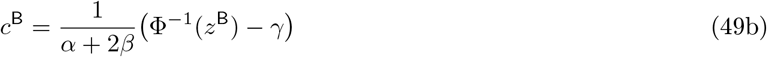

where we have defined the shift

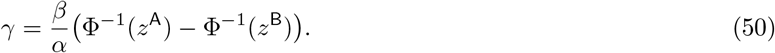

Note that *γ* is positive, as *α, β* > 0 and Φ^−1^(*z*^A^) > Φ^−1^(*z*^B^), which in turn indicates that *c*^A^ > *c*^B^.

To conclude the derivation of activity, we evaluate the spanning vectors, ***v**^qs^* (Eq. 28). Because the sensory inputs ***x**^s^* are orthogonal (Eq. 41), spanning vectors with *q* ≠ *s* vanish. Consequently, the activity, ***y**^s^* (Eq. 29), reads

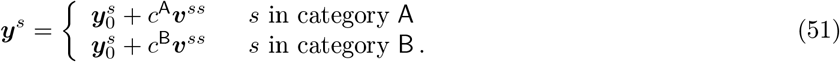

Using Eq. 31, we can rewrite this as

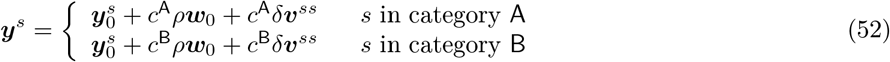

where we used Eq. 30 to define

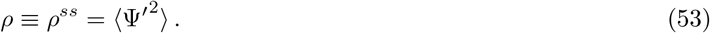

Equation 52 indicates that activity consists of three components. The first one coincide with initial activity, 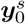, which for this task is fully unstructured. The second one is a shared component along ***w***_0_ (whose strength is category-dependent, as it is given by *c*^A^ or *c*^B^). The third one is a non-shared component along the residuals *δ**v**^ss^*, which represent the components of the spanning vectors that are perpendicular to the initial readout ***w***_0_. For the current task, the latter component is orthogonal across activity vectors, implying that activity vectors only overlap along ***w***_0_. To leading order in *N*, in fact

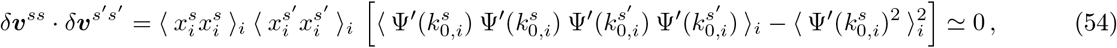

which follows because 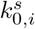 and 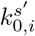 are uncorrelated.

We observe that Eq. 52 is similar, but not identical to the expression that we used in the main text to describe activity evolution (Eq. 3). By setting ***d*** = *ρ**w***_0_, that reads

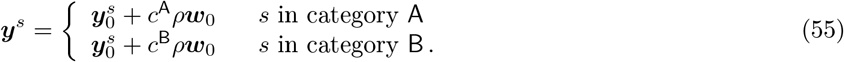

Comparing Eq. 52 with 55, we see that the residuals *δ**v**^ss^* were neglected in the main text. This could be done because, for the current task (but not for the context-dependent one, see Section 5.2), residuals are all orthogonal to each other (Eq. 54). As such, they do not add novel structure to activity, and do not significantly contribute to activity measures. This is showed and justified, in detail, in the next sections.

#### 4.3 Simple task: category selectivity

In this section, we evaluate the category selectivity of neurons in the intermediate layer (Figs. 2B, F and J). For each neuron *i*, we evaluate the standard selectivity index [4], defined in Eq. 2. We repeat that definition here for convenience,

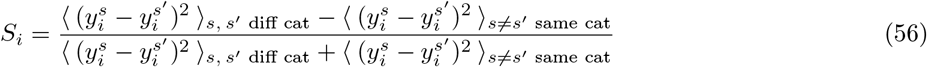

where the notation 〈·〉_*s, s′*_ denotes an average over sensory input pairs associated either with different, or the same, category. To evaluate this expression, we assume that the number of stimuli, *Q* = *P*, is moderately large (1 ≪ *Q* ≪ *N*). We show that, under this assumption, the category selectivity index for each neuron, which is approximately zero at *t* = 0, becomes positive over learning.

We start with

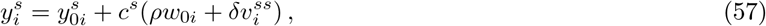

which follows from Eq. 52. The first term of the right-hand side is *O*(1), while both terms in parentheses are *O*(*N*^−1/2^). Thus, when evaluating the denominator in Eq. 56, to lowest non-vanishing order in *N* we can replace *y_i_* with *y*_0*i*_. Doing that, and expanding the square, we have

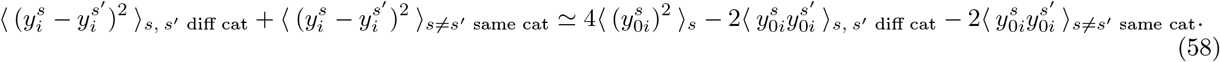

Noting that 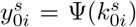, and using Eq. 43, we see that the second two averages in the above equation are both equal to 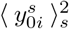. Consequently,

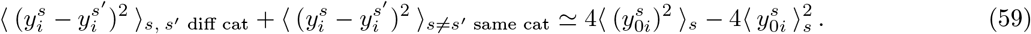

Strictly speaking, this step is accurate only in the large-*Q* limit, but is a good approximation even for moderate *Q*. Since *Q* is moderately large, we can further approximate this as

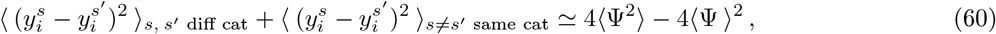

where averages can be computed as described in Section 6.3.

For the numerator of Eq. 56, the minus sign causes the 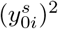 terms to cancel, so we have

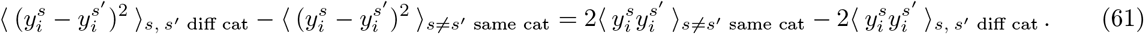

Using Eq. 57, we have (for *s* = *s′*)

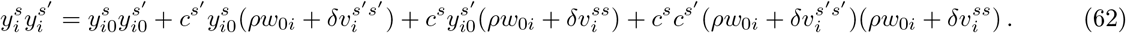

Apart from the first term, and the term proportional to 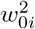, all terms in the right-hand side have essentially random signs. Neglecting those for a moment, we obtain

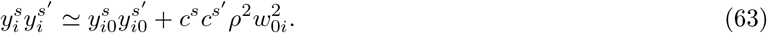

Inserting this into Eq. 61, using the fact that 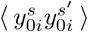 is independent of *s* and *s′*, and performing a small amount of algebra, we arrive at

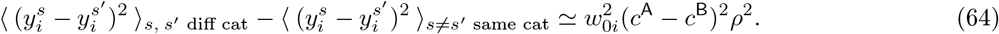

Combining this with Eq. 60, and using Eq. 53 for *ρ*, we arrive at

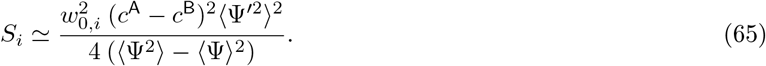

We conclude that single-neuron selectivity vanishes at *t* = 0 (when *c*^A^ = *c*^B^ = 0), and is positive at the end of learning. Furthermore, for each neuron, the magnitude of selectivity is determined by the magnitude of *w*_0,*i*_, which measures initial connectivity with the readout neuron. As a result, neurons with large initial connectivity develop large selectivity values (Figs. 5B-C).

Because of the factor 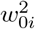, the right-hand side of Eq. 65 is *O*(*N*^−1^). To derive Eq. 65, we neglected terms in the numerator that have random sign and thus contribute as noise. The dominant random terms are *O*(1) in *N*, but *O*(*Q*^−1^) in *Q*. This implies that, in simulated circuits with finite *Q*, random deviations from Eq. 65 occur. For example, Fig. 2B shows that selectivity values at *t* = 0 are small but non-zero; Figs. 5B-C, instead, shows that the values of *S_i_* and 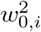, are not perfectly correlated across the population.

We can, finally, average Eq. 65 over neurons, yielding

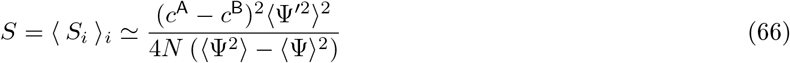

where neglected random terms are now *O*(*N*^−1/2^*Q*^−1^). In Figs. S4 and S5, we compare this approximate analytical expression for average category selectivity with values measured in finite-size circuits, and find good agreement between the two.

##### Category clustering

Our derivation of the average category selectivity, Eq. 66, was based on several assumptions: we assumed that the number of stimuli, *Q*, was large, and that terms with random signs could be neglected. A different, but related, activity measure is given by category clustering [42, 74]. That is defined as

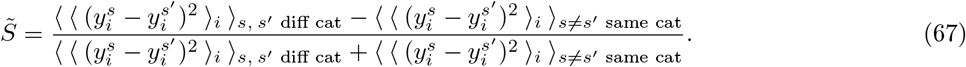

This measure is positive if activity vectors elicited by within-category stimuli are more similar, in norm, than activity vectors elicited by across-category stimuli – and negative otherwise. In contrast to average category selectivity, category clustering can be evaluated straightforwardly, and for any value of *Q*. We show this in the following.

By using the statistical homogeneity of activity vectors, we can rewrite

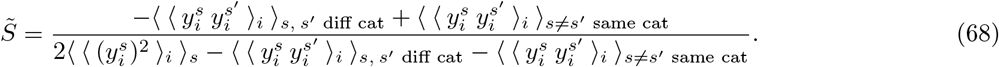

Expressions in the form of 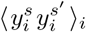 are evaluated in Section 4.5; the derivation involves lengthy, but straightforward algebra. Using those results (Eqs. 90 and 95), we have:

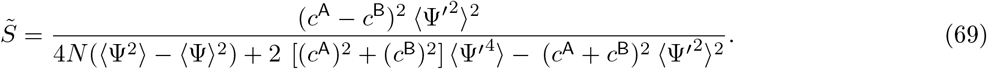

To the leading order in *N*, we obtain

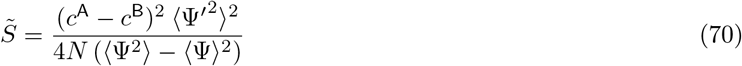

which is identical to the expression obtained for average category selectivity evaluated with *Q* large (Eq. 66).

To better understand the relationship between selectivity and clustering, we observe that clustering coincide with the average selectivity, *S* = 〈*S_i_*〈_*i*_, if the average over the numerator and the denominator of *S_i_* (Eq. 56) is factorized. In general, the numerator and the denominator of Siare correlated, and the average cannot be factorized. We have however shown that, in the limit where both *Q* and *N* are large, *S_i_* can be approximated by an expression where the denominator is independent of *i* (Eq. 65). In that regime, the average can be factorized; average category selectivity *S* and category clustering 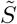 thus take very similar values, as quantified by Eqs. 66 and 70. We conclude that, for our activity expressions, average category selectivity and category clustering are expected to behave similarly when both *Q* and *N* are large. A detailed comparison between average selectivity and clustering within data from simulated circuits is provided in Figs. S4 and S5.

#### 4.4 Simple task: category correlation

To quantify how the population as a whole responds to the two categories, we evaluate category correlation. This quantity, denoted *C*, is given by the average Pearson correlation coefficient of activity in response to stimuli associated with different categories. We have:

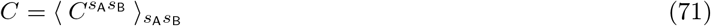

where *s*_A_ and *s*_B_ are indices that denote sensory inputs associated, respectively, with category `A` and `B`. The Pearson correlation *C*^*s*_A_*s*_B_^ is given by

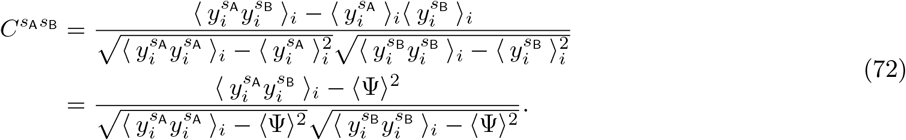

To go from the first to the second line, we used the fact that, for each sensory input,

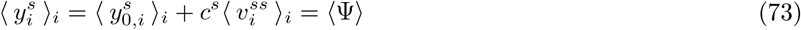

where the second equality follows from 〈*w*_0,*i*_〉_*i*_ = 0 (Eq. 8a), which in turns implies that 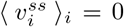 (Eq. 28). Pearson correlation coefficients are displayed in the correlation matrices of Figs. 2C, G and K.

As we show in Section 4.5, in the large-*N* limit, 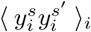 only depends on the category *s* and *s′* are in. This makes the average over *s*_A_ and *s*_B_ in Eq. 71 trivial. Using Eqs. 90 and 95, we arrive at

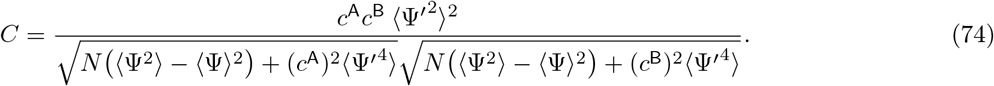

In Figs. S4 and S5, we compare this approximate analytical expression with values measured in finite-size circuits, and find good agreement between the two. We can further simplify Eq. 74 by Taylor-expanding in *N*. To leading order, we obtain

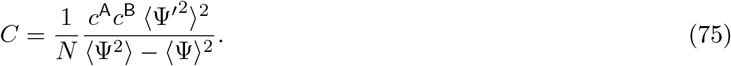

Before learning, *c*^A^ = *c*^B^ = 0, and so correlation vanishes. After learning, *C* is non-zero, and its sign is given by the sign of the product *c*^A^*c*^B^. This has a simple geometric explanation: after mean subtraction, activity vectors associated with opposite categories only overlap along the direction spanned by the initial readout vector ***w***_0_. The coordinates of vectors associated with category `A` and `B` along this direction are proportional, respectively, to *c*^A^ and *c*^B^ (Eq. 55). When *c*^A^ and *c*^B^ have opposite sign, activity vectors acquire opposite components along **w**_0_, which generates negative category correlation. When c^A^and c^B^have identical sign, instead, activity vectors acquire aligned components, which generates positive category correlation.

To determine how the product *c*^A^*c*^B^ depends on parameters, we use Eq. 49 for *c*^A^ and *c*^B^ to write

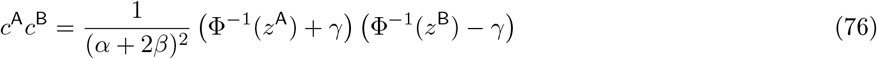

where the (positive) scalars *α* and *β* are defined in Eq. 48, and *γ* in Eq. 50. Consequently, the sign of *c*^A^*c*^B^, and thus, the sign of the category correlation *C*, depends on the value of the target synaptic drives Φ^−1^(*z*^A^) and Φ^−1^(*z*^B^) (Fig. S2F), as well as on *γ*.

In particular, when Φ^−1^(*z*^A^) and Φ^−1^(*z*^B^) have opposite sign, Eq. 76 can only be negative, and thus category correlation can only be negative. When Φ^−1^(*z*^A^) and Φ^−1^(*z*^B^) have identical sign, Eq. 76 can be either negative or positive, depending on the value of the shift *γ*, and thus category correlation can be either negative or positive. For fixed target values *z*^A^ and *z*^B^, the relative sign of Φ^−1^(*z*^A^) and Φ^−1^(*z*^B^) depends on the shape of the activation function of the readout neuron Φ. In the example given in Fig. S2F, we show that the relative sign of Φ^−1^(*z*^A^) and Φ^−1^(*z*^B^) can be modified by changing the threshold of Φ. More in general, changing both the gain and threshold of Φ can change the sign and magnitude of category correlation (Fig. 4A).

What controls the value of the shift *γ* (and, thus, the sign of correlation when Φ^−1^(*z*^A^) and Φ^−1^(*z*^B^) have identical sign)? Combining Eq. 50 for *γ* with Eq. 48 for *α* and *β*, we have

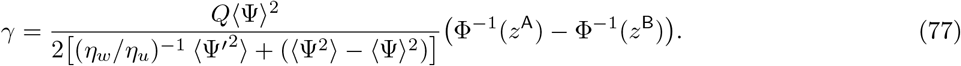

Recall that Φ^−1^(*z*^A^) – Φ^−1^(*z*^B^) is always positive. We observe that *γ* depends on the learning rate ratio *¾_w_*/*η_u_*: increasing this ratio increases the value of *γ* and thus, via Eq. 76, favors negative correlation (Fig. 4C). It also depends on the number of stimuli, *Q*: increasing *Q* increases the value of *γ*, and thus also favors negative correlation (Fig. 4D). Finally, *γ* depends on the activation function of neurons in the intermediate layer, Ψ, through nonlinear population averages; by computing those averages, we find that decreasing the gain and threshold of Ψ favors negative correlation (Fig. 4B).

##### Alternative definition

For completeness, we observe that an alternative way of quantifying category correlation consists of averaging activity over stimuli first (Fig. 2D, H and L), and then computing the Person correlation coefficient between averaged responses. The correlation values obtained via this procedure are displayed in the legend of Figs. 2D, H and L. This alternative definition yields qualitatively identical results to Eq. 71; we show this below.

We start by defining the category-averaged activity

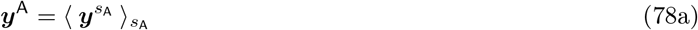

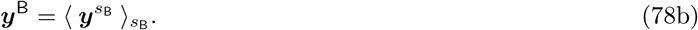

We then define category correlation as

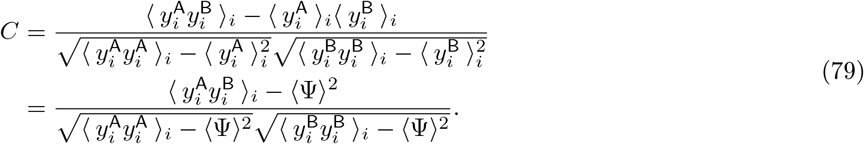

Then, using Eqs. 90 and 95 from Section 4.5, we have

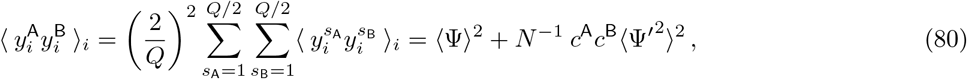

while

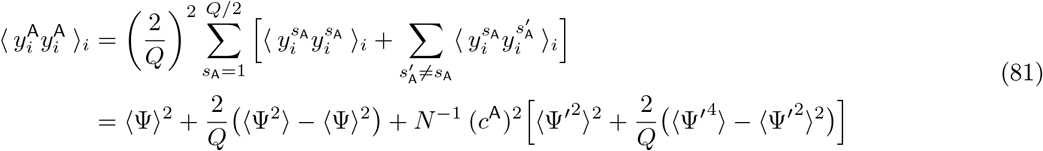

and similarly for 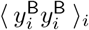, by replacing *c*^A^ with *c*^B^. Inserting this into Eq. 79, we arrive at

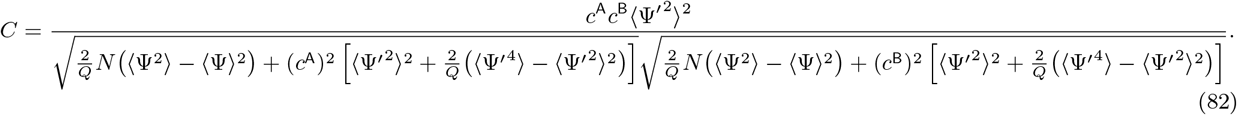

Although the denominator of this expression is different from Eq. 74, the numerator is identical. As the denominators in both expressions are positive, the qualitative behaviour of Eq. 82 is identical to Eq. 74. Furthermore, to leading order in *N*, we obtain

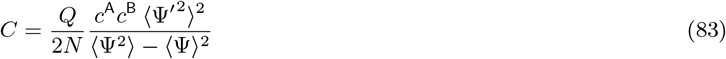

which is proportional to Eq. 75, with constant of proportionality equal to *Q*/2.

#### 4.5 Simple task: computing normalized dot products

We now compute the normalized dot products among pairs of activity vectors; namely

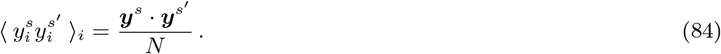

Those were used above to derive the behaviour of category clustering (Section 4.3) and correlations (Section 4.4).

The dot product takes different values depending on whether or not sensory inputs *s* and *s′* coincide. We start with the former,

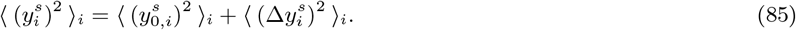

We used the fact that the cross-term 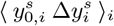 vanishes on average,

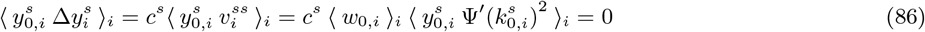

where we used Eq. 51 for the first equality, Eq. 28 for the second, and Eq. 8a for the third. By definition,

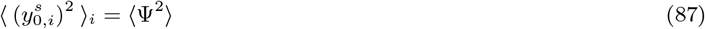

while

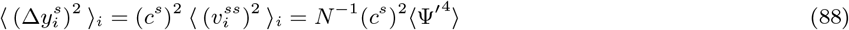

where we have used the fact that, from Eq. 28

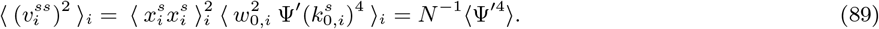

Putting these results together, we have

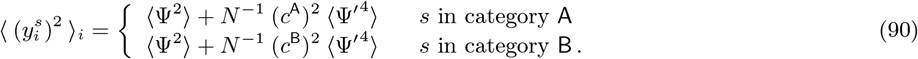

Note that activity vectors associated with different categories are characterized by different norms (unless coordinates are fine-tuned to be symmetric: *c*^A^ = –*c*^B^, which occurs when Φ^−1^(*z*^A^) = –Φ^−1^(*z*^B^), as in Figs. 2E-H). Asymmetry of activity in response to different categories is discussed in detail in Section 4.6.

For dot products among different activity vectors, we have

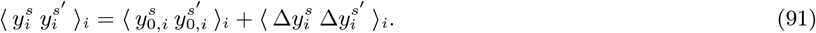

with *s* = *s′*. In this case,

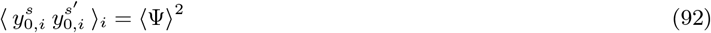

while

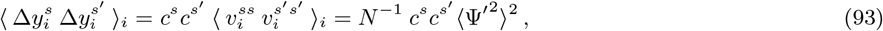

which comes from

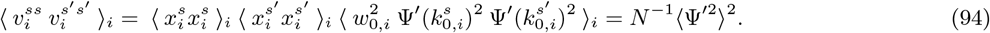

Putting this together, we arrive at

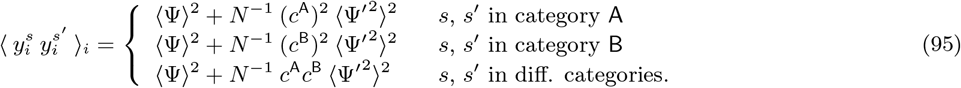

Equation 95 has a simple geometric interpretation. The first term in the right-hand side, 〈Ψ〉^2^, is generated by the overlap between the activity vectors along the direction spanned by the unit vector 1. This component is due to the activation function Ψ being positive, and is approximately constant over learning. The second term on the right-hand side emerges over learning. This arises because activity vectors become aligned, via the spanning vectors (Eq. 30), along the direction spanned by the initial readout vector ***w***_0_. Note that the components of activity that are aligned with the residual directions *δ**v**^ss^* (Eq. 52) do not contribute to the dot product. This can be verified by computing the dot product directly from Eq. 55, where residuals are neglected, and observing that the same result is obtained. This was expected, as we have showed in Eq. 54 that, for the current task task, residuals are orthogonal to each other.

#### 4.6 Asymmetry in category response

In Fig. 2L in the main text, activity in response to category `A` and `B` is asymmetric: the number of neurons that respond more strongly to category `A` is significantly larger than the number that respond more strongly to `B`. Furthermore, the mean and variance of activity across the population is larger in response to `A` than to `B`. Such asymmetry is not present at *t* = 0 (Fig. 2D), and is thus a consequence of learning. Asymmetry has been reported in experimental data as well [18], where it was referred to as *biased category representations*. Here we discuss in detail why and how response asymmetry arises in the model. We show that asymmetry is controlled by the value of the target readout activity, *z*^A^ and *z*^B^, and also by the shape of the activation functions of the intermediate and readout layer, Ψ and Φ.

Figure 2L displays activity in response to category `A` and `B` averaged over stimuli; those are denoted, respectively, by ***y***^A^ and ***y***^B^ (Eq. 78). We start deriving an explicit expression for ***y***^A^, from which the mean and variance across the population can be computed. Since initial activity is symmetric, we focus on the part of activity that is induced by learning. Combining Eq. 51 with 28, we have

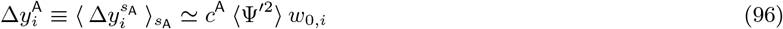

where the last approximate equality follows if *Q* is sufficiently large. The variance across the population is, therefore, given by

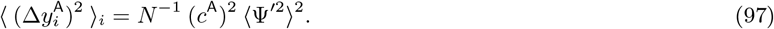

For the variance across the population in response to category `B`, we simply replace *c*^A^ with *c*^B^.

Consequently, the variances in response to categories `A` and `B` are identical only if (*c*^A^)^2^ = (*c*^B^)^2^. From Eq. 49, we see that this happens only if Φ^−1^(*z*^A^) = –Φ^−1^(*z*^B^), which yields *c*^A^ = –*c*^B^. Figure 2H shows a circuit where the activation function of the readout neuron, Φ, was chosen to satisfy this relationship. In general, however, the two variances differ, and can have either (*c*^A^)^2^ > (*c*^B^)^2^ (the variance in response to `A` is lather than to `B`), or (*c*^A^)^2^ < (*c*^B^)^2^ (the opposite). Figure 2L corresponds to the first scenario, (*c*^A^)^2^ > (*c*^B^)^2^; this was achieved by setting Φ^−1^(*z*^A^) > Φ^−1^(*z*^B^) > 0, which yielded *c*^A^ > *c*^B^ > 0. Figures S2A-B correspond to the second scenario, (*c*^A^)^2^ < (*c*^B^)^2^; this was achieved by setting Φ^−1^(*z*^B^) < Φ^−1^(*z*^A^) < 0, which yielded *c*^B^ < *c*^A^ < 0. Note that in both cases, *c*^A^ > *c*^B^, as it must be (Eq. 49).

In Fig. 2L, activity in response to category `A` is not only characterized by larger variance, but also larger mean. This observation does not emerge immediately from our analysis, since our equations predict that the mean of activity changes vanishes both in response to A and B: from Eq. 8, we see that in response to category `A`,

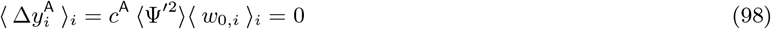

and similarly for category `B`. To understand how Eq. 98 can be reconciled with Fig. 2L, recall that the equations we use for activity changes (Eq. 29) provide a linearized estimate of activity changes, which is strictly valid only in infinitely wide networks. In finite width networks, a non-zero mean response can emerge from higher-order terms in the expansion of Eq. 27. The leading higher-order terms of this expansion are quadratic, implying that the behaviour of the mean is controlled by the second-order derivative of the activation function of neurons in the intermediate layer, Ψ′. When the threshold of Ψis positive (so that activity is initialized close to the lower bound of Ψ), the second-order derivative Ψ′ is positive on average. Combined with (*c*^A^)^2^ > (*c*^B^)^2^, this implies that the mean of activity in response to category `A` is larger than to `B`; this case is illustrated in Fig. 2L. When the threshold of Ψ is negative (so that activity is initialized close to the upper bound of Ψ), the second-order derivative Ψ″ is negative on average. Combined with (*c*^A^)^2^ > (*c*^B^)^2^, this implies that the mean of activity in response to category A is smaller than to B; this case is illustrated in Figs. S2C-D.

Finally, Eq. 98 suggest that non-vanishing mean activity could also be obtained if the initial readout weights *w*_o,*i*_ have a non-zero mean. This is likely to be verified in the brain, where intra-area connectivity is mainly excitatory. We leave the incorporation of non-zero mean connectivity, along with Dale’s law, to future investigations.

#### 4.7 Characterizing variability

In Section 4.2, when computing the value of activity coordinates *c^s^*, we neglected the second terms within the left-hand side of Eq. 42; because of this, the coordinates took on only two values, namely *c*^A^ and *c*^B^ (Eq. 46). The neglected terms do not self-average, and thus fluctuate at random across model realizations. Had we included these variable terms, Eq. 46 would have read

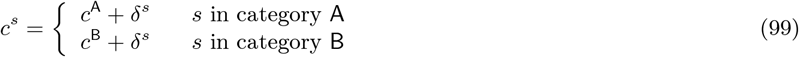

where the *δ^s^* obey the linear system of equations

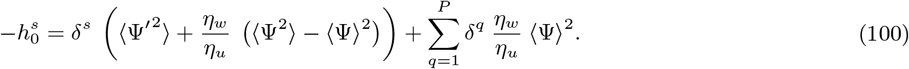

Here, we further characterize the behaviour of the neglected terms *δ^s^*. For simplicity, we consider the case in which plasticity in the readout weights is much slower than plasticity in the input connectivity (*η_w_* ≪ *η_u_*). In that regime, Eq. 100 greatly simplifies, and we obtain

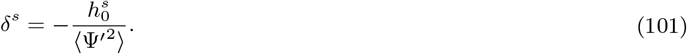

There are two sources of random fluctuations in 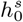: different realizations of the circuit (via different initializations of the intermediate and readout connectivity, ***u*** and ***w***), and different sensory inputs. In the following, we show that these two sources of variability can be decomposed, and one can write

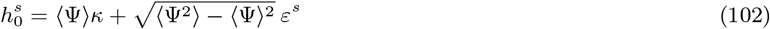

where *κ* and *ε^s^* are zero-mean, unit-variance Gaussian variables. For a given circuit realization, the value of *κ* is fixed, while the value of *ε^s^* fluctuates across different sensory inputs. Combining Eq. 102 with 99, we conclude that two different forms of variability (one that is frozen for a given circuit realization, represented by *κ*, and one that is not, represented by *ε^s^*) impact activity coordinates *c*^A^ and *c*^B^; the absolute and relative amplitude of the two contributions is controlled by the shape of the activation function Ψ. Such factorization of variability is illustrated, for an example simulated circuit, in Fig. S2E.

To derive Eq. 102, we consider a given circuit realization, and assume that the number of stimuli *Q* is sufficiently large, so that averages over stimuli approximately self-average. We start from Eq. 6b, and compute the mean of of 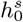 over sensory inputs, which yields

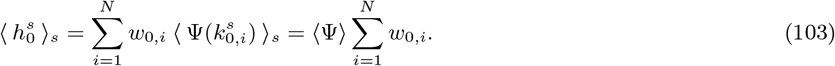

By defining 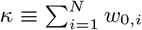, the first term in the right-hand side of Eq. 102 follows. We then compute the variance of of 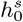 over sensory inputs. By using:

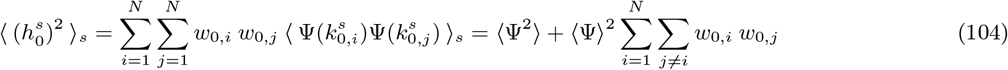

and, from Eq. 103

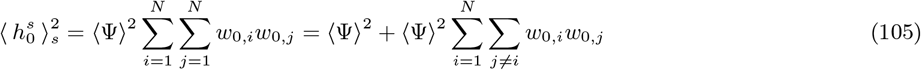

we conclude that:

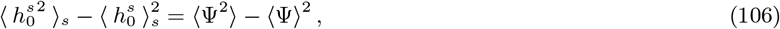

from which the second term in the right-hand side of Eq. 102 follows.

Equations 66 and 74 indicate that activity measures such as category selectivity and correlation depend on the value of activity coordinates *c*^A^ and *c*^B^. As coordinates are variable (Eq. 99), activity measures are variable as well. Importantly, activity measures involve averages over sensory inputs (see Eqs. 56 and 71). This implies that the two forms of variability described by Eq. 102 are expected to contribute in different ways: variability originating from the second term (which fluctuates across stimuli, and thus can be averaged out) is expected to be small, while variability originating from the first term (which is fixed for each circuit realization) is expected to be large.

Variability in simulated circuits is quantified in Figs. S4–S5, where it is represented as error bars. Figures S4A and S5A show that variability in *c*^A^ and *c*^B^ is modulated by properties of the activation function Ψ(third column); this is in agreement with Eq. 102, which indicates that the magnitude of variability is Ψ-dependent. Figures S4B-C and S5B-C show, furthermore, that variability in correlation is typically much larger than in average selectivity. This can be explained by observing that average selectivity (Eq. 66) only depends on the difference between *c*^A^ and *c*^B^, so variability originating from the first, frozen term of Eq. 102 is expected to cancel; this is not the case for correlation (Eq. 74), for which the cancellation does not occur.

#### 4.8 Simple categorization task with structured inputs and heterogeneity

The circuit and task we considered so far are characterized by several simplifying modelling assumptions, which allowed us to analyze activity evolution in great detail and develop useful analytical intuition. One important assumption is that sensory input vectors corresponding to different stimuli are orthogonal to each other. This choice was motivated by two observations: first, in many tasks from the experimental literature, sensory stimuli are taken to be very different from each other, and thus sensory inputs are expected to be uncorrelated [2, 6, 8]; second, in tasks where sensory stimuli obey a continuous statistical structure [4], pre-processing from sensory brain regions [75] is expected to decorrelate, at least partially, inputs to higher-level associative areas. A second important assumption is that neurons in the intermediate layer are statistically homogeneous, as they receive statistically identical inputs and are characterized by the same nonlinearity Ψ.

For some tasks and brain regions, those two assumptions might be inaccurate. For example, data collected during passive conditions [76] indicate that some LIP neurons [4, 6, 18] display weak, but significant direction tuning, which might be due to structured sensory inputs. Furthermore, activity profiles are heterogeneous, with different neurons characterized by different baseline activity levels. To investigate whether our findings extrapolate beyond our two simplifying hypotheses, here we construct a more biologically-grounded model, and use simulations to systematically investigate activity evolution in the resulting circuit.

To begin with, we use sensory input vectors characterized by a continuous statistical structure, which implies continuous tuning in the intermediate layer activity prior to learning. We set

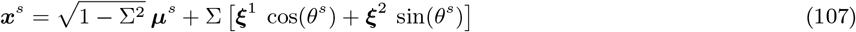

where Σ is a scalar that measures the fraction of inputs variance that is continuous. We fixed Σ = 1/3. Like ***μ**^s^*, entries of the vectors *ξ*^1^ and *ξ*^2^ are generated at random from a zero-mean, unit-variance Gaussian distribution. We furthermore set

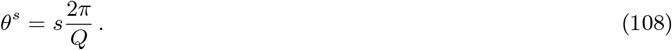

With this choice, when *s* ≠ *s′*, we have 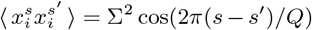, so stimuli with similar values of *s* are more strongly correlated than stimuli with very different values of *s*. As in [4], we take *Q* = 12. Similar to the standard task we analyzed so far, sensory inputs with *s* = 1,…, *Q*/2 are associated with category `A`, while *s* = *Q*/2,…, *Q* are associated with category `B`. Note that, as in the simple categorization task we analyzed so far, sensory input vectors are linearly separable for every value of Σ.

To introduce heterogeneity in the intermediate layer, we add an offset, so Eq. 5 becomes

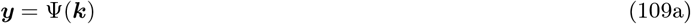

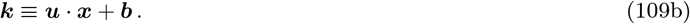

The entries of ***b*** are fixed bias terms that control the value of baseline activity for each neuron. We generate those entries from a zero-mean Gaussian distribution with standard deviation 0.2.

In contrast to the model we analyzed so far, initial activity is characterized by non-trivial activity measures. Specifically, initial population tuning is characterized by non-vanishing category correlation; the latter is modulated both by heterogeneity (which tends to increase signal correlations) and the continuous inputs structure (which tends to decrease them). For our choice of parameters, these two effects roughly balance each other, so that initial activity is characterized by initial correlation that is small in magnitude (Figs. S6A).

We investigated numerically the evolution of activity with learning for this model. Two sample circuits are shown in Figs. S6B-C; extensive analysis is presented in Figs. S6D-E. We find that the behaviour of both category selectivity and correlation is qualitatively consistent with the behaviour of the simpler model analyzed so far. Specifically, we find that average category selectivity increases over learning (Figs. S6D); this behaviour is robust, and does not depend on circuit details. For completeness, we tested two definitions of category selectivity. The first one is identical to Eq. 56; as initial activity is structured, this gives slightly positive initial values; the second one (which is used in related experimental work [3, 4]) is again identical to Eq. 56 – but pairs of stimuli *ss′* are subsampled in a way that is tailored to inputs structure to yield vanishing initial selectivity. We show in Fig. S6D that both selectivity definitions give qualitatively similar results. Whether category correlation increases or decreases over learning depends, on the other hand, on parameters (Figs. S6B-C and E). Correlation depends on parameters in a way that is consistent with the simple task: it is strongly modulated by properties of the readout activation function Φ (Fig. S6E, different shades of gray). It also depends on the activation function of neurons in the intermediate layer Ψ (Fig. S6E left). Finally, it decreases with the learning ratio *η_w_*/*η_u_* (Fig. S6E center) and with the number of stimuli *Q* (Fig. S6E right).

### 5 Context-dependent categorization task

#### 5.1 Context-dependent task: task definition

The second task we consider is a context-dependent categorization task. On each trial, both a stimulus, and a context cue, are presented to the network. For simplicity, we assume that the number of stimuli and context cues is identical, and is equal to *Q*. As in the simple task, each stimulus is represented by an input vector ***μ**^s^*, with *S* = 1,…, *Q*; each context cue is also represented by an input vector, denoted ***ν***^*C*^, with *C* = 1,…, *Q*. The entries of both vectors, ***μ**^s^* and ***ν**^C^*, are generated independently from a zero-mean, unit-variance Gaussian distribution. The total sensory input on each trial, ***x**^s^*, is given by the linear combination of the stimulus and context cue inputs,

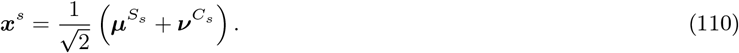

All combinations of stimuli and context cues are permitted; the total number of trials and sensory inputs is thus *P* = *Q*^2^. Each trial s is thus specified by a stimulus and context index: *s* = (*S_s_C_s_*). In contrast to the simple task, sensory input vectors are not orthogonal among each other; using Eq. 110, we see that to the leading order in *N*,

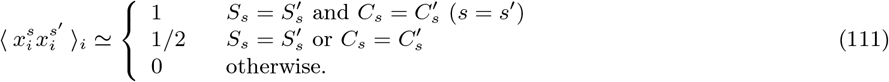

The task is defined as follows. When the context cue *C* ranges between 1 and *Q*/2, context takes value `1`. In context `1`, the first half of the *Q* stimuli is associated with category 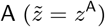, and the second half with 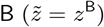. When the context cue *C* ranges between *Q*/2 and *Q*, context takes value `2`. In context `2`, stimuli-category associations are reversed: the first half of the *Q* stimuli is associated with category 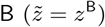, and the second half with 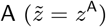.

Correlations in the sensory inputs (Eq. 111) are such that, for every value of *Q*, inputs are not linearly-separable [39]. For *Q* = 2, the task is equivalent to a classical XOR computation. We focus however on *Q* > 2, for which each context is signaled by more than one context cue. As in experimental work [8, 9, 52], this allows to dissociate the activity dependence on the abstract variable context from the sensory variable context cue (see Eqs. 122 and 123 in Section 5.3).

We start by writing down explicit expressions for the activity (Eq. 29) in the current task (Sections 5.2). We then derive the expressions that quantify how activity measures, such as selectivity and correlations, evolve over learning (Sections 5.3, 5.4 and 5.5). These expressions are rather complex, and require numerical evaluation. To gain further mathematical insight, in Sections 5.6, 5.7 and 5.8 we consider specific cases and quantities, and derive their behaviour analytically.

#### 5.2 Context-dependent task: computing activity

We start by computing the value of coordinates *c^s^*, which are solution to the linear system in Eq. 26. As in Section 4.2 (see also Section 4.7), we neglect the variable term 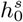 in the left-hand side of that equation and, after a small amount of algebra, we find that it can be rewritten as

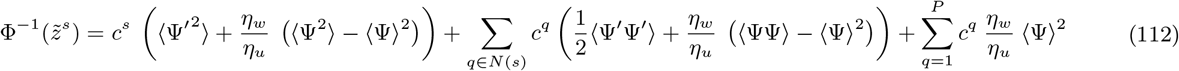

where we used the short-hand notation *N*(*s*) to indicate the set of trials that are *neighbours* to s (i.e., trials that have either the same stimulus or the same context cue of *s*). We have used the notation 〈*FF*〉 to indicate the average over the product of two nonlinear functions, *F*, whose arguments are given by two zero-mean and unit-variance Gaussian variables with covariance 1/2. I.e.,

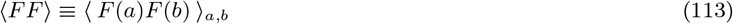

where both *a* and *b* are zero-mean, unit-variance Gaussian random variables with covariance 1/2. Detail on how these averages are computed numerically is given in Section 6.3 (Eq. 186).

As in the simple task (Eq. 46), because the left-hand side can take on only two values, the coordinates *c^s^* can take on only two values,

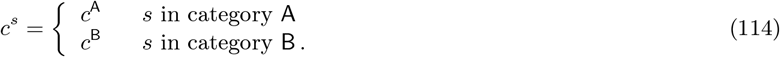

The values of *c*^A^ and *c*^B^ are determined by the same linear system as in Eq. 47, except now *α* and *β* are given by

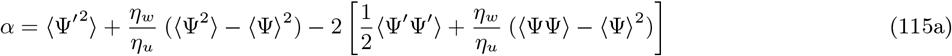

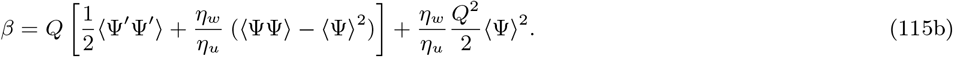

To derive the expression above, we used the fact that every sensory input has 2*Q* – 2 neighbours, of which *Q* – 2 are associated with the same category, and *Q* with the opposite one. The final expression for *c*^A^ and *c*^B^ is thus given by Eq. 49; that expression depends on 7, which is given in Eq. 50.

By comparing Eq. 115 with 48 we see that, with respect to the simple task, the expressions for *α* and *β* include extra terms (shown in square brackets in the right-hand side of Eq. 115). These arise because, unlike in the simple task, different inputs can be correlated (Eq. 111). The extra term in the expression for *β* (Eq. 115b) scales with *Q*, while the extra term for *α* (Eq. 115a) does not; this indicates the typical value of *γ* (Eq. 50), which is proportional to *β/α*, is larger in this task than in the simple one. This in turn implies that the parameter region where one has approximately *c*^A^ ≃ *c*^B^ is larger in the current task than in the simple one; this approximation will later be used in Section 5.8. In the simple task, the parameter region where *c*^A^ ≃ *c*^B^ coincided with the region where category correlation were negative (Eq. 74, Section 4.4). This suggests that the parameter region where correlations are negative, also, is larger in this task than in the simple one. As it will be shown in Section 5.4, however, the expressions for correlations are much more complex in the current task than Eq. 74; this hypothesis thus needs to be carefully verified – which is done, using numerical integration, in Fig. S8C.

Since this task is an extension of the XOR task, sensory inputs are not linearly-separable. This shows up as a singularity when the intermediate layer is linear (e.g., Ψ(***x***) = ***x***). Indeed, in that case, the value of *γ* (Eq. 50) diverges, which in turn means both *c*^A^ and *c*^B^ diverge (Eq. 49). That’s because *γ* is proportional to the ratio *β/α*, and α vanishes, while *β* does not. To see that *α* vanishes, we use Eqs. 115a to write

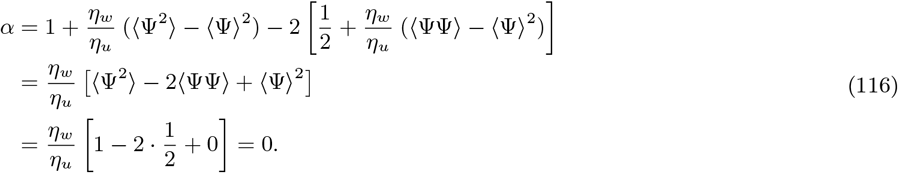

When the activation function Ψ is nonlinear, instead, the values of *c*^A^ and *c*^B^ are finite; their magnitude depends on how close to linear Ψ is in its effective activation range.

To conclude our characterization of activity, we evaluate spanning vectors, ***v**^q,s^*, by combining Eqs. 28 and 111. Unlike in the simple task, for each activity vector, ***y**^s^*, there exists more than one spanning vector; those are given by ***v**^ss^*, and all vectors ***v**^qs^* for which *q* ∈ *N*(*s*). Eq. 29 thus reads

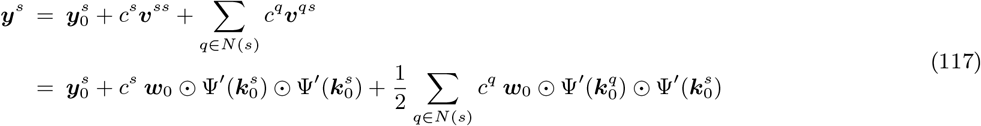

where the second line follows from Eq. 28 and the coordinates *c^q^* take values *c*^A^ or *c*^B^ depending on the category ***x**^q^* is associated with (Eq. 114). Using the notation *s* = (*S_s_C_s_*), Eq. 117 can also be written in the compact form

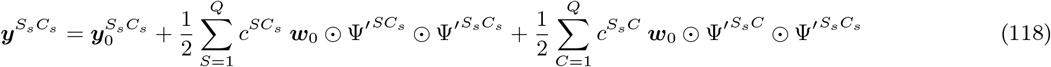

where we used the short-hand notation 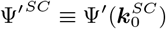.

To isolate the effect of the nonlinearity Ψ, it will be instructive (see Sections 5.6 and 5.7) to also compute the synaptic drive, ***k**^s^*, after learning. Using Eq. 5b and 18a, it is easy to see that

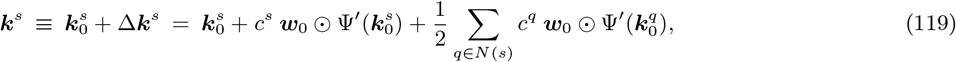

or, equivalently,

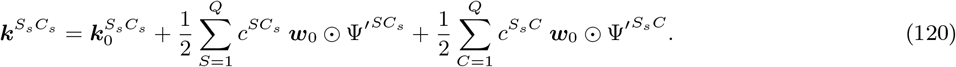

We conclude with a remark on the geometry of the spanning vectors, ***v**^qs^*. As in the simple task, those include a component that is aligned with the initial readout vector, ***w***_0_, and a residual component that is perpendicular to it, *δ**v**^qs^* (Eq. 31). In the simple task, residual components could be neglected (Eq. 55) because they were orthogonal to each other, and did not contribute to novel activity structure. In this task, residual components are not, in general, orthogonal to each other, and thus cannot be neglected. In fact, we have

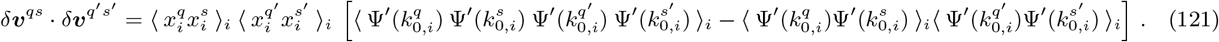

The term in the right-hand side can be non-zero even when *sq* are different from *s′ q′*; this is due to Eqs. 43b and 111, which imply that *k*_0,*i*_ variables can be correlated among each other. The fact that residuals *δ**v**^qs^* cannot be neglected implies that activity evolution is not effectively one-dimensional, as it was the simple task, but higher-dimensional (this is evident in the PC plots in Fig. S7C-D). All the directions along which activity evolve are, however, correlated with the initial readout vector ***w***_0_ (Eq. 30).

#### 5.3 Context-dependent task: category and context selectivity

In the present task, we can compute category, as well as context selectivity. In analogy with category selectivity, Eq. 56, context selectivity is defined as

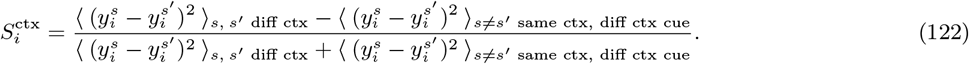

Note that, in the average over pairs of trials from the same context, we excluded pairs of trials with the same context cue. This was done to exclude the possibility that context selectivity increases simply because activity in response to the same context cue become more similar over learning. For completeness, we also compute

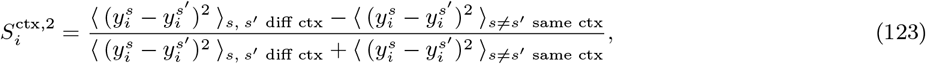

which we plot in Figs. S7A-B. Those plots show that the behaviour under this definition is similar to that of Eq. 122.

We are interested in deriving theoretical expressions for average category and context selectivity, obtained by averaging Eqs. 56 and 122 (or 123) over *i*. For the present task, that is hard. Consequently, we use results from the simple task (Section 4.3) which indicated that, in the limit *N* ≫ *Q* ≫ 1, average category selectivity can be approximated with the category clustering measure, Eq. 67; the latter is equivalent to separately averaging the numerator and denominator of selectivity over neurons.

For category, clustering is the same as in the simple task, Eqs. 67 and 68, which we repeat here for convenience,

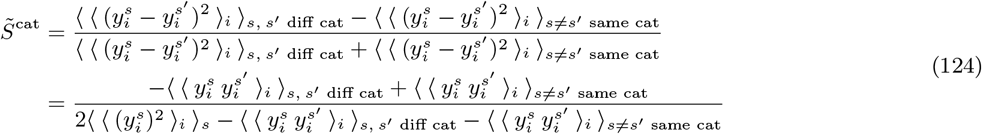

where we used the statistical homogeneity of activity vectors. Similarly, for context selectivity, we may write

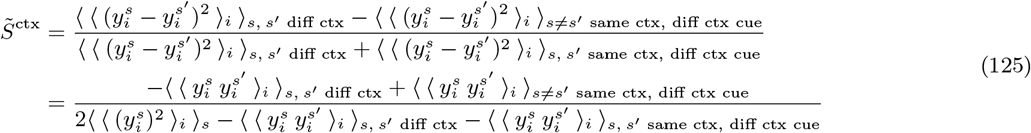

and

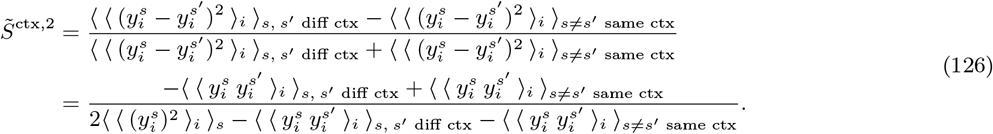

To evaluate those expressions, we need the normalized dot products over activity, 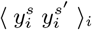. These are computed in Section 5.5. Finally, averages over trials are performed numerically. The resulting theoretical estimates for 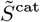 and 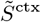 are shown in Figs. S7B and S10.

In Figs. S10A-C, we compare theoretical estimates with simulations. Agreement is relatively good, although it is worse than for the simple task; as argued in Section 3.4, that is expected. Note that the values of average selectivity and clustering are not close (this is only verified in the *N* ≫ *Q* ≫ 1 limit, and would require values of N larger than those used in simulations); the qualitative behaviour of the two quantities is, however, identical. In Fig. S7B, we plot the theoretical estimates across a broad range of task and circuit parameters. These theoretical estimates indicate that, in all cases, category (Eq. 124) and context (Eqs. 125, 126) selectivity increase. This is in agreement with simulations, which are reported in Fig. S7A.

#### 5.4 Context-dependent task: category and context correlation

To quantify how the population as a whole encodes category and context, we evaluate category and context correlations. Those quantities, denoted *C*^cat^ and *C*^ctx^, are given by the average Pearson correlation coefficient for trials in different categories and contexts. *C*^cat^ is defined as in Eq. 71. Similarly, *C*^ctx^ is defined as

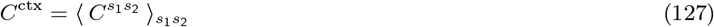

where *s*_1_ and *s*_2_ are indices that denote, respectively, trials from contexts `1` and `2`. Similarly to Eq. 72, the Pearson correlation coefficient *C*^*s*_1_*s*_2_^ given by

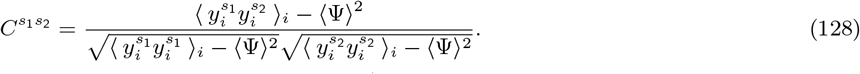

To evaluate these expressions, we use the normalized dot products 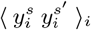 that are computed in Section 5.5. Averaging over trials is, finally, done numerically.

For completeness, we also consider the alternative definition of correlations, where activity is averaged over trials first, and then the Pearson correlation is computed. The alternative definition for category correlation is identical to Eq. 79. The alternative definition for context correlation is given by

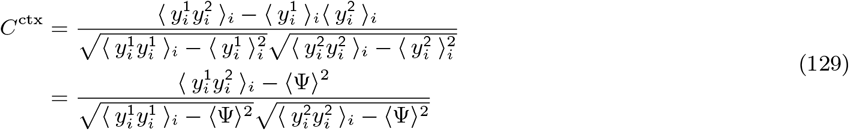

where we have defined

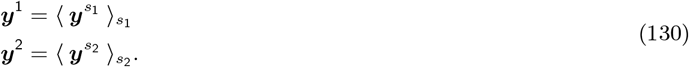

For the current task, there exists no simple mathematical relationship between correlations obtained from the standard, and the alternative definition. We thus checked numerically the behaviour of both quantities; results are reported in Fig. S7B. As in the simple task, we found that the qualitative behaviour of both quantities is not fixed, but depends on task and circuit parameters. This is in agreement with simulations, which are illustrated in Fig. S7A.

#### 5.5 Context-dependent task: computing normalized dot products

To conclude, we illustrate how normalized dot products, Eq. 84, are computed for the current task. We start from Eq. 91, which we repeat here for completeness,

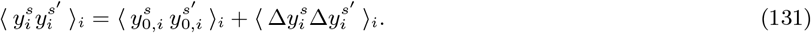

The first term of the right-hand side reads

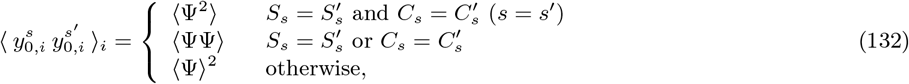

where we used Eq. 5 together with Eqs. 43b and 111. Using Eq. 29 together with 28, the second term of the right-hand side of Eq. 131 reads

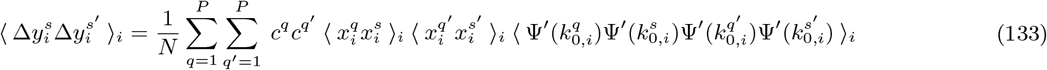

where sensory input correlations, 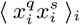, are given in Eq. 111.

Because 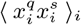 can be non-zero even when *s* ≠ *q* (Eq. 111), the number of non-zero terms in the sum in Eq. 133 is, in general, large. Each term contains an average, 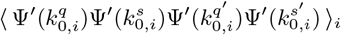, that includes four nonlinear functions. The value of those averages is specified by the correlations among the arguments, *k*_0, *i*_, which in turn depend on the values of *s, q, s′* and *q′* (Eq. 111, via Eq. 43b). Averages are evaluated numerically; detail on how this is done is given in Section 6.3.

This procedure yields a set of normalized dot products that can be used to evaluate, numerically, the expressions for activity selectivity and correlation derived in Sections 5.3 and 5.4. As we rely on numerics, the results we obtain in this way are hard to interpret. For this reason, in the next sections we focus on specific cases were results can be obtained analytically; this allows us to extract a more intuitive understanding of how activity measures evolve over learning.

#### 5.6 Detailed analysis of context selectivity

We start clarifying how context selectivity increases over learning. Results from simulations, and numerical integration of Eq. 125, indicate that context selectivity increases for the synaptic drive, ***k**^s^*; this increase is then reflected in the activity, ***y**^s^* (Figs. S7A-B and S10B). In this section, we analyze the behaviour of context selectivity for the synaptic drive. Focusing on the synaptic drive, instead of activity, allows us to derive results analytically. In the following, we start from Eq. 125 and show that, for the synaptic drive ***k**^s^*, the value of 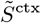 increases over learning. At the end of this section, we comment on the insights provided by such derivation.

We start by simplifying the sums over trials contained in Eq. 125, which involve pairs of trials *ss′* from the same, or different context. To this end we observe that, because of task symmetries, these sums involve a large number of identical terms; for example, the term with *s* = (11) and *s′* = (12) is identical to *s* = (21) and *s′* = (22) (both pairs of trials are neighbours, and are associated with the same category). We thus perform averages over a reduced, and less redundant subset of pairs of trials. First, we consider only two values of *s*: for concreteness, we take *s* = (11) and 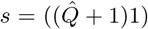, where we defined

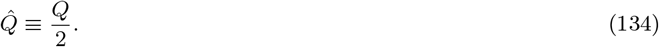

These *s* trials are associated, respectively, with category `A` and `B`. Second, for each value of *s*, we consider *s′* trials with context cue equal to *C* = 2 and 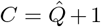; these are associated, respectively, with context `1` and `2` (note that *C* = 1 must be avoided, as trials with the same context cue must be excluded, see Eq. 125). This allows us to rewrite the averages contained in Eq. 125 as

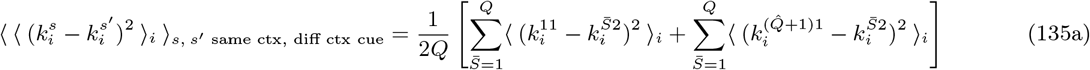

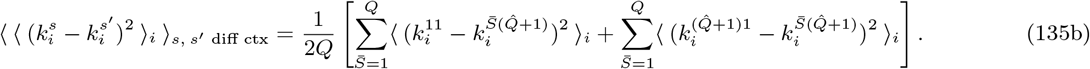

The sums over 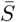 can further be simplified. By using again symmetries, we have:

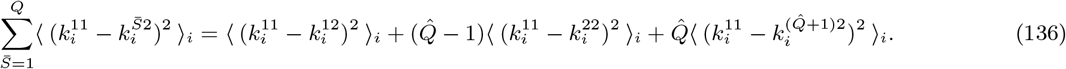

We can do the same for the other sums, yielding:

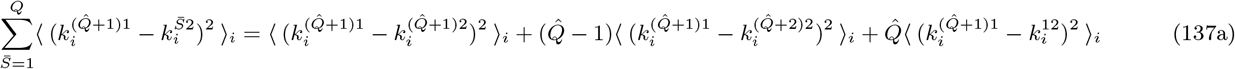

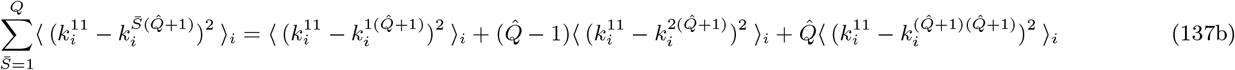

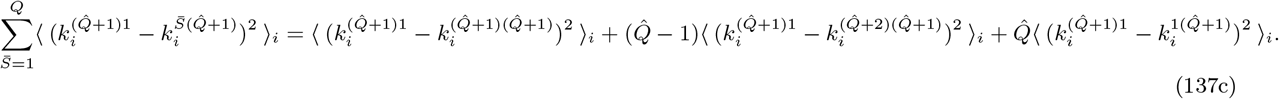

It is easy to verify that, before learning starts, the right-hand sides of Eqs. 135a and b are identical. This implies that the initial value of context selectivity, Eq. 125, vanishes (Fig. S10B). To show that context selectivity increases over learning, we thus need to show that the numerator of Eq. 125 becomes positive over learning. This is equivalent to show that Eq. 135a is smaller than Eq. 135b. Using Eq. 136 and 137, this condition can be rewritten as

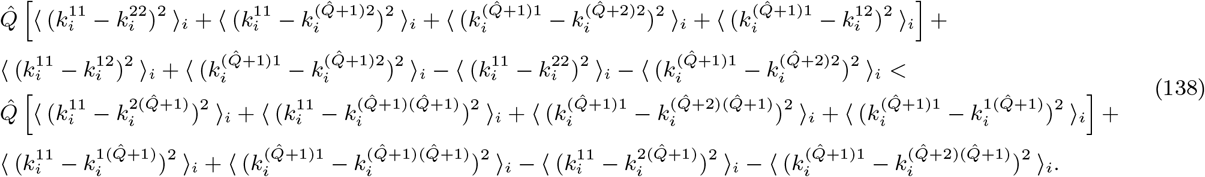

We now use Eq. 119 to write

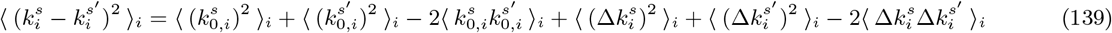

where the terms containing the cross-products between 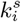 and 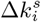 vanish on average because of Eq. 8a. By using the statistical homogeneity of activity across contexts, we can rewrite Eq. 138 as

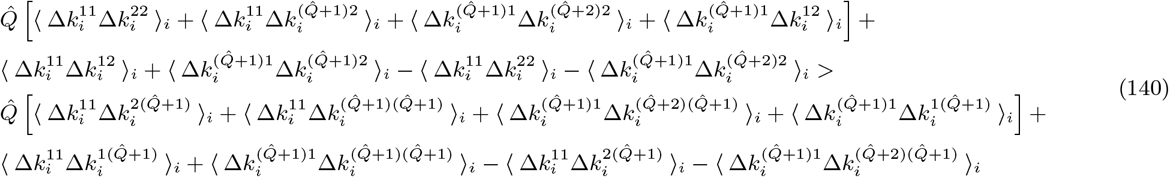

or, re-arranging terms,

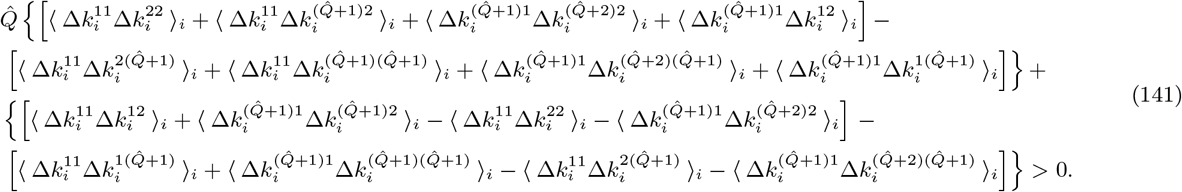

To show that context selectivity increases over learning, we need to verify that the equation above holds. To this end, we evaluate analytically the normalized dot products 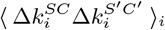 for each pair of trials involved. This is done in the next paragraph; here we simply use those results (Eqs. 156, 157, 158 and 159).

We start evaluating the difference within the first set of curly parenthesis of Eq. 141, which correspond to the dominant contribution in *Q*. By using Eq. 159, we see that this can be rewritten as

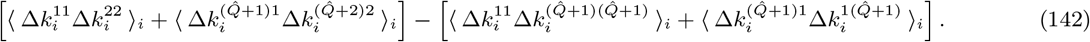

Using Eq. 158, this becomes

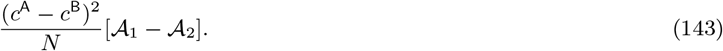

We then evaluate the difference within the second set of curly parenthesis. Using Eqs. 156, 157, 158 and 159 it is straightforward to see that that difference vanishes. Putting results together, our condition to verify (Eq. 141) becomes simply:

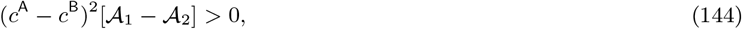

which is satisfied whenever 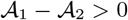. This is always verified, as from Eqs. 150 and 151 we have

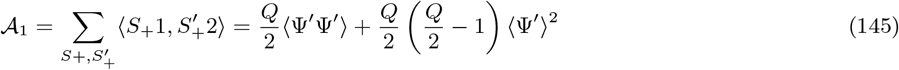

while

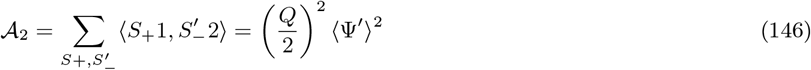

so that

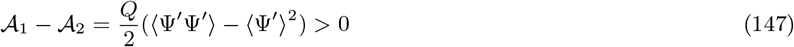

which concludes our derivation. We remark that Eq. 147 vanishes when Ψ is linear. This indicates that, even if context selectivity also increases for synaptic drives (which are a linear transformation of the sensory inputs), this phenomenon is due to the nonlinearity of activation functions.

##### Computing normalized dot products

We now compute the normalized dot product expressions, 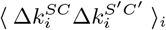, for each pair of trials involved in Eq. 141. We illustrate in detail how one example dot product, 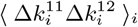, is computed. Other expressions are computed in a similar way; results are given below (Eqs. 157, 158 and 159).

We start from:

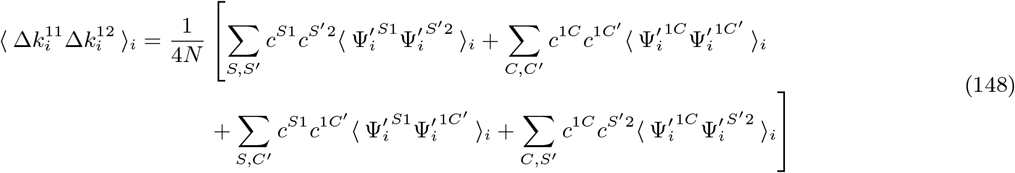

which was derived from Eq. 120 together with Eq. 8a. We then rewrite the sums in the right-hand side by expanding each index in two set of indices: one running from 1 to *Q*/2 (denoted by the subscript +), and one running from *Q*/2 + 1 to *Q* (denoted by the subscript –). The first sum in Eq. 148 becomes:

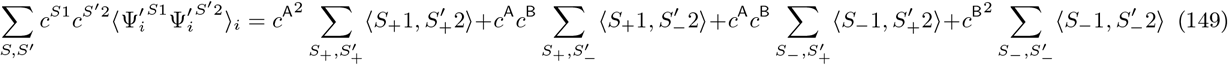

where we have used the short-hand notation 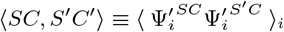. We now observe that

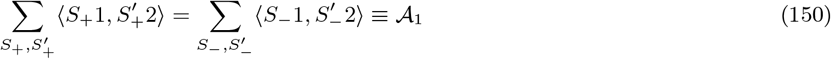

while

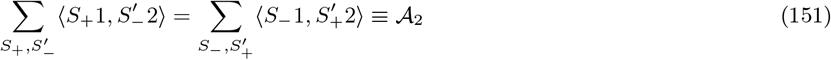

so that

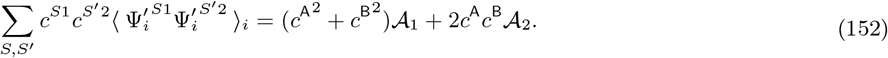

The second sum in Eq. 148 gives:

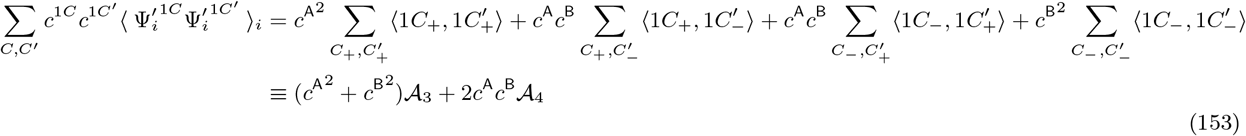

by appropriately defining 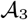 and 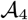. The third sum gives:

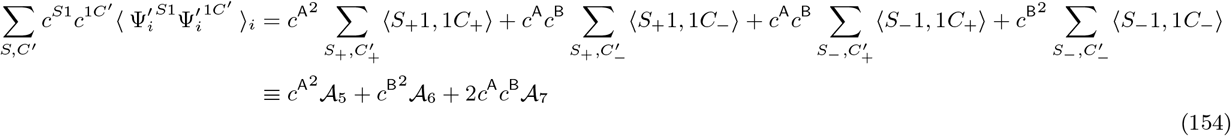

and, similarly, the fourth one:

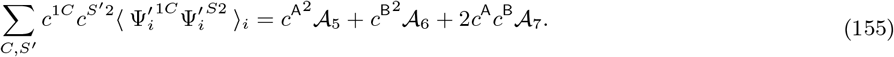

By putting those results together, we conclude that

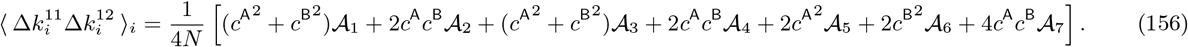

We can use the same procedure to evaluate dot products for all the remaining pairs of trials. This gives:

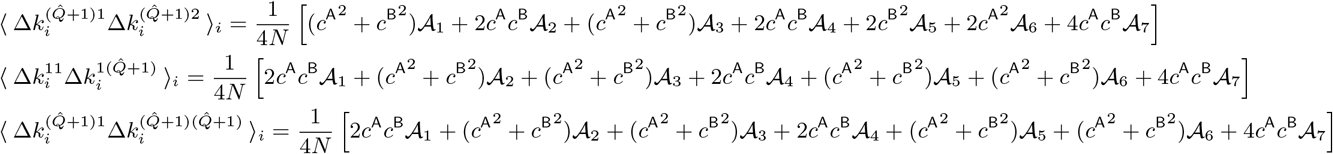

while

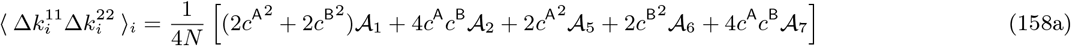

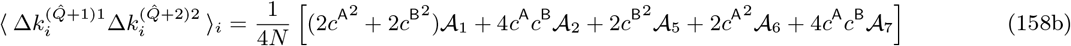

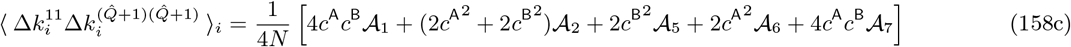

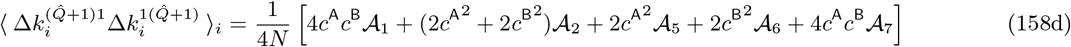

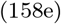

and

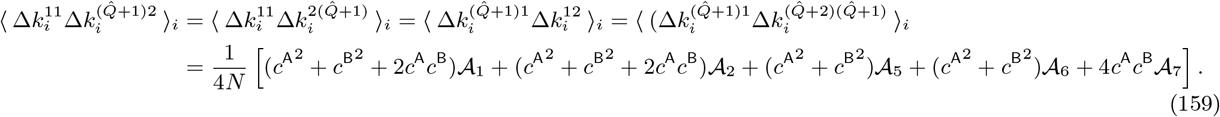

All the 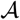 coefficients can easily be evaluated analytically. However, we have shown in the previous paragraph that the only coefficients that do not cancel in Eq. 141 are 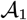 and 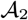; these two are evaluated analytically in Eqs. 145 and 146.

##### Extracting intuition

Can we derive a more intuitive picture of why and how context selectivity increases over learning? We have seen in the previous paragraphs that context selectivity increases because the difference within the first set of curly parenthesis of Eq. 141 is positive (while the difference within the second set of curly parenthesis vanishes). To simplify the math, we assume that *c*^A^ = –*c*^B^; this condition thus reads:

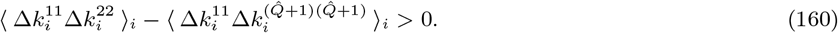

(With respect to Eq. 142, we could get rid of pairs of trials with 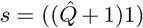 because, when *c*^A^ = –*c*^B^, they give identical results to *s* = (11).)

Eq. 160 indicates that, over learning, activity from trial *s* = (11) becomes closer (i.e., more correlated) to activity from trials with the same category and context, such as *s′* = (22), than trials with the same category but different context, such as 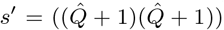. On the contrary, activity from trial *s* = (11) becomes equally close to activity from trials with different category and same context, such as 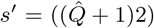, and trials with different category and different context, such as 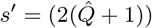. This can be seen from Eq. 159, from which

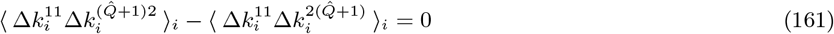

follows.

The geometrical relationships implied by both Eqs. 160 and 161 can be easily verified in Fig. S7C, which shows the synaptic drive from simulated circuits; the middle panel shows a circuit for which we have exactly *c*^A^ = –*c*^B^. Taken together, Eqs. 160 and 161 indicate that the increase in context selectivity comes from activity clustering by context over learning; such clustering is, however, category-dependent. This leads to the emergence of four statistically-distinguishable clouds, one for each combination of category and context. This is visible in simulated activity from Fig. S7C, and is illustrated in Figs. 7A-C.

#### 5.7 Detailed analysis of category selectivity

We now provide extra detail on the behaviour of category selectivity. We start explaining why, as observed in Figs. 6A and S10A, initial selectivity does not vanish, but is weakly negative. This phenomenon is observed both for the synaptic drive ***k**^s^* and the activity ***y**^s^*; for the sake of simplicity, we focus on the former.

Consider for a moment the case *Q* = 2 (XOR computation). The geometry of the initial synaptic drive is in that case particularly simple, and is illustrated in Fig. S8D. As can be easily verified by using Eqs. 5b and 110, each synaptic drive is given by the linear superposition of two vectors: a vector among 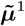 and 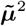, and a vector among 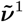 and 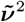. Vectors 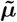 and 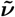 are obtained by applying the initial connectivity ***u***_0_ to vectors ***μ*** and ***ν*** (Eq. 110); for example, 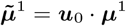. In the plane spanned by vectors 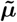 and 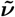, the geometry of synaptic drives is square-like (Fig. S8D). To verify that, observe that the squared distance between consecutive vertices is identical – for example,

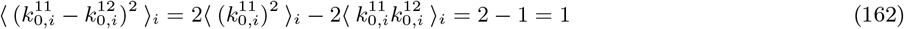

where we used Eq. 43 together with 111. Opposite vertices have instead double squared distance – for example

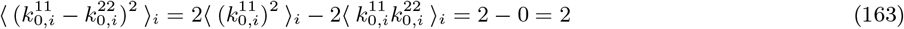

as expected for a square. Importantly, consecutive vertices are associated with different categories, while opposite vertices are associated with the same category; this implies that initial category selectivity is negative. In fact, using Eqs. 162 and 163 into Eq. 124 yields:

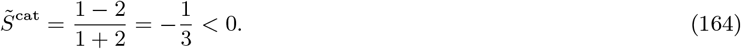

It is easy to see that initial category selectivity is negative also when *Q* > 2. However, its magnitude converges to zero as the number of stimuli and context cues, *Q*, increases (Fig. S10A). This is due to the fact that, as *Q* becomes large, both the within-category and the across-category averages in Eq. 124 become dominated by pairs of trials with different stimulus and context cue; activity from those pairs of trials are characterized by identical initial distances (=2, as in Eq. 163), and thus the two averages become similar.

We now shed light on a second phenomenon: the fact that category selectivity increases over learning for the activity ***y**^s^*, but remains identical for the synaptic drive ***k**^s^*. This is observed both in simulations (Figs. S7B and S10A), and in numerical integration of theoretical expressions (Figs. S7A and S10A). To see why this happens, we assume that the number of stimuli and context cues, *Q*, is fairly large (1 ≪ *Q* ≪ *N*). As discussed above, in this limit, initial category selectivity is approximately close to zero. To compute selectivity after learning, we use Eq. 124, and evaluate the within-category and the across-category averages. We compute averages to the dominant terms in *Q*, which correspond to pairs of trials with different stimulus and context cue. Using the same *s* and *s′* trials as in Section 5.6, we obtain

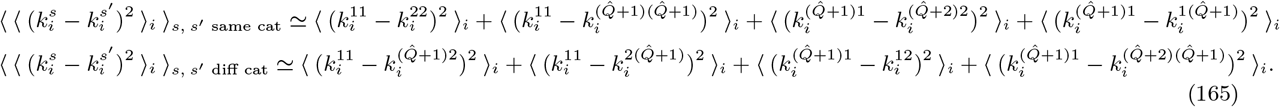

To show that category selectivity does not change over learning, we need to show that the two lines above are identical. Using Eq. 139, this condition can be written as:

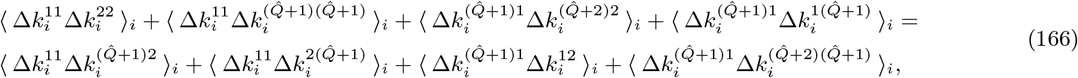

which can now be easily verified by using Eqs. 158 and 159.

Equation 166 indicates that, on average across contexts, synaptic drives from trials with the same category are as close as trials with different category. This geometrical relationship can be easily verified in Fig. S7C, which shows the synaptic drive from simulated circuits. We focus on the middle panel, where we have *c*^A^ = –*c*^B^. The four activity clouds corresponding to different combinations of category and context values are approximately arranged on the vertices of a square; consecutive vertices are associated with different categories, while opposite vertices are associated with the same category. To see why Eq. 166 holds, note that squared distances among synaptic drives associated with different category are approximately identical, while squared distances among synaptic drives associated within the same category are either 0 (approximately, half of the times), or twice the across-category distance (the other half). It is interesting to observe that this square-like configuration, which emerges over learning from an almost unstructured one (Fig. S7C left), strongly resembles the initial configuration of the XOR task (Fig. S8D).

A fundamental feature of this configuration is that synaptic drives are not linearly-separable by category. The activity vectors ***y**^s^*, on the other hand, are linearly-separable. Before learning, linear separability is guaranteed by the nonlinearity Ψ, which makes activity vectors linearly-separable along random directions [39]. After learning, activity vectors become linearly-separable also along task-relevant directions. In the simplified scenario where *η_w_* ≪ *η_u_*, the activity vectors become linearly-separable along ***w***_0_; in the general case, they become linearly-separable along a direction that is correlated with **w**_0_. This is shown in Fig. S7D: the configuration of activity is very similar to synaptic drives, but activity vectors associated with different categories clusters, and thus become linearly separable, along an emerging, orthogonal direction. This drives the increase in category selectivity that was observed both in equations and simulations (Figs. S7A-B and S10A). A further insight on the relationship between selectivity and activity geometry is given in the next section.

We conclude with a remark. Although for activity variables category selectivity robustly increases, the fact that selectivity is weakly negative before learning implies that asymptotic values can be small, or even negative. This is compatible with findings in [74], where very small values of category clustering (Eq. 124) were observed. This observation stresses the importance of measuring, in experimental preparations, neural activity across multiple stages of learning.

#### 5.8 Analysis of patterns of context and category selectivity

In this section, we investigate how changes in context and category selectivity are distributed across neurons.

In the simple task, we found that the magnitude of selectivity changes for a given neuron, *i*, was correlated with the magnitude of the *i*-th entry of the initial readout vector ***w***_0_ (Eq. 65, Figs. 5B-C). This vector defines the direction along which clustering by category takes place. In fact, if one draws the vector joining the centers of the activity clouds associated with different categories, ***y***^A^ and ***y***^B^ (Eqs. 78), the resulting direction is correlated with ***w***_0_ (Eq. 96). This direction is indicated with ***d*** in the main text; cloud centers ***y***^A^ and ***y***^B^ are plotted, in Figs. 3B-C and S1B, as magenta triangles.

In analogy with the simple task, we now hypothesize that the magnitude of changes in context and category selectivity for a given neuron, *i*, is related to the magnitude of the i-th entry of the context and category directions, ***d***^ctx^ and ***d***^cat^. Those coincide with the directions along which clustering to context and category emerges (Figs. 7B-C), and are given by the vectors joining the centers of the activity clouds associated with different contexts (Eq. 130) and categories (Eq. 78). The cloud centers for category and context are plotted, in Figs. 7B-C and S7C-D, as magenta and pink triangles. This assumption is verified in Figs. S9A-B, which shows that selectivity changes and context and category directions are highly correlated. Our reasoning implies that, in order to understand how selectivity changes are distributed across neurons, we need to evaluate the entries of the context and category directions; this is done, analytically, in the rest of this section.

As we are interested in selectivity changes, we focus on activity changes, and approximate

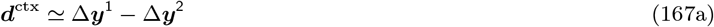

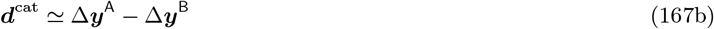

where, similarly to Eqs. 78 and 130, we have taken

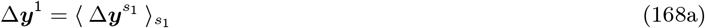

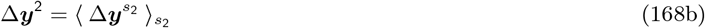

and

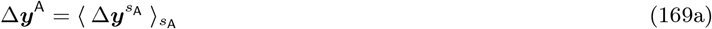

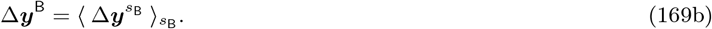

We start with context. We have seen in Section 5.6 that context selectivity can also be studied at the level of the synaptic drive ***k**^s^*, which greatly simplifies the analysis. Starting from Eq. 120, we thus compute

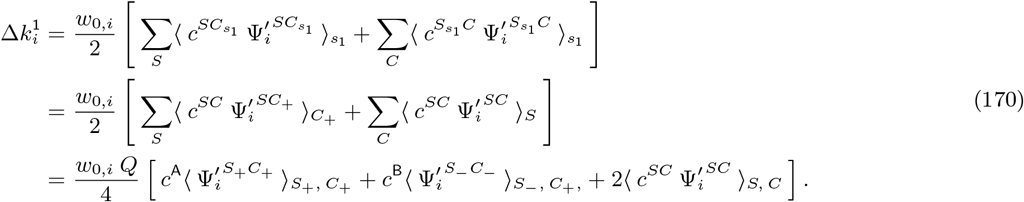

As in Section 5.6, indices *S*_+_ and *S*_−_ (and, similarly, *C*_+_ and *C*_−_) run, respectively, from 1 to *Q*/2 and from *Q*/2 + 1 to *Q*. Similarly,

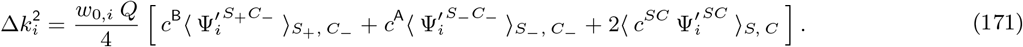

Note that, because of the first two terms in the right-hand sides, Eqs. 170 and 171 are not identical.

To further simplify the analysis, we assume that *c*^B^ ≃ *c*^A^. As discussed in Section 5.2, in the current task, this represents a good approximation for a large space of parameters; we verified with simulations that our main results also hold, qualitatively, in circuits where this approximation fails (notably, in the circuit illustrated in the third column of Fig. 6, see Figs. S9C-D). Combining Eq. 167a with Eqs. 170 and 171, we then obtain

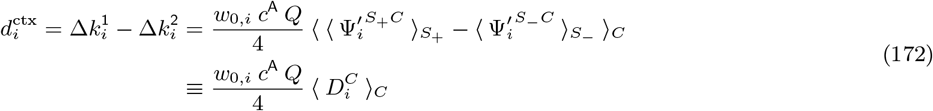

where we have defined

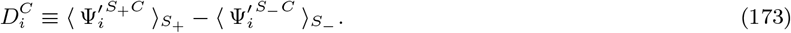

Equation 172 indicates that neurons exhibiting a strong increase in context selectivity are characterized by: (i) strong readout connectivity, before learning, as quantified by *w*_0,*i*_, and (ii) a large value of 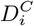, averaged over context cues. 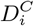 is a function of the response gain function, Ψ′, evaluated before learning; specifically, 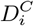 measures the difference in the initial gain in response to the two classes of stimuli (the first half, *S*_+_ = 1,…, *Q*/2, and the second half, *S*_−_ = *Q*/2,…, *Q*). These predictions, which were derived for the synaptic drive ***k**^s^*, also hold, qualitatively, for the activity ***y**^s^* (Fig. 8).

We next compute the category direction ***d**^cat^*; we focus again on the synaptic drive ***k**^s^* rather than activity ***y**^s^*. We observe that, before learning, the centers of synaptic drive vectors associated with category A and B are perfectly identical. In fact,

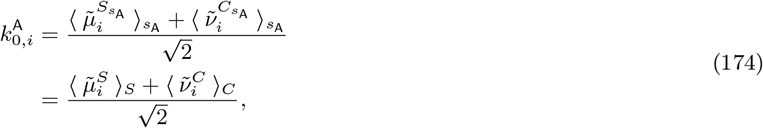

and an identical expression is obtained for 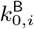. The fact that the centers are identical is due to the fact that sensory inputs for the two categories are collinear, and perfectly intermingled (Fig. S8D). We now consider the synaptic drive changes over learning. Starting from Eq. 120, we have

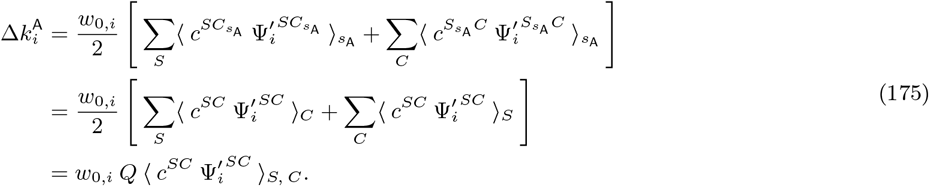

It is easy to show that 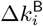 yields the same result, implying that the centers for synaptic drive vectors associated with categories A and B remain identical over learning (Fig. S7C, magenta triangles). This happens because the synaptic drive vectors associated with category A and B remain intermingled, and non linearly-separable, over learning. We conclude that the category axis ***d***^cat^ (Eq. 167b) vanishes, which is in agreement with the observation that category selectivity does not change for synaptic drives (Section 5.7).

To compute ***d***^cat^, we thus turn to activity ***y***. We start from Eq. 118, and write

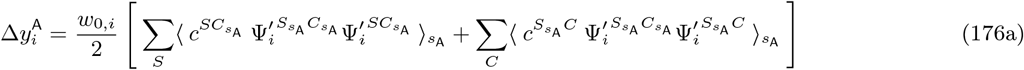

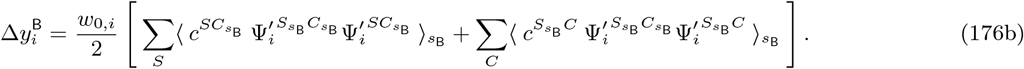

We then expand indices over stimuli and context cues, which yields

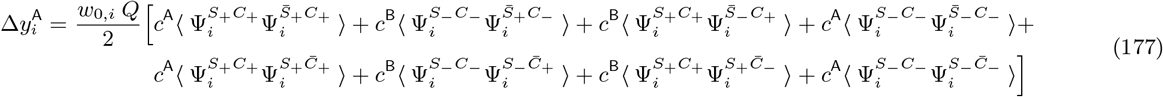

and

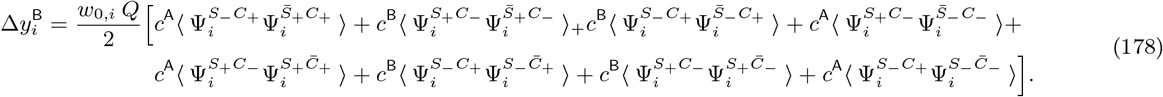

To reduce the clutter, we have removed subscripts after brackets 〈.〉; those indicate an average taken over all the *S* and *C* indices contained within.

As will become clear shortly, the two now centers differ (Fig. S7D, magenta triangles). To simplify those expressions, we again assume that *c*^B^ ≃ *c*^A^; this allows us to write

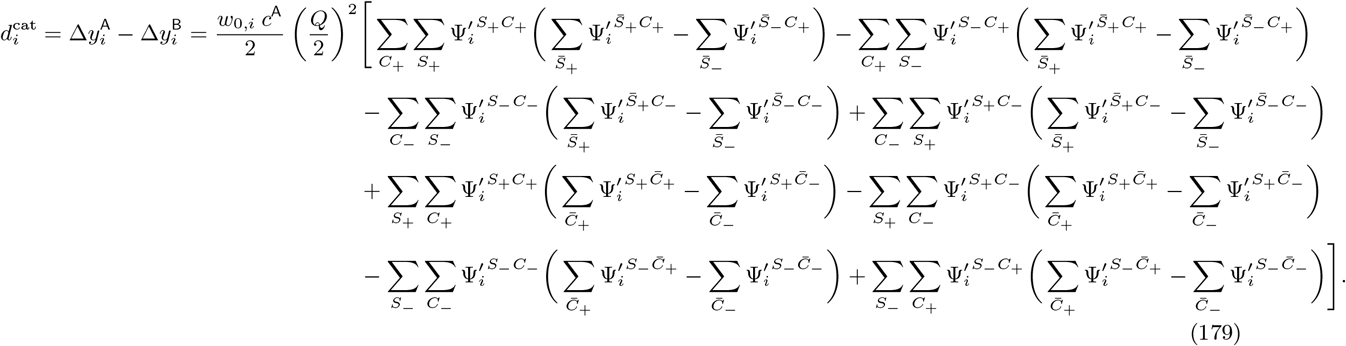

With a little algebra, we can see that

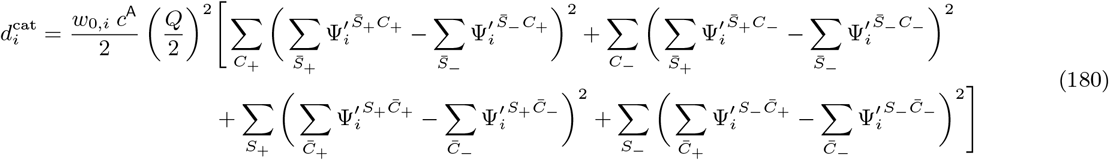

or, equivalently

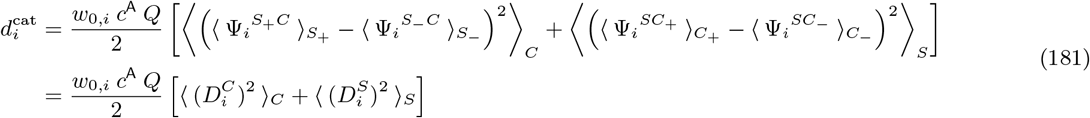

where we have defined

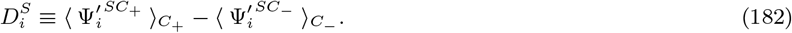

Equation 181 indicates that neurons characterized by a strong increase in category selectivity are characterized by: (i) strong readout connectivity, before learning, as quantified by *w*_0,*i*_, and (ii) large values of 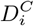 and/or 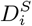, averaged, respectively, over context cues and stimuli.

Note that neurons that are characterized by a strong increase in context selectivity (Eq. 172), which have large *w*_0,*i*_ and 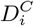 values, are also characterized by a strong increase in category selectivity (Eq. 181). On the other hand, neurons with large *w*_0,*i*_ and 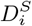 values are characterized by a strong increase in category selectivity (Eq. 181), but not context (Eq. 172). Overall, strongly selective neurons can thus be classified in two groups: one displaying mixed selectivity to category and context, and one displaying pure selectivity to category. By defining the quantity:

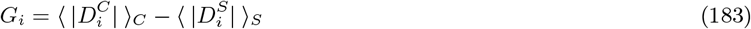

we see that the former group is characterized by larger values of Giwith respect to the latter. This is verified and illustrated in Figs. 8B-C.

### 6 Software

#### 6.1 Circuit simulations

Simulations were implemented with the Python programming language. Gradient-descent learning was implemented with the PyTorch package. We used the *SGD* optimization function, with loss *MSELoss*. On every learning epoch, the batch included all sensory input vectors. Training stopped when the loss function dropped below 10^−5^. Learning rates were taken to be *η* = 0.1 for input connectivity ***u***, and *η* · *η_w_*/*η_u_* (with values of *η_u_*, and *η_w_* as indicated in Section 6.2) for readout connectivity ***w***.

Code will be made available in a public repository upon publication.

#### 6.2 Table of parameters

We summarize below the parameters chosen for the simulations reported in Figures and Supplementary Figures. For figures not included in the tables below, parameters have been detailed in figures captions.

We have taken everywhere *z*^A^ = 0.75, *z*^B^ = 0.25 (note that activity variables range between 0 and 1).

#### 6.3 Evaluation of averages

Evaluating the approximate theoretical expressions for activity measures given in Sections 4 and 5 requires computing a number of Gaussian integrals over nonlinear functions. We compute those averages numerically; details are provided below.

The simplest average,which only involves one nonlinear function, was denoted by 〈*F*〉 (Eq. 45). We rewrite Eq. 45 in an integral form, yielding

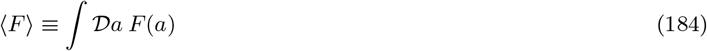

where we have used the short-hand notation

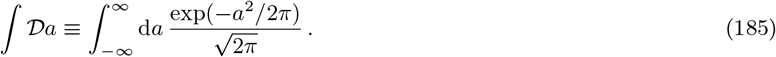

This integral was computed numerically via Hermite-Gaussian quadrature.

Averages involving two nonlinear functions were denoted by 〈*FF*〉 (Eq. 113). We rewrite Eq. 113 in an integral form, yielding

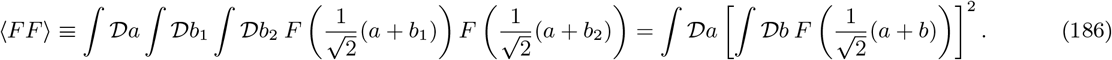

This integral was computed again via Hermite-Gaussian quadrature.

Averages involving four nonlinear functions, such as 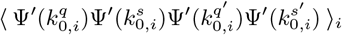 from Eq. 133 (Section 5.5) were computed instead via the function *nquad* from the Python *scipy.integrate* package. We start by rewriting the argument of the average as:

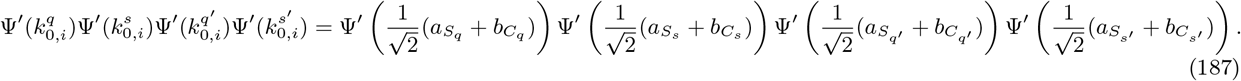

For each value of the stimulus index *S* and the context cue index *C, a_S_* and *b_C_* are two independent, zero-mean and unit-variance Gaussian variables. If the values of *S* and *C* are different across the four trials *q, s, q′* and *s′*, then all *a* and *b* variables involved in Eq. 187 are different, and the average reads

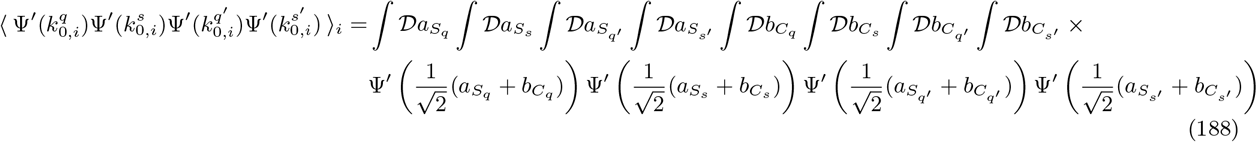

which simplifies into

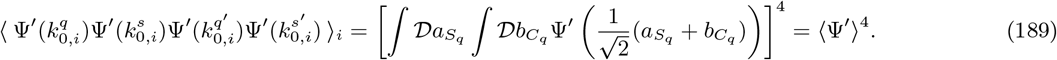

If the stimulus *S* or the context cue *C* are, instead, identical across two o more trials (*q, s, q′* and *s′*), then some of the *a* and *b* variables in Eq. 187 are shared across nonlinear functions. This generates correlations, which determine the final value of the average. For example, assume *S_q_* = *S_s_*, while all other S and C values are different among each other. Then the average reads

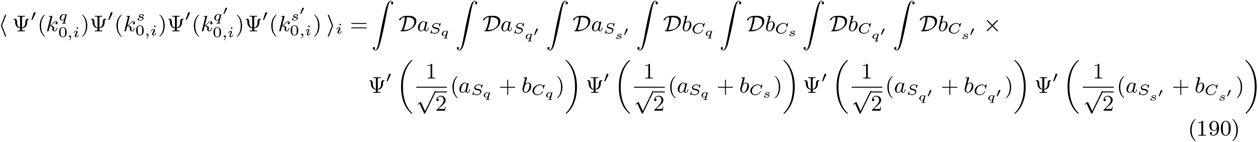

which simplifies into

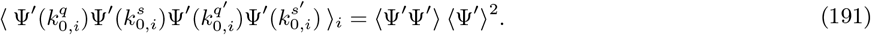

We considered all the possible configurations of *S* and *C* indices that can occur in the context-dependent task, and all the resulting correlation patterns. Then, we used analytics to simplify integrals when possible (as in the cases described above). We finally used numerics to evaluate the remaining integral expressions.

## Supplementary Figures

**Fig. S1:**
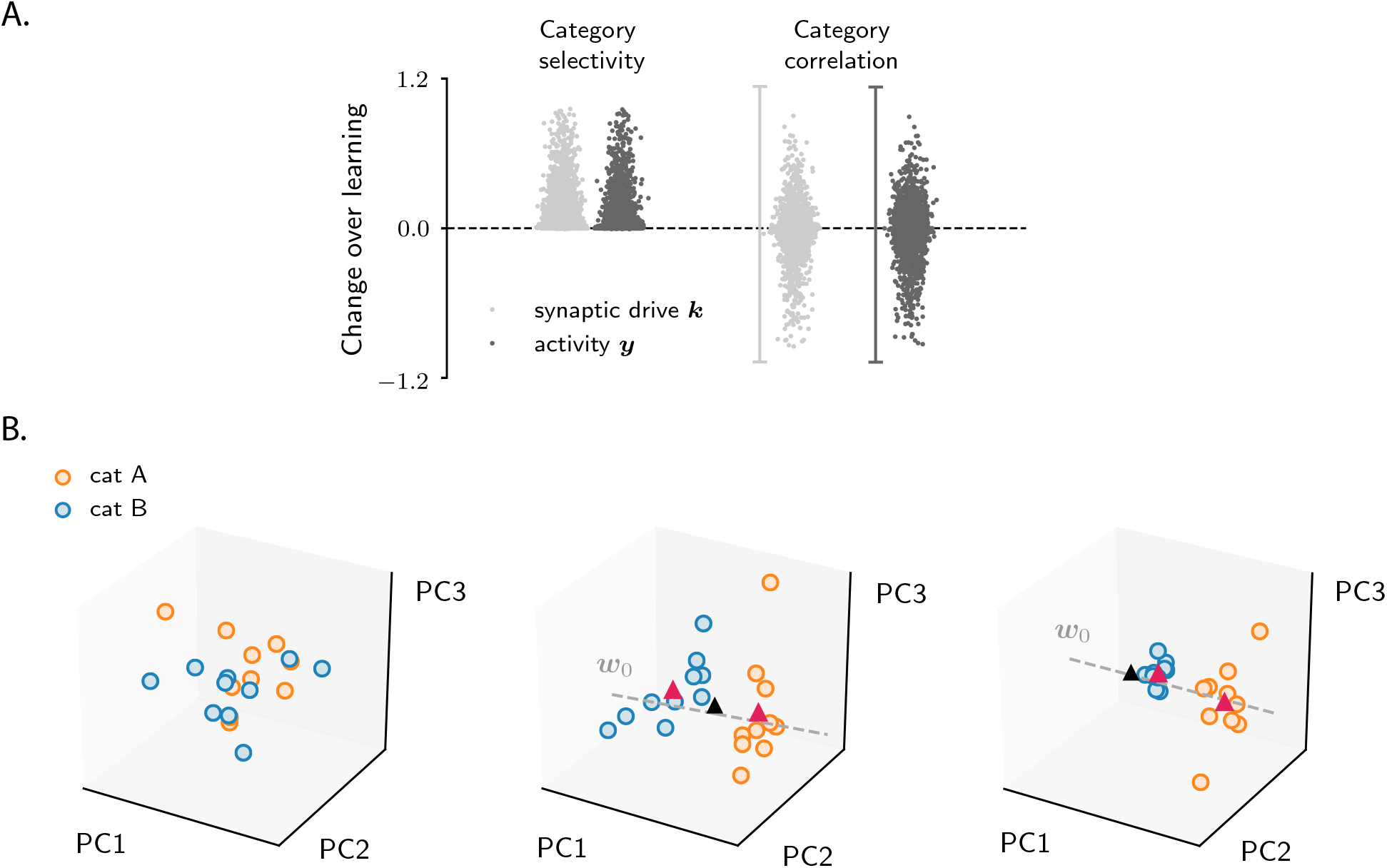
Characterization of activity evolution during the simple categorization task; additional results, part I. **A.** Extensive results from simulations. Each point represents a model. We simulated 5000 different models, with parameters drawn randomly and uniformly from the following ranges: the number of stimuli, *Q*, between 10 and 30; the learning rates ratio, *η_w_*/*η_u_*, between 0 and 1; the gain of Ψ and Φ between 0.05 and 1; the threshold of Ψ and Φ between −2 and 2. Light points refer to the synaptic drive, ***k***, and dark ones to the activity, ***y***. To quantify category selectivity, we used Eq. 56, and then averaged over neurons. To quantify category correlation, we used Eq. 71. The vertical bars in the category correlation plots indicate the range of results obtained when quantifying correlation with the alternative definition in Eq. 79. Note that selectivity changes are positive, while correlation changes take both positive and negative values. For a few parameter sets, learning convergence was pathological (i.e., the loss displayed strong oscillations over epochs, or did not converge); those cases have been excluded from the analysis. **B.** Activity from circuits in Figs. 2 A-D (left), E-H (center) and I-L (right) projected on the first three principal components. Before computing the principal components, we subtracted from activity the mean across trials and neurons. Over learning, activity develops two clouds, one for each category, as schematically illustrated in Figs. 3A-C. The clustering direction is approximately given by ***w***_0_ (dashed grey line). The center of the two clouds is indicated by magenta triangles; the center of initial activity is indicated by the black triangle.

**Fig. S2:**
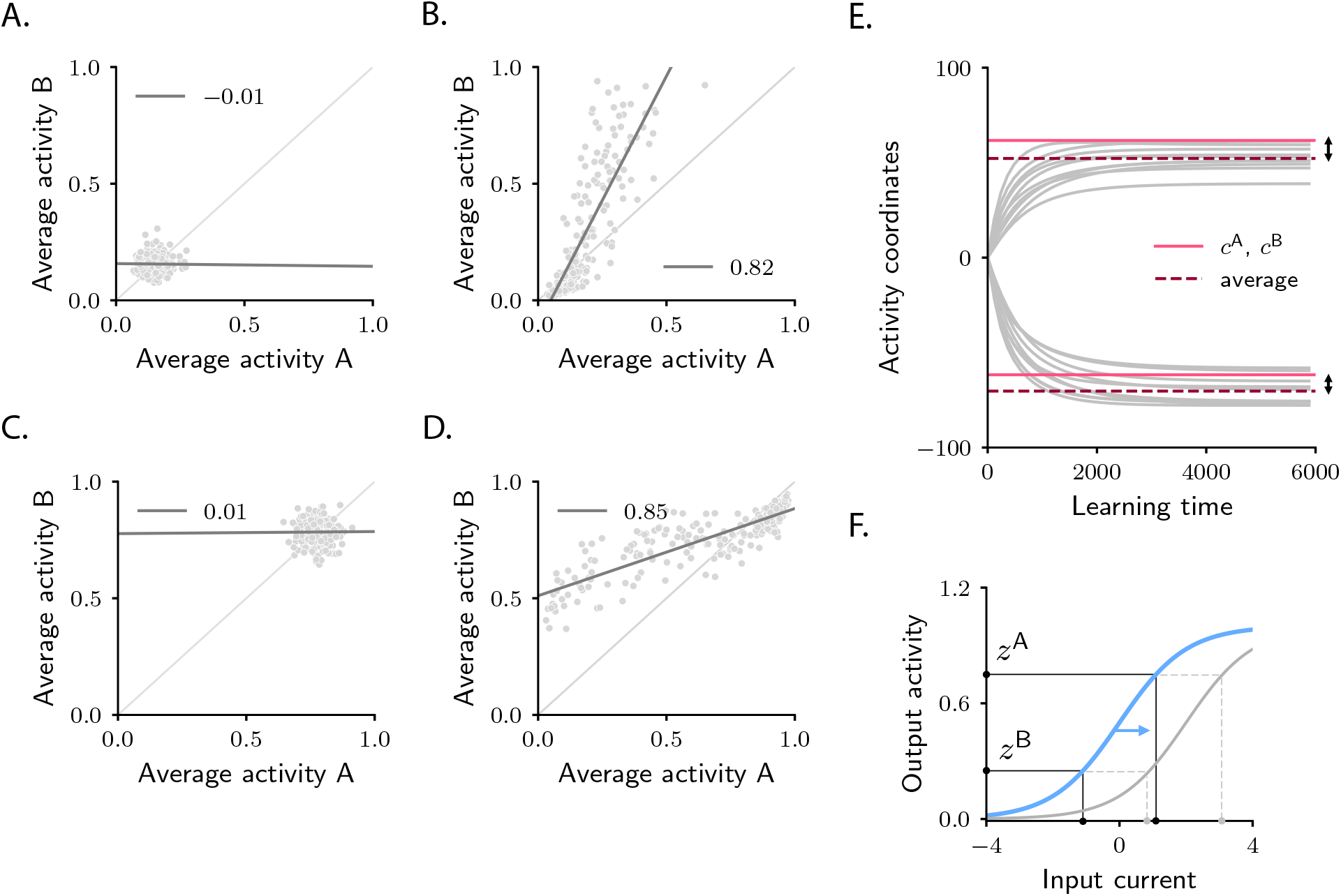
Characterization of activity evolution during the simple categorization task; additional results, part II. **A-D.** Population response to categories A and B, averaged over stimuli. Each dot represents a neuron. Panels A-B and C-D display two sample circuits, different from those displayed in Fig. 2. Panels A and C and B and D display, respectively, pre- and post-learning activity. All details and parameters are as in Fig. 2I-L, except: in A and B, we set the threshold of Φto −2; in C and D, we set the threshold of Ψ to −1.5. In panel B, in contrast to Fig. 2L, the variance of activity in response to B is larger than the variance in response to A. In panel D, as in Fig. 2L, the variance of activity in response to A is larger than the variance in response to B. In contrast to Fig. 2L, however, the mean activity in response to B is larger than the mean activity in response to A (Methods 4.6). Note that, in all cases, category correlation remains positive. **E.** Variability across circuits realizations (Methods 4.7): illustration from a sample circuit. Grey lines illustrate activity coordinates, *c^s^*, as a function of learning time, for all sensory inputs. Those were estimated by taking the dot product Δ***y**^s^*·***w***_0_, and then dividing by 〈Ψ′^2^〉 (Eqs. 55 and 53). Note that *c^s^* values corresponding to sensory inputs in the same category do not saturate to identical values; their average is indicated by dashed lines. Pink continuous lines indicate the values of *c*^A^ and *c*^B^ computed through the theory by neglecting variability (Eq. 49). As predicted by Eq. 99, the dashed and continuous pink lines do not coincide (Methods 4.7). **F.** Activity coordinates *c*^A^ and *c*^B^ depend on the values of Φ^−1^(*z*^A^) and Φ^−1^(*z*^B^) (i.e., the input current needed by the readout neuron to to produce the target activity). For fixed targets *z*^A^ and *z*^B^, those depend on the activation function Φ. Increasing the threshold of Φ can cause Φ^−1^(*z*^A^) and Φ^−1^(*z*^B^) to shift from having opposite sign to having the same sign. This in turn can cause a shift from negative to positive category correlation, see Methods 4.4.

**Fig. S3:**
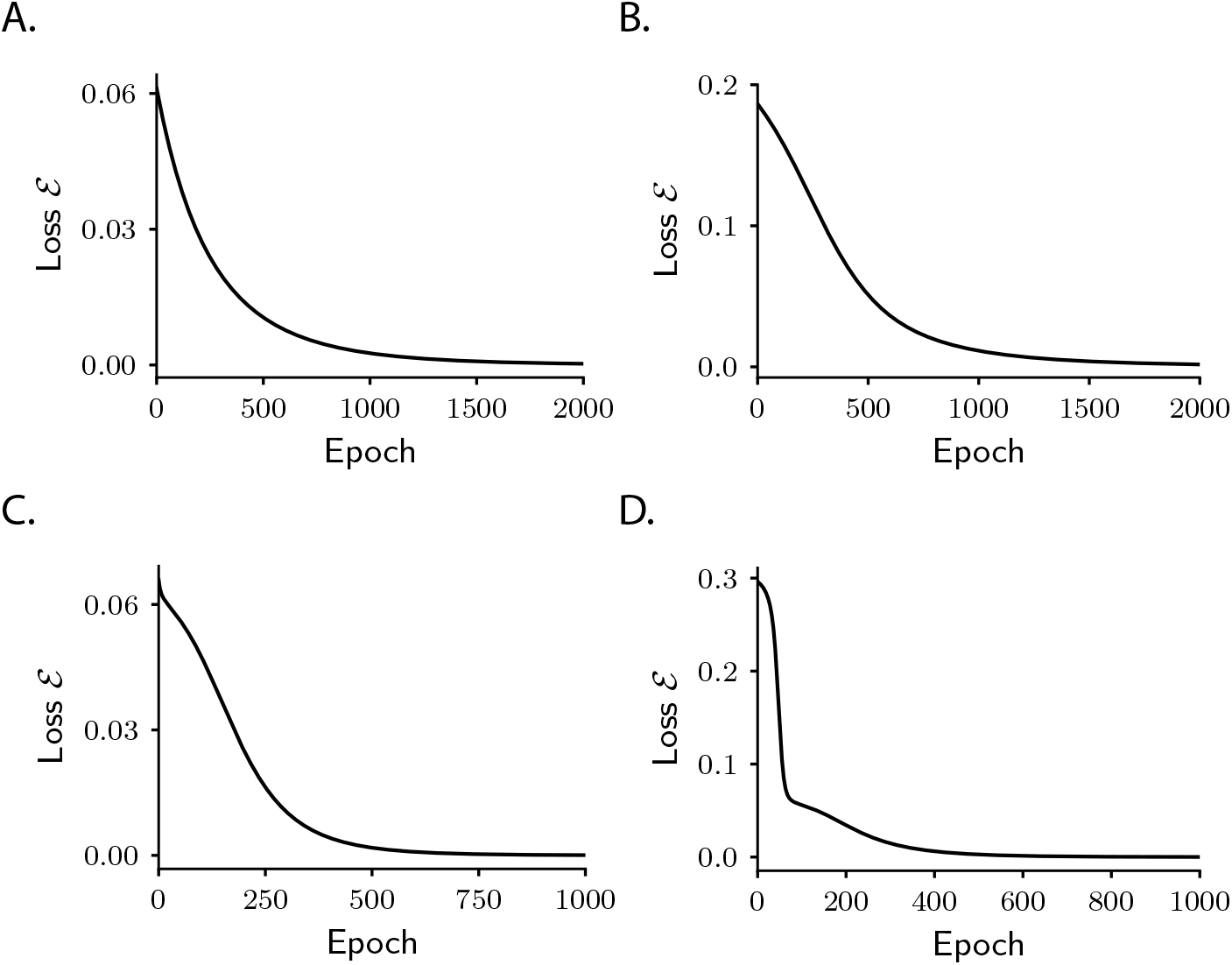
Learning curves. Behaviour of the loss function (Eq. 9) over learning epochs in four sample networks. **A-B** refer to the simple categorization task; parameters are, respectively, as in Fig. 2E-H and Fig. 2I-L. **C-D** refer to the context-dependent categorization task; parameters are, respectively, as in Fig. 6D-F and Fig. 6G-I.

**Fig. S4:**
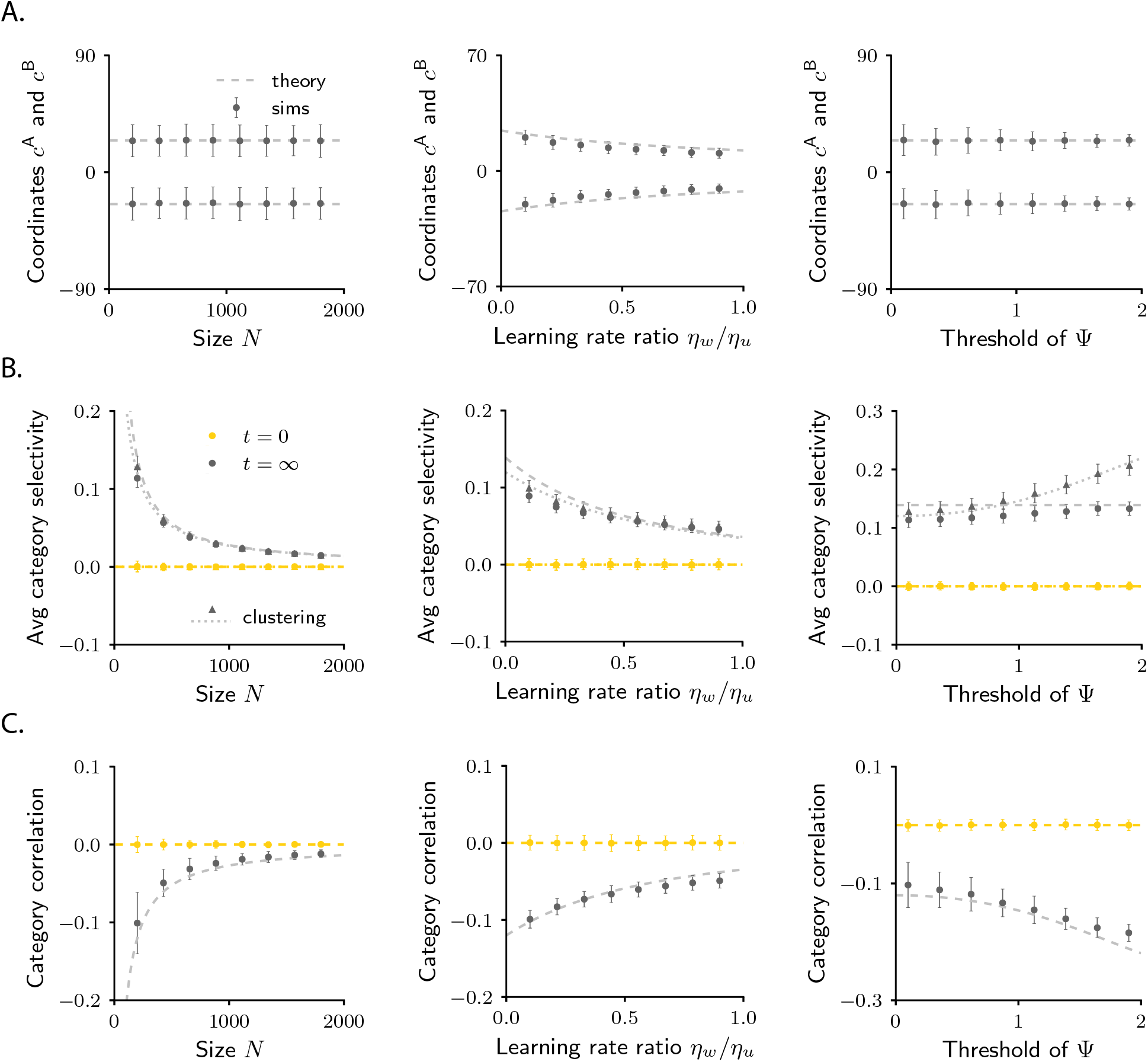
Comparison between finite-size networks and approximate mathematical description for the simple categorization task, part I. Dashed lines show the theoretical predictions; dots show the average over 400 simulations where both the initial connectivity and the sensory inputs were drawn at random. Error bars show the standard deviation across simulations; they thus quantify variability across initializations (Methods 4.7). Yellow: pre-learning; gray: post-learning. **A.** Activity coordinates *c*^A^ and *c*^B^. Theory was computed from Eq. 49. Simulations results were computed by taking the dot product Δ***y**^s^*·***w***_0_ for two activity vectors associated with different categories, and then dividing by 〈Ψ′^2^〉 (Eqs. 55 and 53). **B.** Average category selectivity. Theory was computed from Eq. 66. Simulations results were computed by using Eq. 56, and then averaging across neurons. We also plot results for category clustering. Theory (dotted lines) was computed from Eq. 69. Simulations results (triangles) were computed by using Eq. 67. Note that average selectivity and clustering are not identical for all parameters (although they become close in the limit 1 ≪ *Q* ≪ *N*, as discussed in Methods 4.3). **C.** Category correlation. Theory was computed from Eq. 74. Simulations results were obtained by applying Eq. 71 to simulated data.

**Fig. S5:**
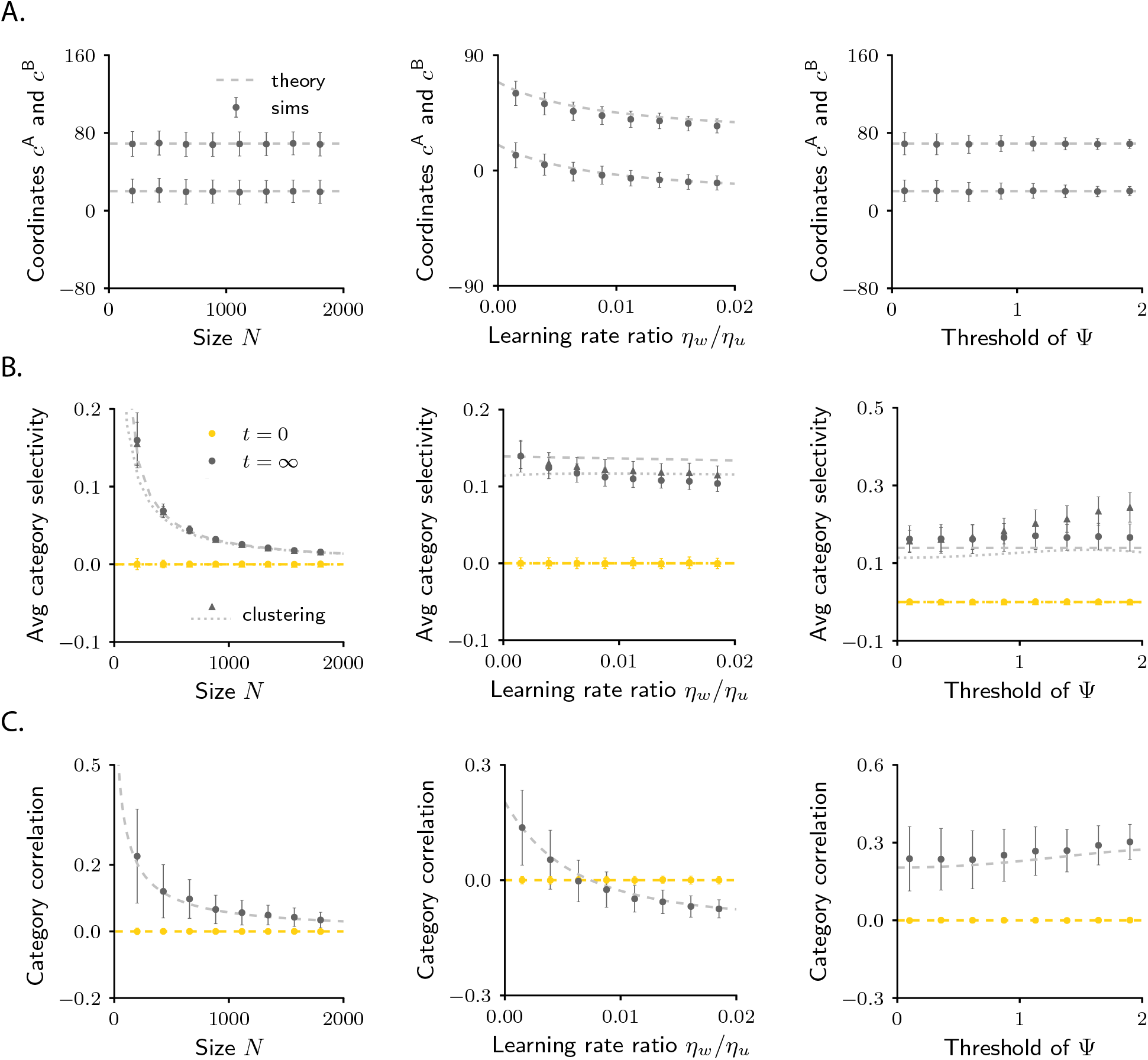
Comparison between finite-size networks and approximate mathematical description for the simple categorization task, part II. Details as in Fig. S4. We used different parameters (see Table 2), which lead to positive category correlation.

**Fig. S6:**
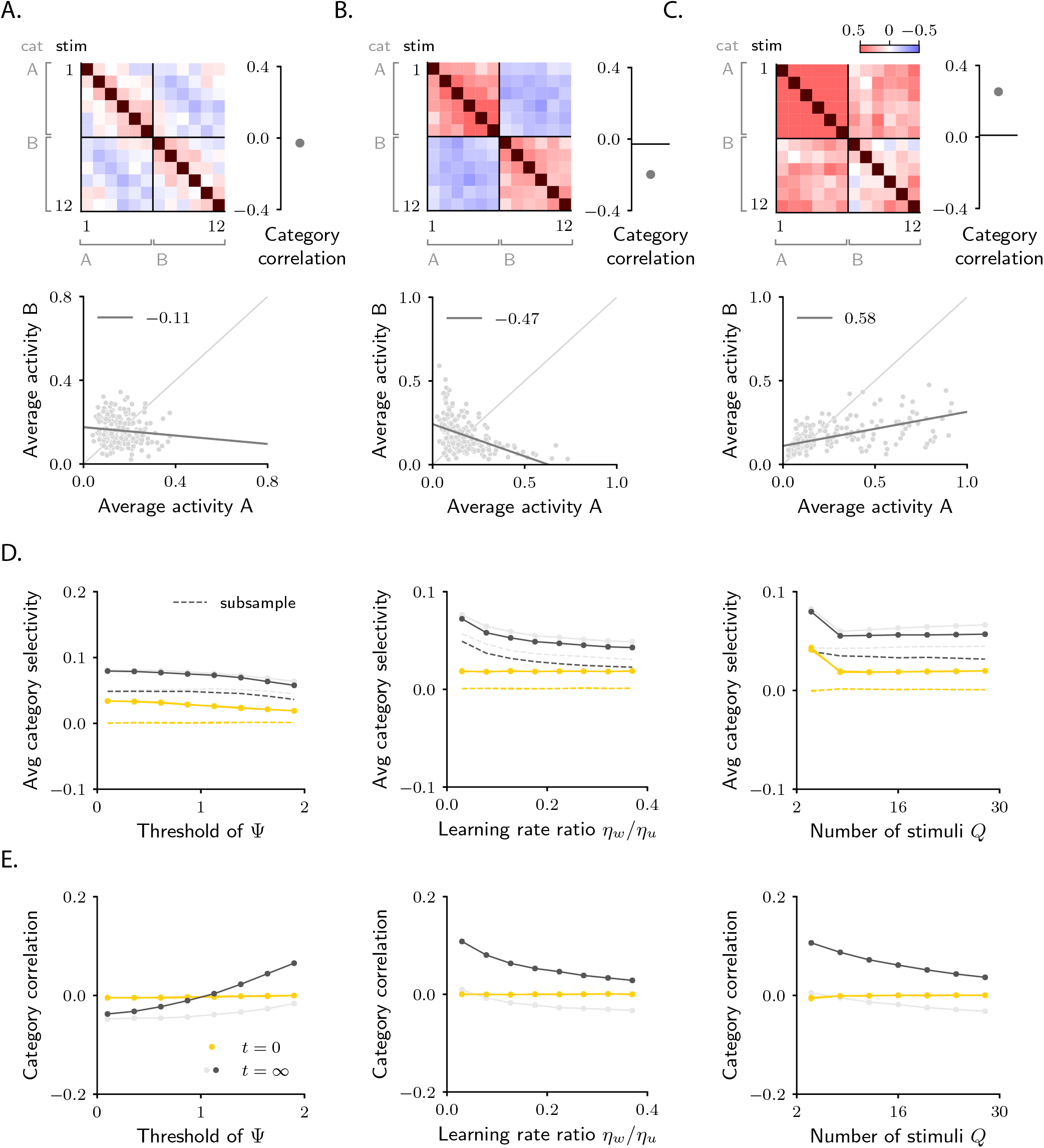
Simple categorization task with structured inputs and heterogeneity (Methods 4.8). **A-C.** Analysis of category correlation for a naive circuit (A), and two trained ones (B-C). Details are, respectively, as in Figs. 2C-D, G-H and K-L. **D.** Extensive analysis of average category selectivity across circuits and task parameters. Details are as in Fig. S4 (except that no theory, only simulations, are shown). As in Figs. 4C-D, dark and light grey indicate different values for the threshold of the activation function Φ. Dashed lines show results obtained by subsampling pairs of sensory inputs in the category selectivity definition (Eq. 56) so that initial selectivity vanishes (see Methods 4.8 for details). **E.** Extensive analysis of category correlation across circuit and task parameters. Details as in panel D.

**Fig. S7:**
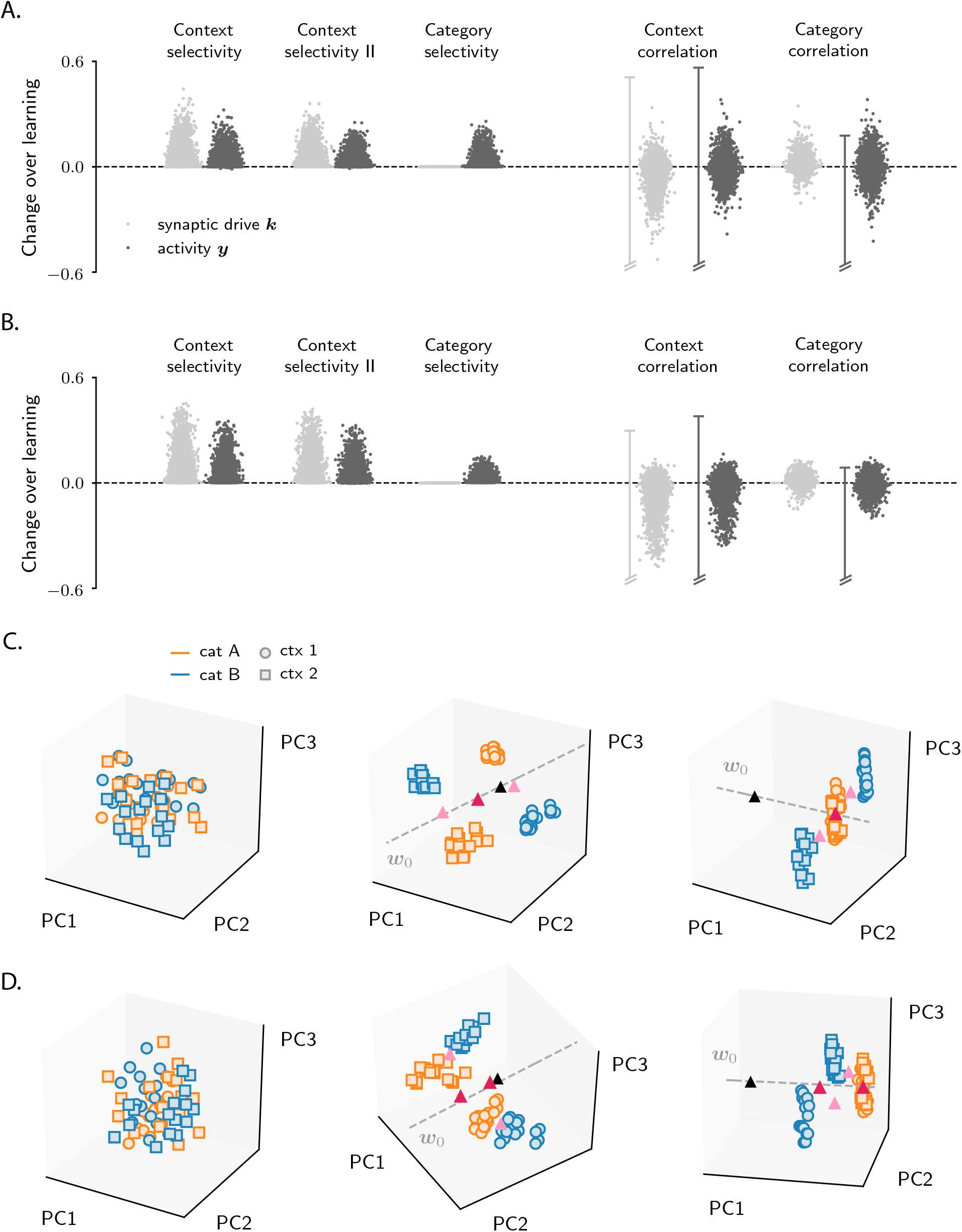
Characterization of activity evolution during the context-dependent categorization task; additional results, part I. A. Extensive results from simulations. Each point represents a model. We simulated 5000 different models, with parameters drawn randomly. Details as in Fig. S1A, except that the number of stimuli and context cues, *Q*, ranges between 4 and 20. To quantify context selectivity, we used Eqs. 122 and 123, and then averaged over neurons. To quantify category selectivity, we used Eq. 56, and then averaged over neurons. To quantify context and category correlation, we used respectively Eqs. 127 and 71. The vertical bars in correlation plots indicate the range of results obtained when quantifying correlation with the alternative definitions in Eqs. 129 and 79; for illustration purposes, the range was cut on the negative semi-axis. For a few parameter sets, learning convergence was pathological (i.e., the loss displayed strong oscillations over epochs, or did not converge); those cases have been excluded from the analysis. Note that, as in Fig. S1A, selectivity changes are positive, while correlation changes take both positive and negative values. **B.** Same as in A, except that changes were computed from theoretical expressions. For context selectivity, we used Eqs. 125 and 126. For category selectivity, we used Eq. 124. For context and category correlation, we used the same equations as in panel A, but evaluated them via the theoretical framework. **C-D.** Activity from circuits in Figs. 6A-C (left), D-F (center) and G-I (right) projected on the first three principal components. Details as in Fig. S1B. Panels C and D display, respectively, results for the synaptic drive ***k*** and the activity ***y***. The center of activity clouds for category `A` and `B` (resp. context `1` and `2`) is indicated by magenta (resp. pink) triangles; the center of initial activity is indicated by the black triangle.

**Fig. S8:**
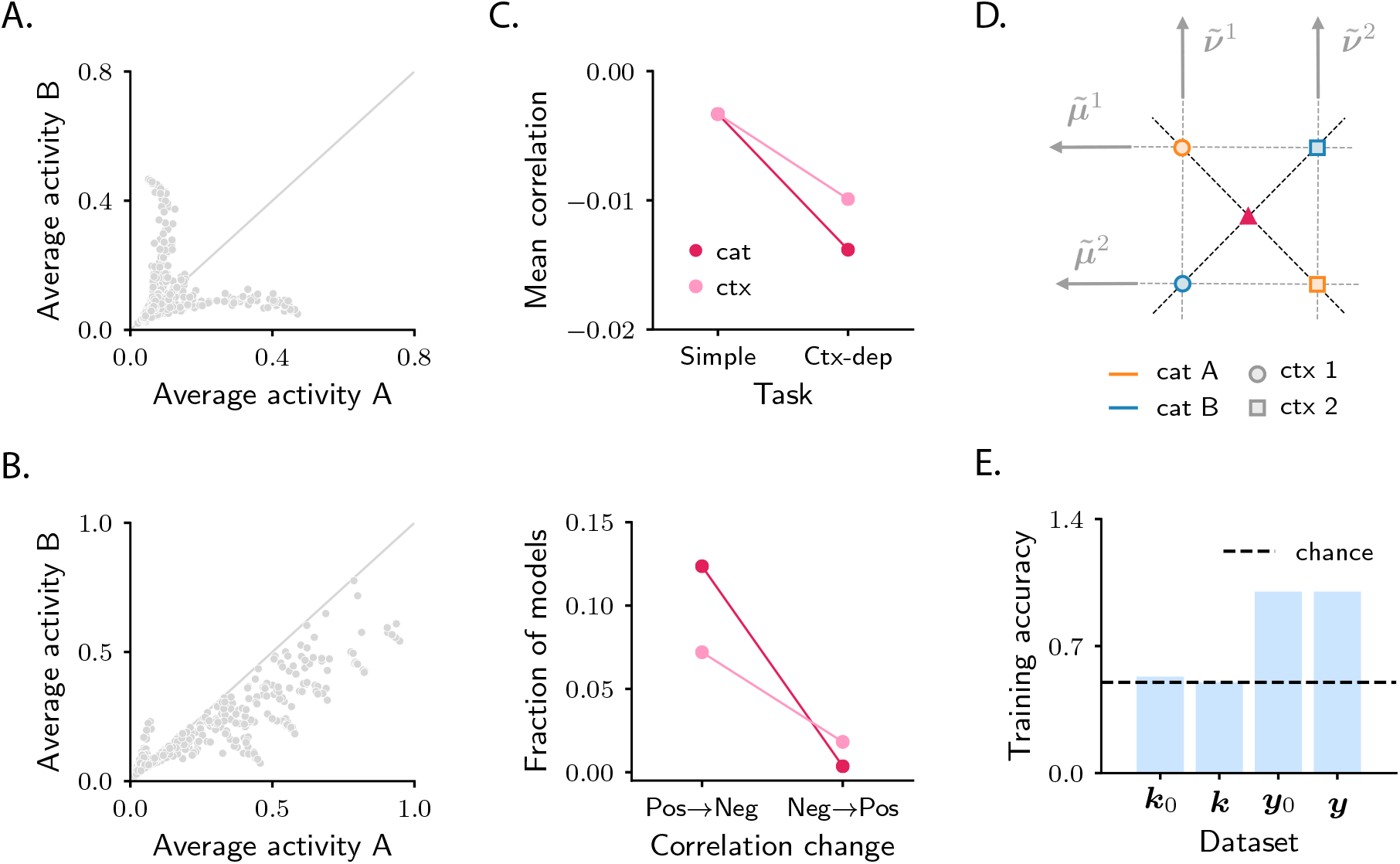
Characterization of activity evolution during the context-dependent categorization task; additional results, part II. A-B. Population response to categories A and B, averaged over trials. Each dot represents a neuron. Panels A and B correspond to two sample circuits, characterized by different values of the threshold of the activation function Φ. Note that responses in panels A are approximately symmetric, while those in panel B are strongly asymmetric. **C.** Context dependence makes categorization more complex and causes a drop in signal correlations: extensive quantification. We computed changes in correlations for the simple and the context-dependent task across a broad range of models; details as in Fig. S7B. The top panel displays the mean of correlation changes across models. Note that correlation changes are more negative in the context-dependent task. For the context-dependent task, we computed both the category and context correlation; results are displayed, respectively, in magenta and pink. In the bottom panel, we computed the fraction of models where correlation changes were positive in the simple task, and became negative in the context-dependent one (left), and viceversa (right). Note that the former substantially outnumbered the latter, which confirms that context-dependency shrinks the parameter space where changes in correlations are positive. **D.** Schematic representation of the initial synaptic drive, 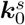, in the contextdependent task with *Q* = 2 (XOR). The centers of activity vectors associated with category `A` and `B` are indicated by magenta triangles; note that the two triangles coincide exactly (Methods 5.8). **E.** Accuracy of a linear classifier trained to decode category from initial and final synaptic drive, ***k***_0_ and ***k***, and initial and final activity, ***y***_0_ and ***y***. Data are from the model displayed in Fig. 6D-F. To train the linear classifier, we used the function `svm.SVC` with a linear kernel from the package `sklearn`; the regularization parameter *C* was fixed through the `GridSearchCV` routine.

**Fig. S9:**
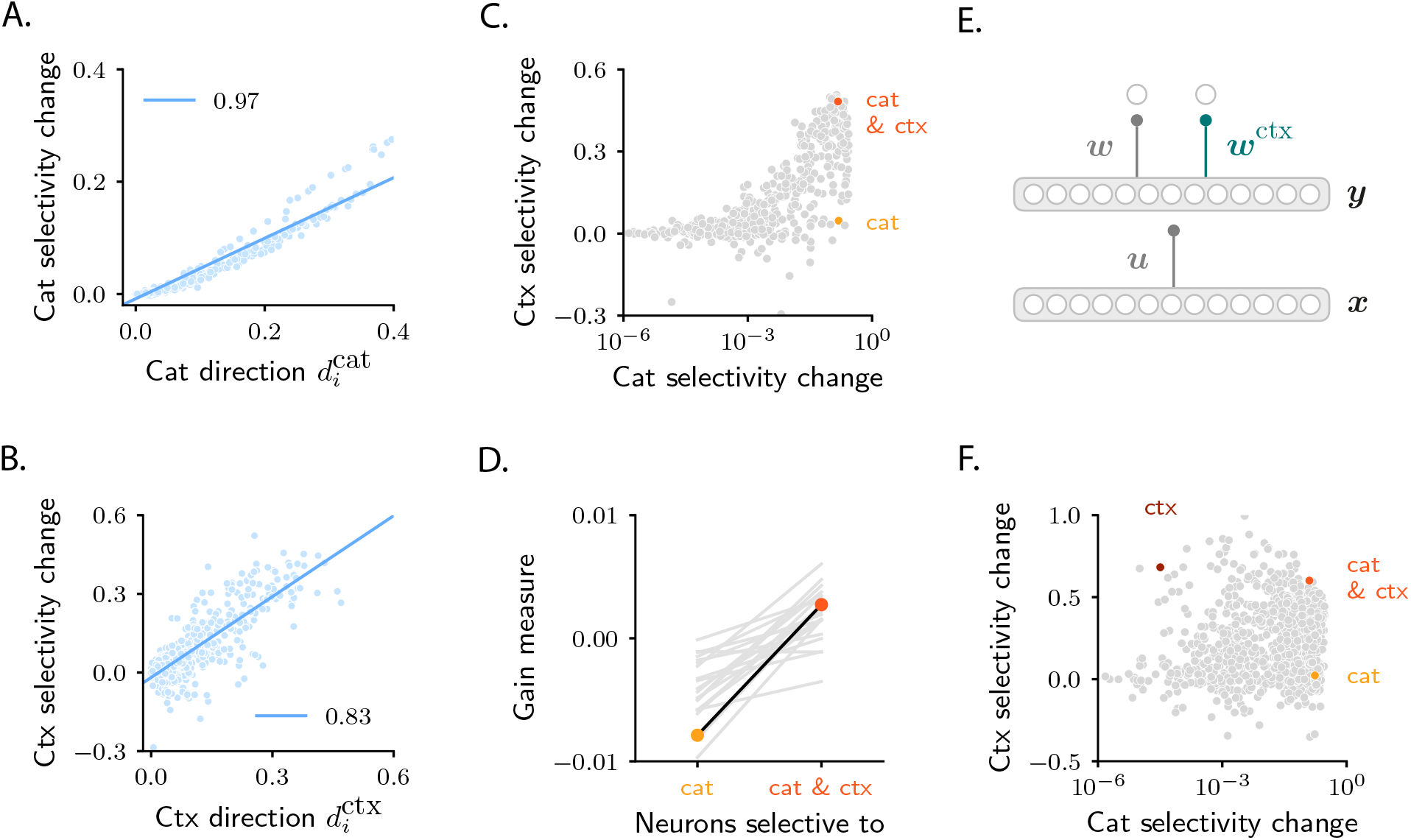
Characterization of activity evolution during the context-dependent categorization task; additional results, part III. A-B. Changes in category (panel A) and context (panel B) selectivity as a function of the components on the category and context directions, 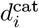 and 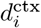 (in absolute value). The latter are defined in Eq. 167, Methods 5.8. Each dot represents a neuron. Data are from the same model displayed in Fig. 8. Blue line: linear fit, with Pearson correlation coefficient shown in the figure legend. Note that correlations for both panels are high. **C-D.** Same analysis as in Figs. 8B-C, but for the sample circuit displayed in the third column of Fig. 6. **E-F.** Analysis of a circuit that includes a second readout neuron, trained to report context. Panel E: circuit architecture; this is identical to the standard model (Fig. 1B), except for the presence of the extra readout. All details and parameters are as in Fig. 8. To make sure that the learning process associated with the two readouts drives activity changes of similar magnitude in the intermediate layer, the target values for the context readout were taken to be *z*^A^ = 0.95, *z*^B^ = 0.05. Panel F: analysis as in Fig. 8B. Note that here, in contrast to Fig. 8B, some neurons display a strong increase in selectivity to context, but not category (brown sample neuron).

**Fig. S10:**
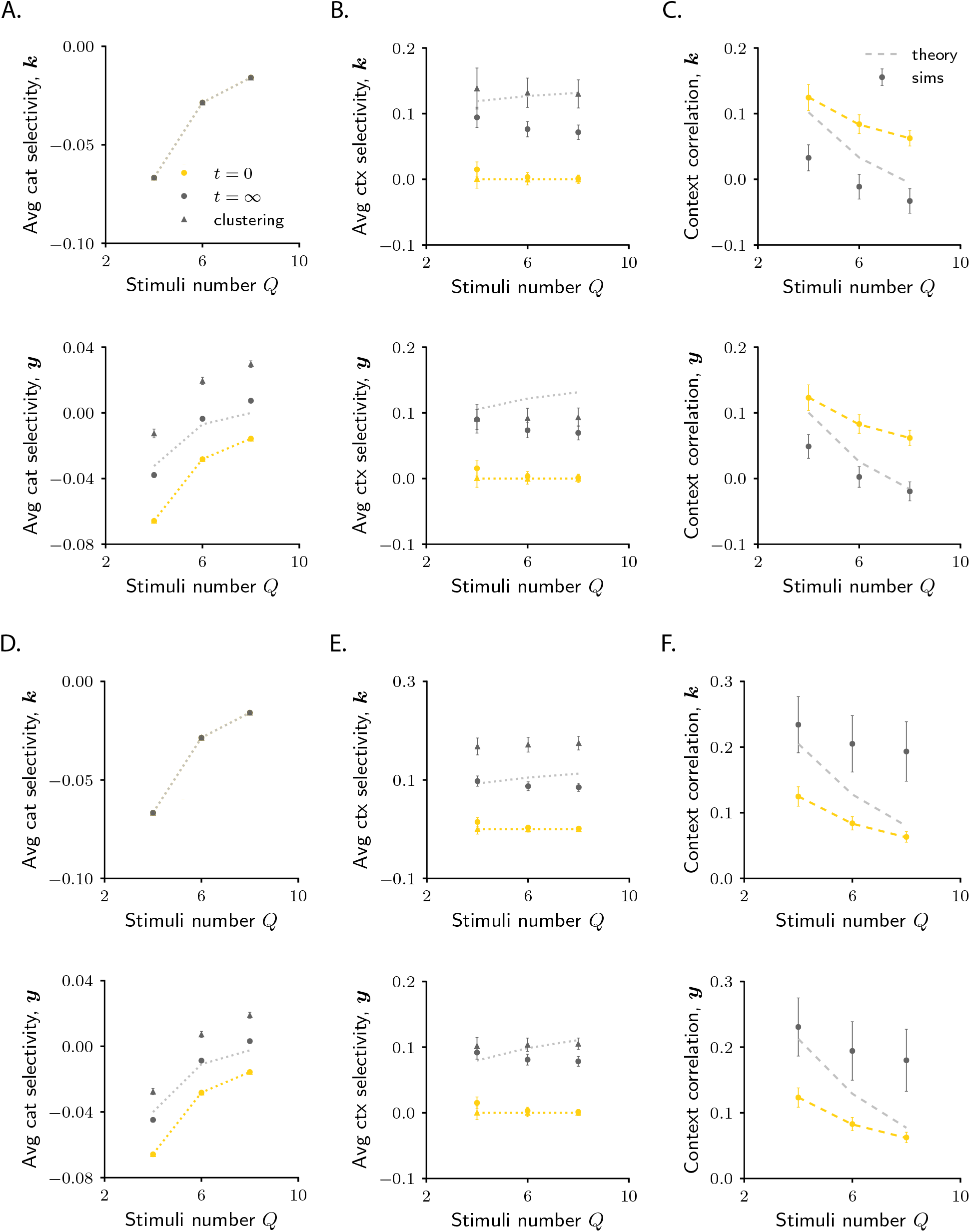
Comparison between finite-size networks and approximate mathematical description for the context-dependent categorization task. Details are as in Fig. S4. In all panels, the top and bottom plots show, respectively, results for the synaptic drive ***k*** and the activity ***y***. **A.** Average category selectivity. Theory was computed from Eq. 124. Simulations results were computed by using Eq. 56, and then averaging across neurons. We also plot results for category clustering (triangles); clustering was computed by applying Eq. 124 to simulated data. **B.** Average context selectivity. Theory was computed from Eq. 125. Simulations results were computed by using Eq. 122, and then averaging across neurons. Context clustering was computed from Eq. 125. **C.** Context correlation. Theory was computed from Eq. 127. Simulations results were obtained applying Eq. 127 to simulated data. **D-E-F.** Details as in A-B-C. We used different parameters (see Table 2), which lead to positive changes in context correlation.

